# Accelerated enzyme engineering by machine-learning guided cell-free expression

**DOI:** 10.1101/2024.07.30.605672

**Authors:** Grant M. Landwehr, Jonathan W. Bogart, Carol Magalhaes, Eric G. Hammarlund, Ashty S. Karim, Michael C. Jewett

**Affiliations:** Department of Chemical and Biological Engineering, Northwestern University; Evanston, IL, 60208, USA; Center for Synthetic Biology, Northwestern University; Evanston, IL, 60208, USA; Department of Bioengineering, Stanford University; Stanford, CA 94305, USA

## Abstract

Enzyme engineering is limited by the challenge of rapidly generating and using large datasets of sequence-function relationships for predictive design. To address this challenge, we developed a machine learning (ML)-guided platform that integrates cell-free DNA assembly, cell-free gene expression, and functional assays to rapidly map fitness landscapes across protein sequence space and optimize enzymes for multiple, distinct chemical reactions. We applied this platform to engineer amide synthetases by evaluating substrate preference for 1,217 enzyme variants in 10,953 unique reactions. We used these data to build augmented ridge regression ML models for predicting amide synthetase variants capable of making 9 small molecule pharmaceuticals. Our ML-guided, cell-free framework promises to accelerate enzyme engineering by enabling iterative exploration of protein sequence space to build specialized biocatalysts in parallel.

## Main

Engineered enzymes are poised to have transformative impacts across applications in energy^1^, materials^2^, and medicine^3^. To create such enzymes, a protein’s amino acid sequence is changed to enhance native function or facilitate new chemical reactions. This process typically involves identifying enzymes with natural plasticity and promiscuity for the reaction of interest, followed by using directed evolution^4,5^. Unfortunately, current approaches to directed evolution are limited because they can only map sequence-function relationships in a narrow region of sequence space. For example, screening strategies are generally low throughput, which constrains re-sampling mutations in iterative site saturation mutagenesis campaigns and can miss epistatic interactions that capture beneficial pairwise (or greater) synergies when the single mutations are neutral or even detrimental^6^. Additionally, selection methods for directed evolution focus on “winning” enzymes for a single transformation, which limits the ability to collect positive and negative sequence-function relationships for forward engineering of similar reactions^7^.

Computational technologies have emerged to accelerate existing directed evolution approaches. *De novo* protein design can create new-to-nature enzymes, but the diversity of chemistries and applications remain limited^8–10^. Machine learning (ML) models have been used to discover enzymes by inferring fitness based on related homologs and/or protein sequences from all organisms (a so-called zero-shot prediction) as well as to navigate protein-fitness landscapes based on assayed fitness data (e.g., nonlinear regression using site-specific one-hot encodings)^11–13^. While ML-assisted enzyme engineering methods show promise, rapidly building datasets to navigate vast sequence space remains a challenge^14^, especially considering most genotype-phenotype links are lost in high-throughput enzyme engineering campaigns^15^.

Here, we developed a high-throughput, ML-guided approach to enable exploration of fitness landscapes across multiple regions of chemical space for forward design of biocatalysts (**Fig. 1**). A key feature of our approach is the use of cell-free gene expression (CFE) systems to allow for the rapid synthesis and functional testing of proteins^16–21^ in a design-build-test-learn (DBTL) workflow. This framework first maps sequence-function relationships for enzyme variants with single-order mutations for a specific chemical transformation identified from an evaluation of enzymatic substrate promiscuity. Then, these data are used to fit supervised ridge regression ML models augmented with an evolutionary zero-shot fitness predictor and extrapolate higher-order mutants with increased activity. Importantly, our ML models can be run on the central processing unit (CPU) of a typical computer making our entire approach user-friendly and accessible.

**Fig. 1:**
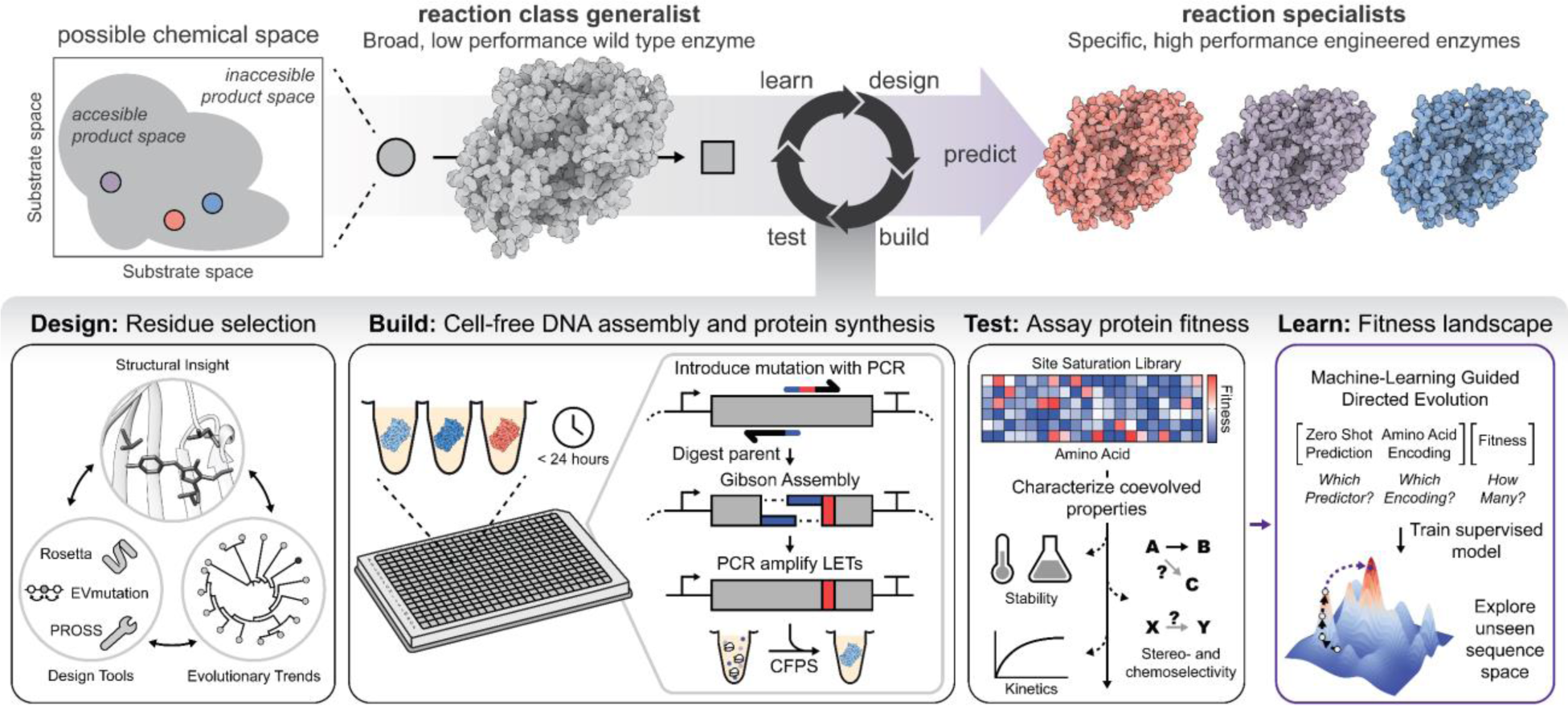
An ML-guided, cell-free enzyme engineering platform. Schematic shows how a design-build-test-learn workflow is applied to rapidly map sequence-function landscapes. Putative residues directing enzyme catalysis are rationally selected based on structural insights, evolutionary trends, and computational tools (e.g., ROSETTA^34^, EVmutation^35^, PROSS^36^) (design). Site saturation mutagenesis and cell-free gene expression are carried out in less than 24 hours to generate sequence-defined libraries (build). The libraries can then be screened for desirable protein fitness metrics (test). Information from the test phase, including failures, is used to identify functionally important amino acid residues that feedback on iterative designs, as well as fit ML models (learn).

We applied our framework to carry out divergent evolution, converting an amide bond-forming generalist enzyme into multiple, distinct specialist enzymes. The biocatalytic formation of amide bonds—a motif ubiquitously found in pharmaceuticals, agrochemicals, polymers, fragrances, flavors, and other high-value products^22^—could offer unique advantages over synthetic counterparts^23–25^ (e.g., mild reaction conditions and chemo-, stereo-, and regioselectivities) and facilitate sustainable biomanufacturing^26–29^. McbA from *Marinactinospora thermotolerans*^30^ is one representative ATP-dependent amide bond synthetase involved in the biosynthesis of marinacarboline secondary metabolites^31^. McbA, and its close homolog ShABS^32^, have been shown to have a relaxed substrate scope, accepting several simple acids and amines commonly found in pharmaceuticals^30,33^. This backdrop suggests that McbA serves as a flexible starting point for engineering a generalist enzyme into multiple reaction specialists each capable of carrying out a different chemical reaction.

## Results

### Exploring the biocatalytic synthesis landscape of McbA

The goal of this work was to develop an ML-guided, DBTL workflow that expedites simultaneous directed evolution campaigns for biocatalysis by reducing screening burden. This goal required generating sequence-fitness data for unique chemical transformations, from which to create predictive ML models. To identify reactions of interest, we first explored the possible amidation reaction space of wild-type McbA (wt-McbA) by evaluating enzymatic substrate promiscuity (**Fig. 2A**). We studied an extensive array of substrates that deviated from the heterocyclic acids and primary or aromatic amines preferred by wt-McbA. These substrates included primary, secondary, alkyl, aromatic, complex pharmacophore, electron poor or rich species, and substrates containing other heteroatoms, halogens, and “unprotected” nucleophiles or electrophiles. More challenging substrates (e.g., complex heterocyclic acids and amines, enantiomers, and substrates containing both acids and amines or multiple acids and amines) were also included to determine the innate limitations and preferences of wt-McbA. We carried out 1,100 unique reactions with low enzyme concentration (∼1 µM) and high substrate concentrations (25 mM), covering 21 molecules of known value including pharmaceuticals, fragrances, and polymers (**Fig. 2B**).

**Fig. 2:**
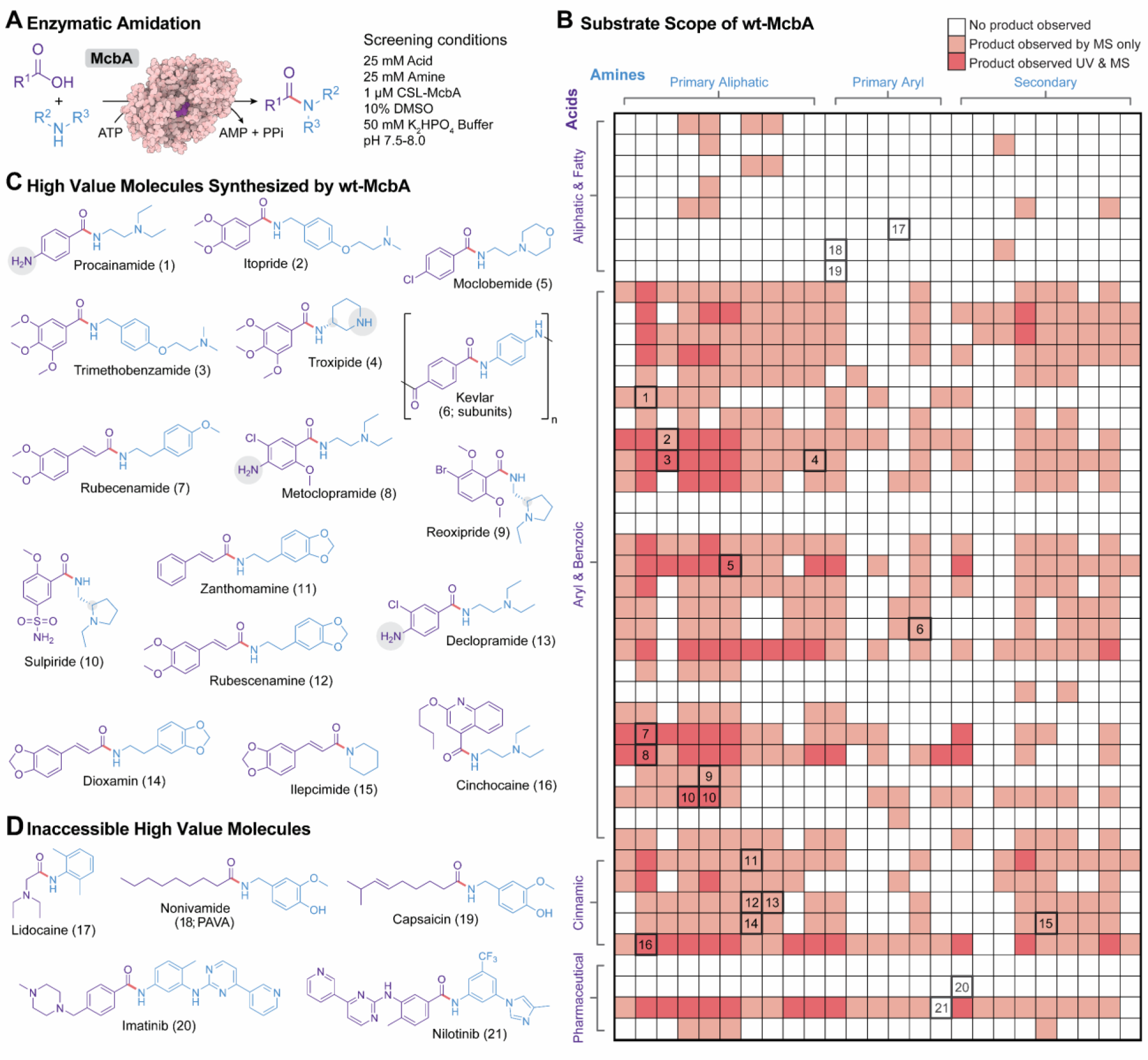
The diverse accessible chemical space of McbA suggests a biocatalyst capable of synthesizing several high value molecules. **(A)** Reaction scheme and screening conditions for exploring the substrate scope of McbA for the enzymatic synthesis of amides. McbA was expressed using CFE and the reaction was initiated by the addition of different combinations of acid and amine substrates. **(B)** The all-by-all substrate screen for McbA, analyzed with RP-HPLC (*n* = 1). Darker red corresponds to a product that was observable by UV absorbance while lighter red corresponds to trace amounts only detectable by MSD. A complete list of substrates can be found in **Fig. S1**. **(C)** Among the 21 high value molecules that were possible in the substrate scope, we observed that McbA was able to synthesize 16 (11 of which are small-molecule pharmaceuticals). **(D)** Example high value molecules that McbA was unable to synthesize under the tested reaction conditions.

Interestingly, wt-McbA displayed a tolerance to multiple “unprotected” functional groups and geometries. Generally, aliphatic acids were poorly tolerated while aryl, benzoic, and cinnamic acids were readily accepted substrates. Charged aryl acids were a unique exception and usually coupled to very few amines. Conversely, wt-McbA readily coupled primary and secondary aliphatic amines but struggled with aryl amines. We observed that McbA was able to synthesize 11 pharmaceutical compounds as well as dozens of hybrid molecules (**Fig. 2C**), ranging from trace amounts detectable only by mass spectrometry (MS) to approximately 12% conversion. In these reactions, we uncovered both stereoselectivity (e.g., strongly favoring the synthesis of S-sulpiride over R-sulpiride) and strict chemo- and regioselectivity preferences (e.g., substrates containing both acids and amines not polymerizing). Given that the reaction mechanism of McbA first begins with the adenylation of the carboxylic acid, we also noticed several instances where only the acyl-AMP intermediate was observed.

### Cell-free protein engineering to rapidly screen sequence-defined protein libraries

With specific chemical transformations identified from our evaluation of enzymatic substrate promiscuity, we next wanted to quickly generate large amounts of sequence-function relationship data of mutant McbA enzymes for training ML models to predict high-activity variants. To do this, we implemented a cell-free protein synthesis approach that does not require laborious transformation and cloning steps (**Fig. 1**). Our approach relied on cell-free DNA-assembly^18^ and CFE^37^ to build site-saturated, sequence-defined protein libraries. This workflow had five steps: (i) a DNA primer containing a mismatch introduces a desired mutation through PCR, (ii) the parent plasmid is digested, (iii) an intramolecular Gibson assembly forms a mutated plasmid, (iv) a second PCR amplifies linear DNA expression templates (LETs), and (v) the mutated protein is expressed through CFE. In this way, hundreds to thousands of sequence-defined protein mutants can be built in individual reactions within a day, and mutations can be accumulated through rapid iterations of the workflow. Our approach avoids any potential biases in typical site-saturation libraries that arise from the use of degenerate primers.

We validated our workflow using the well-characterized, monomeric ultra-stable green fluorescent protein^38^ (muGFP) by targeting four residues that are known to be important for stability and fluorescence^39,40^ (**Fig. S2**). When building our site-saturated library targeting these four residues (77 variants), we found a high tolerance to primer design deviations (e.g., homologous overlaps, melting temperatures) (**Fig. S3 and S4**) and that LETs of muGFP variants conferred all desired mutations (**Fig. S5**). Full-length soluble proteins indicated that changes in fluorescence were not due to changes in expression or solubility (**Fig. S6**). Mapping the protein site-saturated landscape not only highlights residues that are crucial for fitness (e.g., residues composing the fluorophore and impacting hydrophobic core packing were intolerable to mutations^38^) but also provides insight into the general mutability of sites.

After validation, we applied our workflow to McbA to generate sequence-function relationship data that could train ML models to expedite our engineering campaigns. We initially engineered McbA to synthesize three high-value molecules identified by our substrate scope evaluation: (i) the monoamine oxidase A inhibitor, moclobemide, due to McbA’s high promiscuity towards this reaction^33^ (**Fig. 2C** (5); 12% wt conversion); (ii) metoclopramide, due to the unique challenge posed by the acid component containing a free amine that could potentially compete with the intended amine (**Fig. 2C** (8); 3% wt conversion); and (iii) cinchocaine, which contains a unique acid component but shares the same amine fragment as metoclopramide (**Fig. 2C** (16); 2% wt conversion). By performing these engineering campaigns in parallel we hoped to infer mutations that influence substrate specificity for the amine (shared mutations) and the acid (unique mutations) that may lead to general design principles for McbA.

Using relatively high substrate concentrations and low enzyme loading as a step towards more industrially relevant reaction conditions (**Fig. S7**), we performed a hot spot screen (HSS) for each molecule consisting of site-saturation mutagenesis on a wide sequence space to identify residue positions that, when mutated, positively impact fitness (**Fig. 3A**). Guided by the crystal structure of McbA (PDB: 6SQ8), we selected 64 residues that completely enclosed the active site and putative substrate tunnels. Our HSS of these residues (64 residues x 19 amino acids = 1,216 total single mutants) revealed multiple residues that had a positive impact on moclobemide (**Fig. 3B**), metoclopramide (**Fig. 3C**), and cinchocaine (**Fig. 3D**) synthesis when mutated compared to wt-McbA as measured by liquid chromatography-mass spectrometry (LC-MS).

**Fig. 3:**
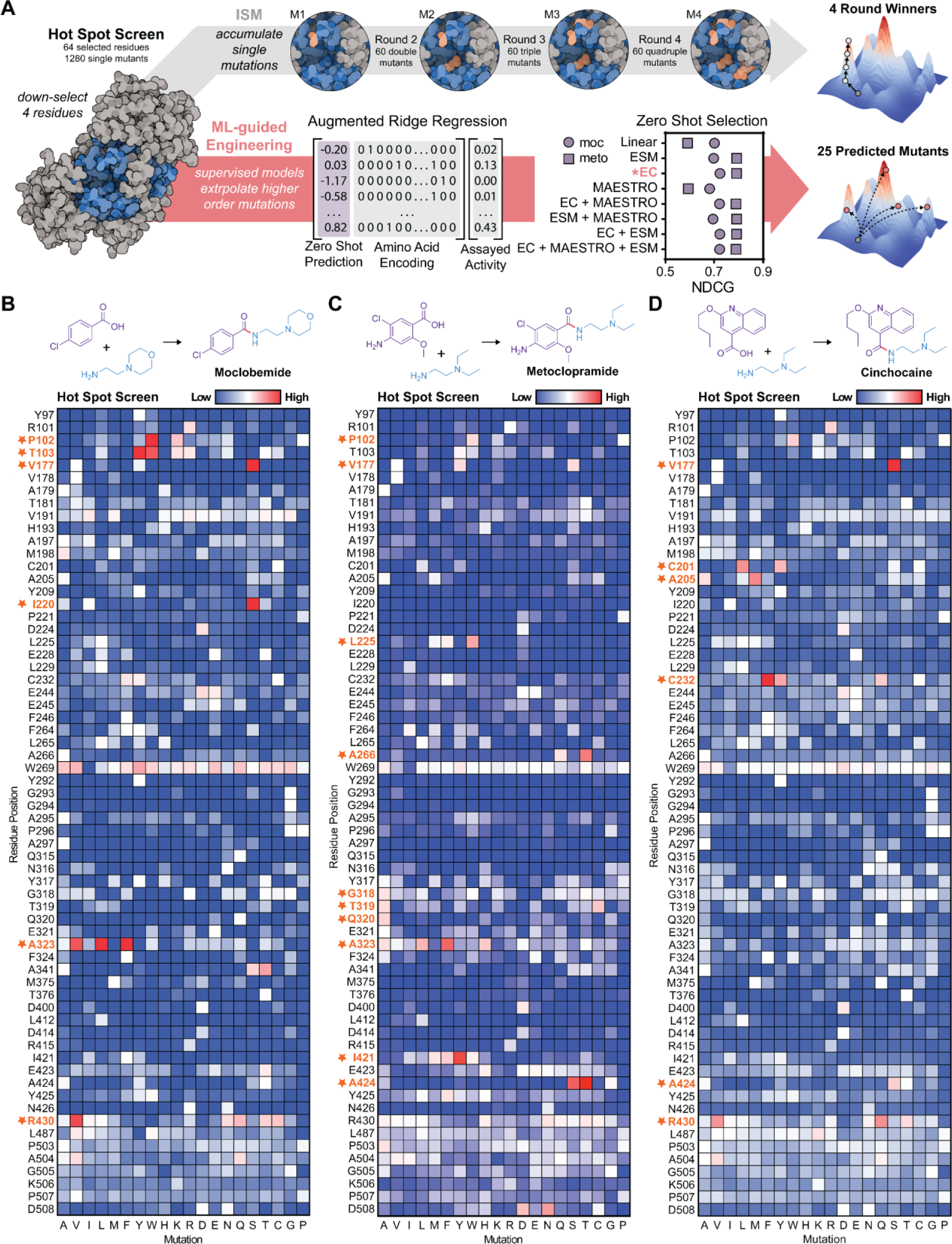
Rapid generation of sequence-fitness landscape data for ML-guided directed evolution of McbA. **(A)** Workflow schematic: a supervised ridge regression model is trained on percent conversion data of four mutable sites selected from the HSS, with sequence features consisting of an amino acid encoding augmented with a zero-shot prediction of enzyme fitness. From a training set of approximately 80 single mutants, we extrapolate higher order mutations and test the top 25 predictions. **(B-D)** Hot spot screen of 64 identified residues in McbA showing percent conversion of (**B**) moclobemide, (**C**) metoclopramide, and (**D**) cinchocaine normalized to WT (*n* = 1). Highly mutable sites, or hot spots, are highlighted.

### ML-guided, cell-free expression for protein engineering

With a large data set at hand for multiple, distinct single McbA mutants, we set out to leverage ML models to accelerate the engineering of McbA for the production of small molecules across diverse regions of chemical space. We sought to use single mutant data derived from the HSS to fit augmented ridge regression models—given their power and simplicity for protein engineering^41^—allowing us to predict higher-order mutants with increased activity (**Fig. 3A**).

We first selected a predictive model architecture. McbA variant feature representations consisted of site-specific amino acid encodings concatenated with a zero-shot fitness prediction^41^. We considered several amino acid encodings, ranging from simple one-hot encodings to more complex descriptors that attempt to incorporate amino acid physiochemical properties^42–45^. We also explored benchmark protein variant fitness predictors to incorporate universal, evolutionary, and structural based zero-shot predictions. We tested three specific fitness predictors: the Evolutionary Scale Modeling (ESM)-1b transformer^46^ trained on the UniRef50 database (universal), an EVmutation (EC)^35^ probability density model trained on an MSA of evolutionarily related sequences (evolutionary), and MAESTRO^47^ to estimate structure-based changes in unfolding free energy (structural). Training and hyperparameter tuning were performed using single mutant data (*n* = 77) from the HSS (top four hot spots; **Fig. 3B** and **Fig. 3C**).

In parallel, we conducted a more traditional directed evolution campaign on each amide product (moclobemide, metoclopramide, and cinchocaine) via iterative saturation mutagenesis (ISM). This would provide valuable higher order mutations to validate and benchmark model performance given our objective of extrapolating from single to higher order mutations.

For moclobemide, we selected six residues from the HSS to mutate over three rounds of ISM (**Fig. S7**). We first fixed the top mutation from the HSS (V177S) and performed site-saturation mutagenesis on the five additional residues. By reintroducing previously fixed mutations in subsequent rounds, we explored potential epistatic interactions (e.g., S177 was saturated in ISM round 2, given V177S was incorporated before A323F). In addition, we completely explored all combinatorial double mutants of the top two residues, which showed directly additive impacts for moclobemide synthesis (**Fig. S7**). After three rounds of ISM, we identified a quadruple mutant (qm-McbA_moc_) with increased activity—from 12% for wt-McbA to 96% conversion—for the synthesis of moclobemide (**Fig. S7**). We characterized the apparent steady-state kinetic parameters and stability of these enzyme mutants from each round of ISM. Specifically, we expressed, purified, and evaluated each McbA variant observing a 42-fold increase in catalytic efficiency from wt-McbA to qm-Mcba_moc_ (k_cat_/K_M_ increased from 18.2 to 769 M^-1^min^-1^) for the amine (**Fig. S8** and **S9**). The melting point did not significantly change between wt-McbA and qm-McbA_moc_, but the second mutation (A323F) increased T_m_ by 5.81 ± 0.09 °C when added to the first mutation (V177S) (**Fig. S10**). Additionally, we showed that we could make milligram quantities of moclobemide in a 10-mL reaction (87% isolated yield) and confirmed the structure by NMR (**Fig. S11** and **Fig. S12**).

Three rounds of ISM for metoclopramide yielded a quadruple mutant that displayed nearly 30-fold activity over wt-McbA (**Fig. S13**). The campaign for cinchocaine was more difficult to navigate and we failed to observe beneficial mutations beyond a double mutant, despite taking multiple ISM paths (**Fig. S14**). This result (i.e., running into dead ends during ISM) supported the need to include ML models in our framework that might capture epistatic interactions. We used the ISM data for moclobemide and metoclopramide containing double, triple, and quadruple mutants (*n* = 243 for moclobemide and *n* = 169 for metoclopramide) to evaluate each model’s performance, while cinchocaine would provide a unique pressure-test for our identified top-performing model.

Model prediction performance was first evaluated using the normalized discounted cumulative gain (NDCG)^14,48^, an evaluation metric that scores models on their ability to correctly rank high-fitness variants (aligning with our experimental goal of discovering high-fitness variants with minimal screening burden), which generally matched results from the Spearman rank correlation coefficient (**Fig. S15**). The evaluated augmented models outperformed the ridge regression model alone. We also tried combining predictors in our variant features (e.g., predictions from both ESM-1b and EVmutation), but no increase in model performance was observed. Lastly, we tested the necessity of the entire site saturation dataset (*n* = 77) for training models to achieve high predictive performance. We withheld variants in the training set to reflect common protein engineering strategies that do not exhaustively search the sequence space, including reduced codon libraries (NDT^49^ and NRT^50^), single amino acid scans^51^ (here, we combine the commonly used glycine, alanine, proline, and cysteine scans), and reduced alphabets that naturally group amino acids by physiochemical properties (BLOSUM^52^). When training the same augmented ridge regression model with Georgiev encodings, this analysis indicated that utilizing all the data gathered from the site saturation dataset provides more predictive power (**Fig. 4A-B**). This can likely be attributed to the nature of the rich datasets mostly containing mutants with non-zero activity (64/77 for moclobemide and 62/77 for metoclopramide), preventing “holes” in training sets^14^. Moving forward, we decided to use the site saturation dataset and the augmented EVmutation model with Georgiev encodings given the strong predictive performance among both compounds and the fact that the already-trained probability density model simplified application to other compounds. EVmutation is also less computationally resource and time intensive than ESM.

**Fig. 4:**
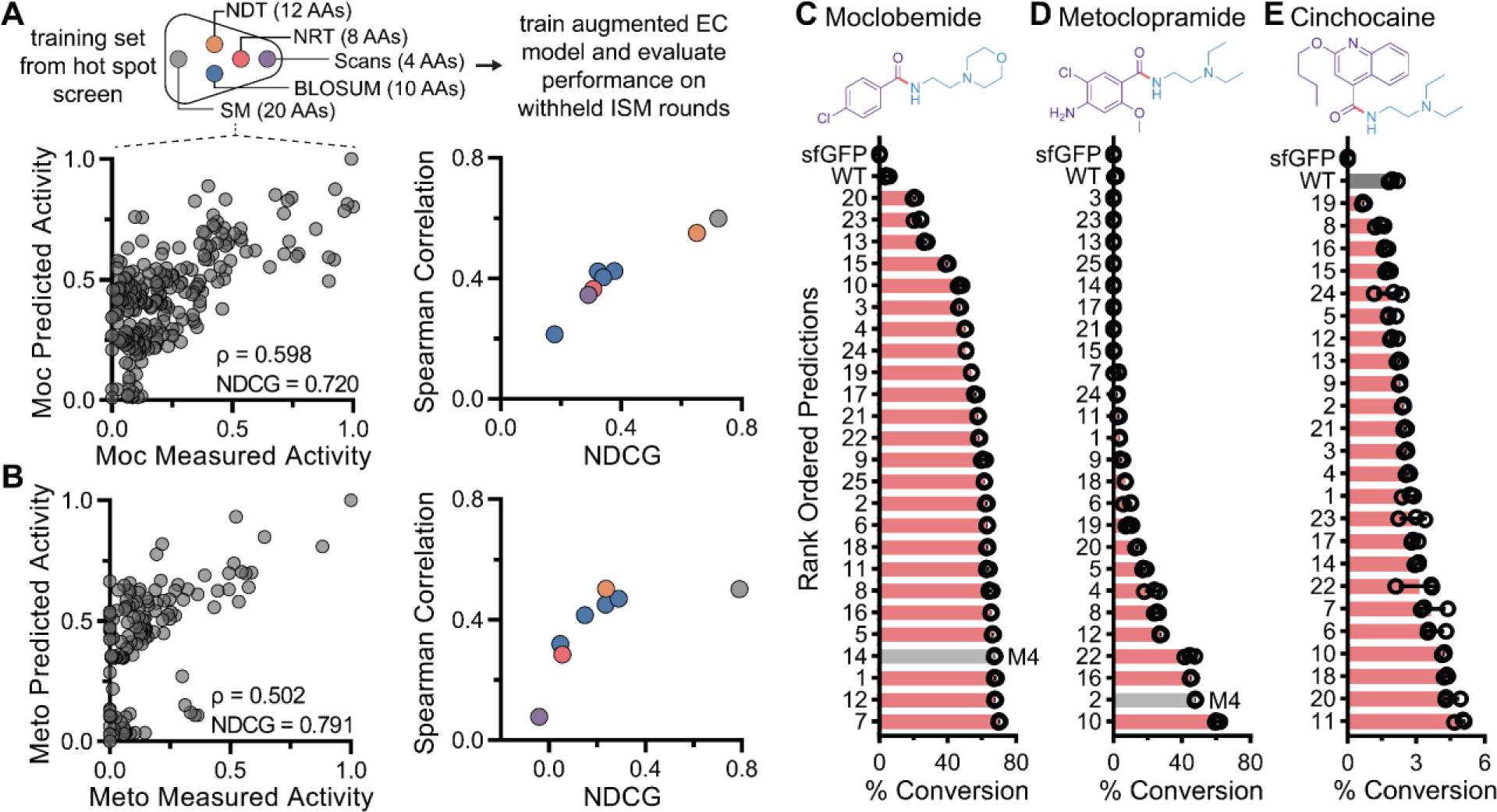
ML-guided directed evolution predicts highly active mutants with a lower screening burden than iterative site saturation mutagenesis. **(A,B)** Analysis of model fidelity with training sets built with smaller libraries than saturation mutagenesis, including reduced codon sets (NDT, NRT) and reduced amino acid alphabets based on BLOSUM50. Comparing measured versus predicted activity on withheld ISM rounds is shown for models trained on the complete saturation mutagenesis dataset for both moclobemide (moc) (**A**), and metoclopramide (meto) (**B**). **(C-E)** The experimentally validated percent conversion (*n* = 3) of ML-predictions for moclobemide (**C**), metoclopramide (**D**), and cinchocaine (**E**) with the quadruple mutant from ISM (M4) colored grey. For cinchocaine, the ML model predictions did not include the highest performing mutant from ISM. Wild type McbA is colored dark gray.

Using our trained ML models, we screened 20^4^ combinatorial enzyme variants for the synthesis of moclobemide, metoclopramide, and cinchocaine *in silico* and selected the top 25 predictions to subsequently build and test. We found that the augmented ML model was able to predict McbA variants enriched in high activity for each amide product when tested experimentally, some even surpassing qm-McbA from both moclobemide and metoclopramide ISM campaigns (**Fig. 4C-E** and **Table S1**). Notably, the best predicted mutant for metoclopramide contained a mutation (A424S) that was superseded in the HSS by a more active mutation (A424T) carried forward in ISM, indicating the model found a superior mutant that would have been overlooked using ISM alone. The best predicted variant for cinchocaine had significantly higher activity than the best single mutation on its own and surprisingly contained a mutation (A205L) that decreased activity compared to wt-McbA in the HSS (**Fig. S16** and **Fig. S17**); we could not rationally select and combine mutations from the HSS to reach the same results. These results show that our ML-guided strategy can discover high fitness variants for a variety of molecules using the same starting enzyme while avoiding path dependencies and reducing the screening burden.

### ML-guided biocatalytic diversification for high-value pharmaceuticals

We next applied our ML-guided framework to predict distinct McbA mutants for the synthesis of an additional six pharmaceutical compounds. Starting with an identified target reaction from our substrate scope screen (**Fig. 2**), we used the same instance of our 1,216 single mutant McbA variant library from above to perform an HSS (7,302 unique reactions total), select four hot spots, and train our ML model to predict higher order mutants with increased activity (**Fig. 5A**; **Fig. S18-23**). For each reaction, the top 24 predictions were tested, and the best variant was expressed, purified, and compared to wt-McbA activity. We observed increases in yields ranging from 1.6-fold to 34-fold over wt-McbA for the six compounds we tested (**Fig. 5B-G** and **Fig. S18-23**). For each compound, the best predicted mutant always outperformed the best rational design (i.e., combining the four best mutations from the HSS without using the ML model; **Table S3-5**). Some mutants give only modest improvements, which may be an artifact of low signal-to-noise in the hot spot screens for some of the target compounds that were only detectable by MS. This can lead to flat fitness landscapes that are more difficult to model. Nonetheless, our framework yielded enzyme mutants with increased activity for multiple products that were initially only observed in trace amounts.

**Fig. 5:**
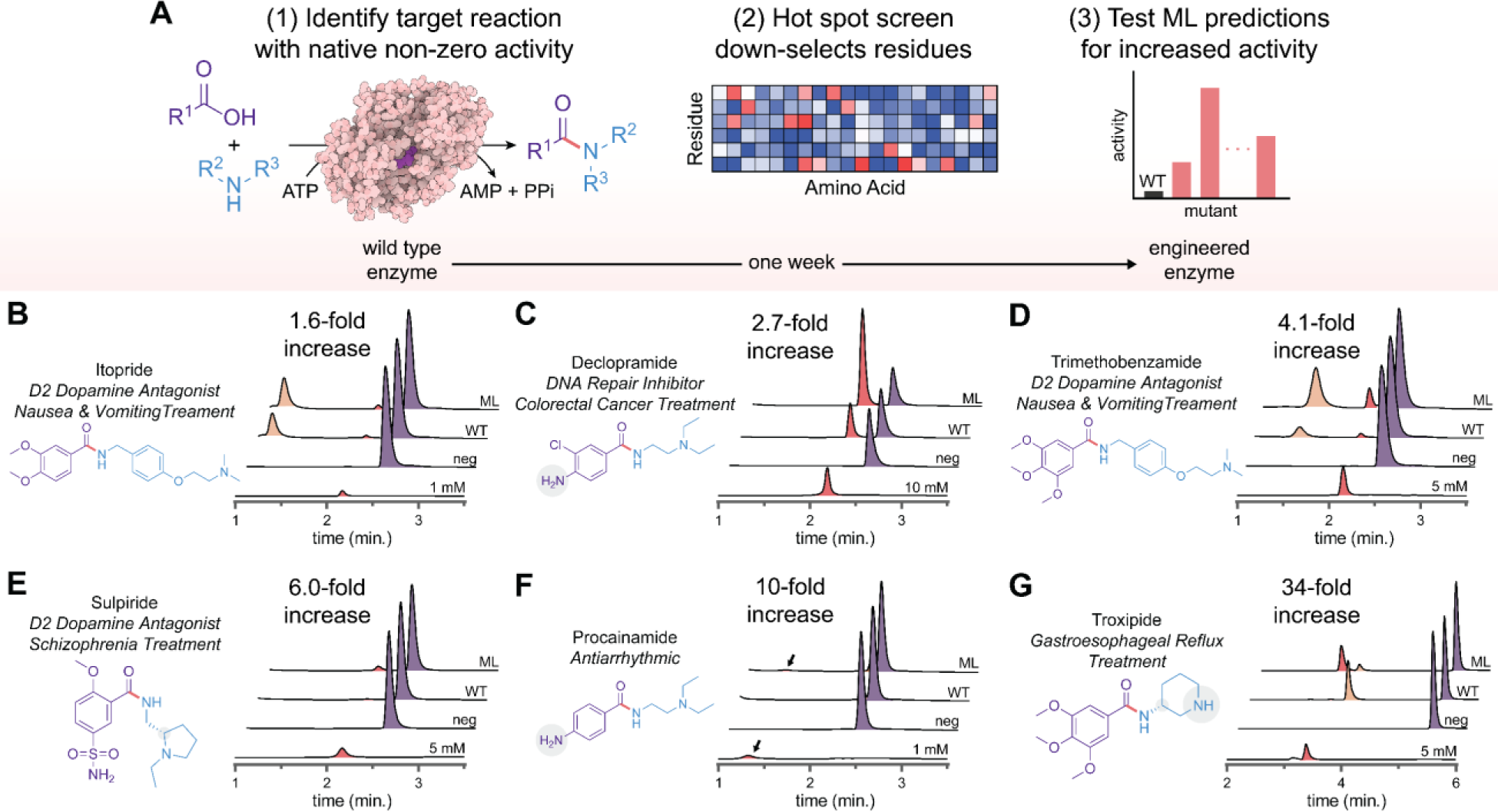
ML-guided engineering of distinct amide synthetases for the biosynthesis of a broad panel of small-molecule pharmaceuticals. **(A)** The strategy we utilized for the machine learning-guided protein engineering of McbA. First, we identified non-native reactions that wt-McbA can catalyze and prioritized those that produce valuable small-molecule pharmaceuticals. Second, an HSS of 64 residues is used to down-select residues that positively impact activity. Third, an augmented ridge regression model is trained on data from the HSS, and ML predictions are experimentally tested. **(B-G)** Comparison of the highest activity predicted variant for a panel of small-molecule pharmaceuticals compared to wt-McbA and an authentic standard. Enzyme concentration was normalized to 0.5 mg/mL (approximately 9 µM) and products were analyzed by (RP)-HPLC. The fold-increase in yield observed compares wt-McbA to ML-McbA (*n* = 3). Representative HPLC traces of product (red), acid substrate (purple), and adenylated acid (orange) for each reaction are shown.

We also compared how efficiently some enzyme variants perform each reaction step (**Fig. S24**). For example, wt-McbA appears to be proficient at the adenylation step for troxipide (adenylating 3,4,5-trimethoxybenzoic acid), but unable to catalyze amide bond formation (**Fig. 5G**). The engineered enzyme variant can subsequently accept the amine, leading to a large decrease in the observed intermediate. Serendipitously, the engineered McbA variants for each target product also displayed strict regioselectivity despite the lack of any selective pressure to maintain it. This is exemplified by the quadruple mutant for troxipide that exhibits a 34-fold increase in activity without any sacrifice in specificity. Similarly, stereoselective preferences with S-sulpiride are maintained. Taken together, our ML-guided framework allows us to use functional data from single mutant enzyme variants to predict superior higher-order mutants rapidly and effectively.

## Discussion

In this work, we establish a high-throughput, ML-guided protein engineering framework for predictive design that does not require specialized computational resources. This framework uniquely integrates a cell-free gene expression and mutagenesis method, ML to expedite directed evolution campaigns, and divergent evolution to convert a generalist enzyme into multiple specialists. We showcased this framework by rapidly navigating nine protein engineering campaigns for the amide synthetase McbA, six of which were performed simultaneously. Through efforts to build ML models and all ISM rounds, we comprehensively mapped the sequence-function landscape of McbA by assessing 2,856 variants of McbA, 1,100 possible amide products, and 12,584 substrate pair-mutant reactions. We identified 19 unique residue positions within McbA that significantly impact biocatalysis. Across all nine engineered McbA variants, we made a total of 21 different mutations occurring across 14 different residues (**Fig. S25**). In all cases, newly generated enzyme variants demonstrated improved activity relative to wt-McbA variants (1.6-fold to 42-fold improvement, including moclobemide). In one example, an enzyme variant for moclobemide synthesis achieved 96% conversion (a 42-fold increase in catalytic efficiency over wt-McbA) and was scaled to milligram quantities.

An important feature of our work was the use of ML models trained on single-residue mutations to predict higher order mutants with improved fitness. Strikingly, we observed that ML-predicted enzyme variants with 4 mutations had greater activity than the combination of the four most active single-residue mutants alone in each of the nine test cases. In other words, our data suggests that one may be able to screen N×20 mutants instead of 20^N in a directed evolution campaign.

Our approach can, in theory, be applied to any enzyme but may require reaction-specific fine-tuning around data collection and ML model generation. In terms of data collection, experimental screening methods for biocatalytic reactions remain a bottleneck. Here, because the product compounds of McbA were stable in the presence of the cell-free expression lysate and chromatographic methods were efficient (e.g., ∼3 min/per sample), liquid chromatography-mass spectrometry provided a manageable solution, as has been found in other examples^53^. As a complement to screening, there will be enzyme engineering applications where selection strategies are beneficial (e.g., when a tractable selection method exists, larger jumps in sequence space can be made). Engineering campaigns with different proteins may also warrant exploring various ML models and parameters. While we saw excellent performance with the augmented EVmutation model, alternative fitness goals may also require alternative fitness predictors. For example, if the goal is to engineer stability, it would reason that a structural-based fitness predictor may be superior. There are numerous additional protein variant effect predictors continuously pushing the state-of-the-art forward that could improve our predictions^54^. More complex models based on natural language processing may also outperform linear regression^46,55^. Finally, we note that training the ML models on more residues, multi-mutant data (including data from multiple rounds of mutagenesis or random combinations of mutants that diversify amino acids across the entire protein of interest), or kinetic measurements could be beneficial in engineering better catalysts.

In sum, our accessible ML-guided, cell-free framework overcomes traditional directed evolution challenges by circumventing path dependencies that constrain sequence search space in state-of-the-art methods. This speeds the pace of engineering relative to ISM alone. Our work further highlights the versatility of the amide synthetase McbA to be directed to catalyze many unique reactions of interest, including those used in small-molecule pharmaceutical production. Looking forward, we anticipate that the approach described here, especially when augmented with *de novo* protein design, will accelerate enzyme engineering campaigns to unlock specialized enzymes with diverse functions and properties.

## Methods

### Cell-free DNA assembly and gene expression

DNA libraries were created for both wt-McbA and muGFP. wt-McbA from *Marinactinospora thermotolerans* (UniProt: R4R1U5) was codon-optimized for *E. coli* and cloned into the pJL1 plasmid (Addgene, 69496) with an N-terminal CSL-tag^56^ (CAT-Strep-Linker fusion containing Strep-tag II). muGFP was codon-optimized for *E. coli* and cloned into the pJL1 plasmid without a purification tag^38^.

The cell-free DNA library generation was performed as follows: (1) the first PCR was performed in a 10-µL reaction with 1 ng of plasmid template added, (2) 1 µL of DpnI was added and incubated at 37°C for two hours, (3) the PCR was diluted 1:4 by the addition of 29 µL of nuclease-free (NF) water, (4) 1 µL of diluted DNA was added to a 3-µL Gibson assembly reaction (self-made)^57^ and incubated for 50 °C for one hour, (5) the assembly reaction was diluted 1:10 by the addition of 36 µL of NF water, (6) 1 µL of the diluted assembly reaction was added to a 9-µL PCR reaction. All cloning steps were set up using an Integra VIAFLO liquid handling robot in 384-well PCR plates (Bio-Rad). Primers were designed using Benchling with melting temperature calculated by the default SantaLucia 1998 algorithm. We have noticed that melting temperatures of alternative primer design tools sometimes deviate from those calculated in Benchling, so users should consider this when designing primers. The general heuristics we followed for primer design were a reverse primer of 58 °C, a forward primer of 62 °C, and a homologous overlap of approximately 45 °C. All primers were ordered from Integrated DNA Technologies (IDT); forward primers were synthesized and received in 384-well plates and normalized to 2 µM for ease of setting up reactions. Additional information on primer design and the codons we used for all 20 amino acids can be found in **Fig. S4** and **Table S7**. All PCR reactions used Q5 Hot Start DNA Polymerase (NEB). Additional information on thermocycler parameters can be found in **Table S8**.

To accumulate mutations for ISM, 3 µL of the “winner” from the diluted Gibson assembly plate was transformed into 20 µL of chemically competent *E. coli* (NEB 5-alpha cells). Cells were plated onto LB plates containing 50 µg/mL kanamycin (LB-Kan). A single colony was used to inoculate a 50 mL overnight culture of LB-Kan, grown at 37 °C with 250 RPM shaking. The plasmid was purified using ZymoPURE II Midiprep kits and sequence confirmed. Successive mutations can then be incorporated via our cell-free DNA library generation method above.

The comprehensive combinatorial double mutant McbA library (used in **Fig. S7e**) was generated by two successive rounds of saturation mutagenesis with no particular residue targeted first. After the first site saturation, plasmids containing each mutation were prepared following the above protocol except 5 mL LB-Kan overnights were used to purify plasmids using ZymoPURE II Miniprep kits. These 20 plasmids were used as templates for the next round of site saturation mutagenesis to accumulate all 400 double mutants.

All machine learning predicted McbA variants were ordered as gblocks from IDT containing pJL1 5’ and 3’ Gibson assembly overhangs. DNA was resuspended at a concentration of 25 ng/µL. A linearized pJL1 plasmid backbone was ordered as a gblock from IDT, PCR amplified, purified using a DNA Clean and Concentrate Kit (Zymo Research), and diluted to a concentration of 50 ng/µL. Gibson assembly was used to assemble the DNA encoding McbA variants with the pJL1 backbone. 10 ng of purified, linearized pJL1 backbone and 10 ng of gblock insert were combined in a 3-µL Gibson assembly reaction and incubated at 50 °C for 30 minutes^18^. The unpurified assembly reactions were diluted in 60 μL of NF water and 1 μL of the diluted reaction was used as the template for a 50-μL PCR reaction (using Q5 Hot Start DNA polymerase) to generate LETs for CFPS.

### Expression and purification of recombinant proteins

pJL1-McbA plasmid was transformed into chemically competent *E. coli* BL21 Star (DE3) cells (Invitrogen) following the manufacturer’s instructions. Cells were plated onto LB-Kan and incubated overnight at 37 °C. A single colony was used to inoculate a 5 mL overnight culture of LB-Kan, grown at 37 °C with 250 RPM shaking. 1 L of Overnight Express TB Medium (Millipore) was prepared following the manufacturer’s instructions and supplemented with 100 µg/mL kanamycin. The TB medium was inoculated the following day using the 5 mL overnight culture and grown at 37 °C with 250 RPM shaking until saturation (∼ 12-16 hours). Cells were harvested by centrifugation (Beckman Coulter Avanti J-26) at 8,000 x g for 10 min at 4 °C. Cell pellets were either flash frozen with liquid nitrogen and stored at -20 °C until future use or resuspended in 25 mL Wash Buffer (100 mM Tris-HCl pH 8.0, 150 mM NaCl, 1 mM EDTA, 10% v/v glycerol). Resuspended cells were lysed by sonication (QSonica Q700 Sonicator) using six 10 seconds ON and 10 seconds OFF cycles at 50% amplitude, and the insoluble fraction was removed by centrifugation at 12,000 x g for 20 minutes at 4 °C. Clarified lysates were incubated with 2 mL of pre-equilibrated Strep-Tactin XT Superflow resin (IBA Lifesciences) with shaking for 30 min at 4°C. Resin was loaded onto a gravity-flow column and washed three times with 20 mL Wash Buffer. McbA protein was eluted with 10 mL of Elution Buffer (100 mM Tris-HCl pH 8.0, 150 mM NaCl, 1 mM EDTA, 50 mM biotin, 10% v/v glycerol) and concentrated with a 15 mL Amicon Ultra Centrifugal filter (Millipore Sigma; 30 kDa cutoff). Purified McbA was buffer exchanged into Storage Buffer (50 mM HEPES pH 7.5, 300 mM NaCl, 10 mM MgCl_2_, 10% v/v glycerol) using a pre-equilibrated PD-10 desalting column (Cytiva). McbA was stored at 4 °C for immediate use (<48 hours) or -20 °C for longer term storage. Protein concentration was quantified by measuring A280 on a NanoDrop 2000c (Thermo Scientific), with McbA extinction coefficient and molecular weight calculated by Expasy ProtParam. wt-McbA and the six engineered McbA variants found in **Fig. 5** were purified in this manner.

### muGFP activity assay

Performance of muGFP variants were quantified by measuring fluorescence on a plate reader (BioTek Synergy H1) using an excitation of 485 nm and emission of 528 nm. 10 µL of crude CFPS reaction containing an expressed muGFP variant was transferred to a black, round bottom 384-well plate (Nunc) prior to measurements.

### Amide synthetase activity assay

All high-throughput assays (hot spot screen, iterative site saturation mutagenesis, substrate scope, ML predictions validation, and ML prediction exploration) were assembled in 384-well plates (Bio-Rad) using an Integra VIAFLO liquid handling robot. A 2x reaction mix containing the substrates (ATP, acid, amine, and DMSO) with excess volume filled with 50 mM potassium phosphate pH 7.5 was dispensed as 3-uL aliquots in a 384-well plate. The amidation assay was initiated by adding 3 µL of crude CFPS reaction containing an expressed McbA variant, with final concentrations of 25 mM ATP, 25 mM acid, 25 mM amine, 10% v/v DMSO, and ∼1 µM of enzyme (determined by ^14^C-leucine incorporation using previously described protocols^17^). Stock solutions of the acids were prepared in DMSO and this was taken into account to reach 10% v/v DMSO. For reactions that were performed in triplicates, 3 µL from the same 10-µL CFPS reaction was used for three separate assays. The reaction was incubated at 37 °C for 16 hours and then quenched with 25 µL of methanol. Plates were stored at -20 °C until prepared for analysis via LC-MS.

Amidation assays for the purified McbA variants found in **Fig. 5** were set up similarly as described above in 384-well plates. 8-µL reactions were assembled in triplicate, containing 25 mM ATP, 25 mM acid, 25 mM amine, 10 mM MgCl_2_, 10 U/mL pyrophosphatase (Sigma I5907), 0.5 mg/mL McbA, 10 % v/v DMSO, and volume to fill of 50 mM potassium phosphate pH 7.5. For assaying the production of cinchocaine and procainamide, substrates were decreased in stoichiometric amounts to 20 mM and 10 mM, respectively. This was to compensate for an observed poor solubility of these two acids (2-butoxyquinoline-4-carboxylic acid and 4-aminobenzoic acid) in the purified reaction at 10% v/v DMSO. Reactions were incubated at 37 °C for 16 hours and then quenched with 25 µL of methanol. Samples were stored at -20 °C until prepared for analysis via LC-MS. The CAS numbers of all chemicals used in the hot spot screens, as well as the amide standards we purchased, can be found in **Table S13**.

### Amide synthetase & ATP regeneration assay

Polyphosphate kinase, PPK12 from an unclassified *Erysipelotrichaceae* (Uniprot: A0A847P5F2_9FIRM), was cloned, expressed, and purified to homogeneity as previously described^58^. 20-µL reactions were assembled in triplicate, containing 25 mM amine, 25 mM acid, 100 mg/mL polyphosphate (Sigma 1.06529), 10 mM MgCl_2_, 10 U/mL pyrophosphatase (Sigma I5907), 0.5 mg/mL McbA, 0.5 mg/mL PPK12, 10 % v/v DMSO, and volume to fill of 50 mM potassium phosphate pH 7.5. A 2-fold serial dilution of AMP was prepared and added to the reaction mix to final concentrations ranging from 25 mM to 0.02 mM. Reactions were incubated at 37 °C for 16 hours and then quenched with 25 µL of methanol and analyzed by LC-MS.

### Preparative scale biosynthesis of moclobemide

Scaled amidation assays for the enzymatic preparation of moclobemide were set up similarly as described above. A 10-mL reaction containing 25 mM ATP, 25 mM acid, 25 mM amine, 10 mM MgCl_2_, 10 U/mL pyrophosphatase (Sigma I5907), 0.5 mg/mL McbA, 10 % v/v DMSO, and volume to fill of 50 mM potassium phosphate pH 7.5. After 16 hours, the reaction was quenched and product was extracted by the addition of 30 mL of ethyl acetate (3 x 10 mL). The organic phases were collected, washed with 0.2 M NaOH (2 x 10 mL), and brine (2 x 10 mL), dried over MgSO_4,_ filtered, and the solvent was evaporated under reduced pressure to afford the desired product as a white powder (58 mg, 87% isolated yield) without any further purification. The ^1^H and ^13^C NMR (found below and in **Fig. S12**) are in good agreement with those previously reported^59^. Spectra for ^1^H and ^13^C NMR were recorded at room temperature with a Bruker Avance III 500 MHz system. Chemical shifts are reported in δ (ppm) relative units to residual solvent peaks DMSO-d6 (2.50 ppm for 1H and 39.5 ppm for 13C). Splitting patterns are assigned as s (singlet), d (doublet), t (triplet), q (quartet), quint (quintet), m (multiplet). Coupling constants are reported as Hz, followed by integration.

**^1^H NMR** (500 MHz, DMSO-d6) δ 8.47 (t, J = 5.7 Hz, 1H), 7.82 – 7.75 (m, 2H), 7.51 – 7.45 (m, 2H), 3.50 (t, J = 4.6 Hz, 4H), 3.30 (d, J = 13.0 Hz, 3H), 2.38 (t, J = 7.0 Hz, 3H), 2.35 – 2.31 (m, 3H)

**^13^C NMR** (126 MHz, DMSO) δ 165.51, 136.38, 133.69, 129.55, 128.85, 66.66, 57.77, 53.76, 37.06.

### LC-MS analytics

Amide products (along with acid substrates and some adenylated acid intermediates) were analyzed using an Agilent G6125B Single Quadrupole LC/MSD system equipped with an electrospray ionization source set to positive ionization mode. The quenched samples were centrifuged for 10 min at 4,500 x g to remove precipitated proteins. A separate 384-well plate for sample injection into the HPLC-MS was prepared by diluting 5 µL of the quenched samples with 25 µL of methanol using the Integra VIAFLO. Trace amounts of compounds were detected using MS, while many compounds were present in high enough concentration to quantify by diode array detector (DAD) at 254 nm. Compounds were separated on a Luna C18 Column (Phenomenex 00D-4251-B0) using mobile phases (A) H_2_O with 0.1% formic acid and (B) Acetonitrile. The general method for chromatographic separation was carried out using the following gradients at a constant flow rate of 0.5 mL/min: 0 min 5% B; 1 min 5% B; 4 min 95% B; 4.5 min 95% B; 5 min 5% B. For hot spot screens, an expediated method was used with the following gradients at a constant flow rate of 0.5 mL/min: 0 min 13% B; 1 min 13% B; 2.2 min 95% B; 3.2 min 95% B; 3.5 min 13 % B. For the MS, capillary voltage was set at 3 kV, and nitrogen gas was used for nebulizing (35 psig) and drying (12 l/min, 350 °C). The MS was calibrated using Tuning Mix (Agilent G2421-60001) before measurements were taken. MS data were acquired with a scan range of 50-600 *m/z* with various SIM *m/z*’s according to which compound we were screening for. LC-MS data were collected and analyzed using Agilent OpenLab CDS ChemStation software. The product yield was estimated by dividing the DAD peak area for the amide product by the sums of the peak areas of both the amide and the acid substrate. An exact quantitative yield for moclobemide was recorded after its preparative scale synthesis and isolation.

### Melting temperature determination

Protein melting temperature was determined using a Jasco J-810 circular dichroism spectrophotometer with a 10 mm path length cuvette monitored at 222 nm. McbA samples were first buffer exchanged into a 1X phosphate buffered saline solution, pH 7.4, and diluted to 0.2-0.4 mg/mL.

### Enzyme kinetics

McbA apparent kinetics for the amine pair of moclobemide (4-(2-aminoethyl)morpholine) were determined by enzymatically coupling amide bond formation (and the concomitant release of AMP from the acyl-AMP intermediate by its substitution with the amine) with the oxidation of NADH (**Fig. S9**). Reactions contained 100 mM MOPS-KOH pH 7.8, 5 mM MgCl_2_, 2.5 mM phosphoenolpyruvate, 5 mM ATP, 0.3 mM NADH, 50 mM 4-chlorobenzoic acid, 15 U/mL pyruvate kinase and lactate dehydrogenase enzyme mix (Sigma-Aldrich P0294), 25 U/mL myokinase (Sigma-Aldrich 475941), and various concentrations (50-200 µg/mL) of the studied McbA variant. As the acid here (4-chlorobenzoic acid) has poor solubility in water and was dissolved in DMSO, the final reactions contained 10% v/v DMSO (equivalent to our amidation screens). 180-µL reactions were first equilibrated at 30 °C for 3 minutes and then initiated by adding 20 µL of amine. The initial velocity was determined for different concentrations of amine (0.1 mM – 50 mM) by measuring NADH absorbance at 340 nM on a Cary 60 UV-Vis (Agilent). Data was collected and analyzed using the Cary WinUV Kinetics Application software (Agilent). Michaelis-Menten graphs were plotted in GraphPad Prism and fit using the default Michaelis-Menten non-linear regression analysis tool.

Kinetics for the acid pair of moclobemide (4-chlorobenzoic acid) were measured similarly as described above, except the amine was held constant at 50 mM and the reaction was initiated by addition of various amounts of the acid. The final DMSO concentration was still held constant at 10% v/v. We observed non-Michaelis-Menten behavior when attempting to determine the kinetics for the acid, in what appeared to be substrate inhibition by the acid (data not shown). We also attempted to measure the acid adenylation step directly by enzymatically coupling acyl-AMP formation (and the concomitant release of PP_i_) with the oxidation of NADH to further probe the reaction mechanism. The Piper™ pyrophosphate assay kit (Fisher Scientific P22062) was used, but the addition of small concentrations of DMSO resulted in the precipitation of enzymes found in the kit.

### Amino acid encodings

Five different amino acid encoding strategies were studied here following the work of Wittman *et al.* and *Vornholt et al.*^14,60^: one-hot, Georgiev, VHSE, z-scales, and physical descriptors. Beyond one-hot encodings (that contain no information about the nature of the amino acid at each position), we also wanted to include encodings that attempt to encapsulate physiochemical properties of amino acids. We briefly explain these encodings below (in order of most to least parameters) and encourage readers to visit these sources for further information. To make informative numerical representations of amino acid properties, these strategies perform principal component analysis (PCA) of different manually curated sets of either experimentally measured or computationally predicted/estimated properties. Georgiev^42^ features (19-parameters) are principal components of the over 500 amino acid indices taken from the AAindex database. VHSE^43^ features (8-parameters) are principal components of 50 variables, focused on hydrophobic, steric, and electronic properties. Z-scales^44^ (5-parameters) features are principal components of 26 variables, focused on lipophilicity, size, and polarity. Physical descriptors^45,61^ (3-parameters) features are derived from a rational ad hoc modification of principal components of hydrophobic and steric properties of peptides. For all strategies, we first generated encodings for the entire combinatorial library tested (stored in a tensor of “4^20^ unique variants” x “4 amino acids” x “*n-*parameters”, where *n*-parameters is equal to the number of amino acids for one-hot). The last two dimensions of the tensor were then flattened to generate a matrix. Specifically for the physiochemical encodings (excluding one-hot), each column of the matrix was standardized (mean-centered and unit-scaled).

### Zero-shot predictions

Evolutionary: The EVmutations^35^ probability density model was trained using the EVcouplings webserver (https://evcouplings.org/) with default parameters, with the input sequence for McbA taken from UniProt (R4R1U5). The model we selected had a bitscore inclusion threshold of 0.7. The model and code for replicating zero-shot predictions are provided in our GitHub repository. The mutation effects prediction code provided in the EVcouplings GitHub repository (https://github.com/debbiemarkslab/EVcouplings/blob/develop/notebooks/model_parameters_m <u>utation_effects.ipynb</u>) was used as a template. Features for the augmented models were derived from the sequence statistical energy relative to wild type.

Universal: Predictions using the ESM-1b^46^ pre-trained transformer language model were made using the code provided from the excellent work of Wittman *et. al* on machine learning-guided directed evolution (https://github.com/fhalab/MLDE) with the ESM-1b model provided in the ESM GitHub repository (https://github.com/facebookresearch/esm). Briefly, a mask-filling protocol was used to predict the probability of different mutants by presenting the model with the entire sequence and “masking” a position of interest. We used a naïve mask-filing approach, which considers each variable position as independent from each other. This mask-filing approach was used as it is less computationally expensive and provided slightly superior predictions than a conditional approach (which does not assume independence of variable positions) in this previous work. A complete description of the code can be found in the original publication and the associated GitHub repository. Features for the augmented models were derived from the sequence log-probability relative to wild type.

Structural: Structural-based predictions were made using the MAESTRO^47^ command line tool for Windows (v1.2.35). We used the Protein Data Bank (PDB) structure for McbA (6SQ8) as the input and calculated changes in stability (unfolding free energy) with the ‘evalmut’ command. Features for the augmented models were derived using the ‘energy’ output.

### Machine learning-guided directed evolution

Ridge regression models were augmented following the code accompanying the elegant work of Hsu *et al.*^41^ (https://github.com/chloechsu/combining-evolutionary-and-assay-labelled-data).

McbA variant sequence featurization was performed by concatenating zero-shot predictions with site-specific amino acid encodings. Zero-shot predictions were first standardized and regularized by a common regularization strength (10^-8^). The L2 regularization strength for ridge regression (α) was determined during hyperparameter tuning using cross-validation. For our complete code used in this work, please see our accompanying GitHub repository at https://github.com/grantlandwehr/accelerated-enzyme-engineering. Given some changes made between initial model development and reimplementation of the code for publication (e.g. hyperparameter tuning cross validation scheme, search range of the regularization coefficient α, etc.) there are minor differences in predictions ranked 23-25 for metoclopramide and moclobemide found in **Fig. 4**.

Model evaluation and selection were first performed retrospectively by using the assay-labeled datasets from our moclobemide and metoclopramide engineering campaigns. Augmented models (using combinations of the above zero-shot predictors and amino acid encodings) were trained on the single site saturation libraries for four residues (*n* ≈ 80) and tested on the withheld higher-order mutants from the additional rounds of saturation mutagenesis (*n* ≈ 200). Hyperparameter turning of α was performed using repeated 5-fold cross-validation (with 20 repeats) by randomly sampling 80% of the training data and testing on the withheld 20%; model performance was evaluated using mean squared error (MSE). With the optimized hyperparameter, all trained models were used to make predictions on the withheld test set. Spearman correlation coefficient and NDCG were used to select the best zero-shot predictor and encoding strategy, with a preference given to NDCG.

After identifying the best model (which in our case was augmenting the EVmutation probability density model with Georgiev encodings), we made predictions on the entire combinatorial dataset (*n* = 160,000). The top 25 predictions for moclobemide and metoclopramide were then experimentally tested (**Fig. 4**). Model training and predictions for the remaining seven amide products was performed similarly as above.

### Data collection and analysis

All statistical information provided in this manuscript is derived from *n* = 3 independent experiments unless otherwise noted in the text or figure legends. Error bars represent 1 s.d. of the mean derived from these experiments. Data analysis and figure generation were conducted using Excel Version 2304, ChimeraX Version 1.5^62^, GraphPad Prism Version 9.5.0, and Python 3.9 using custom scripts available on GitHub. muGFP fluorescence was measured on a BioTek Synergy H1 Microplate Reader and analyzed using Gen5 Version 2.09.2. Autoradiograms were performed as previously described and scanned using the Typhoon FLA 7000 Imager v1.2^63^.

## Acknowledgments

We thank Kosuke Seki, Andrew C. Hunt, and Steve R. Fleming for conversations regarding this work. We acknowledge the use of the Keck Biophysics Facility, a shared resource of the Robert H. Lurie Comprehensive Cancer Center of Northwestern University supported in part by the NCI Cancer Center Support Grant #P30 CA060553. In addition, we acknowledge the use of the computational resources and staff contributions provided for the Quest high performance computing facility at Northwestern University which is jointly supported by the Office of the Provost, the Office for Research, and Northwestern University Information Technology. This work also made use of the IMSERC NMR facility at Northwestern University, which has received support from the Soft and Hybrid Nanotechnology Experimental (SHyNE) Resource (NSF ECCS-2025633), Int. Institute of Nanotechnology, and Northwestern University. Molecular graphics and analyses performed with UCSF ChimeraX, developed by the Resource for Biocomputing, Visualization, and Informatics at the University of California, San Francisco, with support from National Institutes of Health R01-GM129325 and the Office of Cyber Infrastructure and Computational Biology, National Institute of Allergy and Infectious Diseases.

## Funding

Department of Energy (DE-SC0023278)

Defense Threat Reduction Agency (HDTRA1-21-1-0038)

National Institutes of Health (1U19AI142780-01)

## Author contributions

Conceptualization: ASK, GML, JWB, MCJ

Methodology: GML, JWB

Investigation: CM, EGH, GML, JWB

Software: GML

Funding acquisition: ASK, MCJ

Supervision: ASK, MCJ

Writing: ASK, GML, JWB, MCJ

## Data and materials availability

All data are available in the main text or the supplementary materials.

