## Supplementary Information for "Accelerated enzyme engineering by machine-learning guided cell-free expression"

**The PDF file includes:**

Figs. S1 to S25
Tables S1 to S13
References

### Table of Contents

**Supplemental Figures** .......... **3-28**
Substrate scope of wt-McbA .......... 3
Cell-free library generation validation .......... 4-8
Engineering campaign for moclobemide .......... 9
Kinetic and stability characterization of $McbA_{moc}$ .......... 10-12
Scale-up and compound NMR spectra of moclobemide .......... 13-15
Engineering campaigns for cinchocaine and metoclopramide .......... 16-17
ML model selection .......... 18
HSS of expanded substrate set .......... 19-26
Characterization of ML-predicted McbA variants .......... 27-28
**Supplemental Tables** .......... **29-50**
Screening all ML-predicted McbA variants .......... 29-33
DNA sequences of McbA variants .......... 34-41
Cell-free DNA assembly .......... 42-43
Primers used in this study .......... 44-48
BLOSUM50 reduced amino libraries .......... 49
CAS number of compounds used in this study .......... 50
**References** .......... **51**

### Supplementary Figures

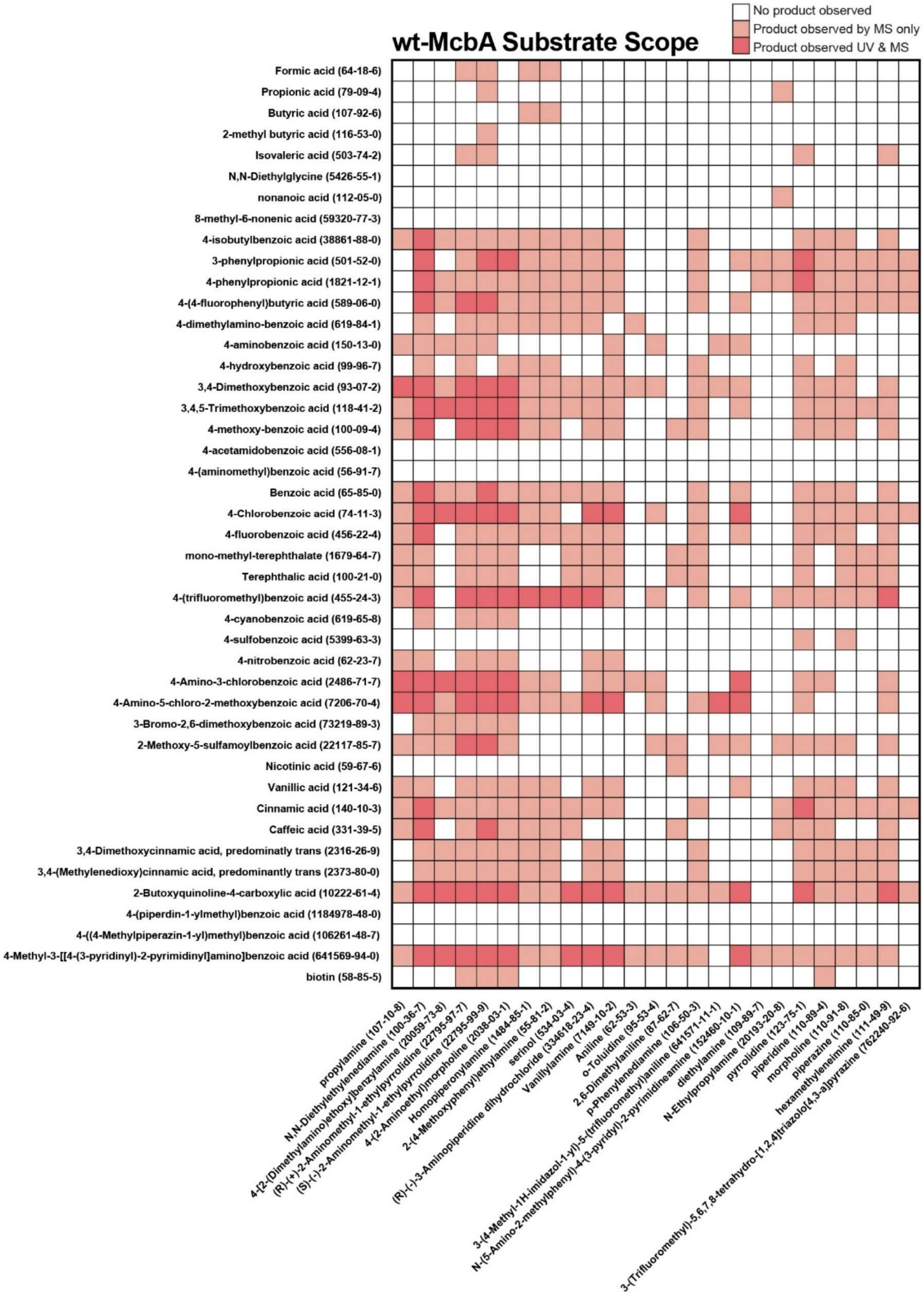


**Figure S1. Substrate scope of wt-McbA.** This figure key corresponds to the substrate scope in **Fig. 2**. All acids (y-axis) and amines (x-axis) tested and their CAS numbers are included. Data shown are from one experiment (*n* = 1).

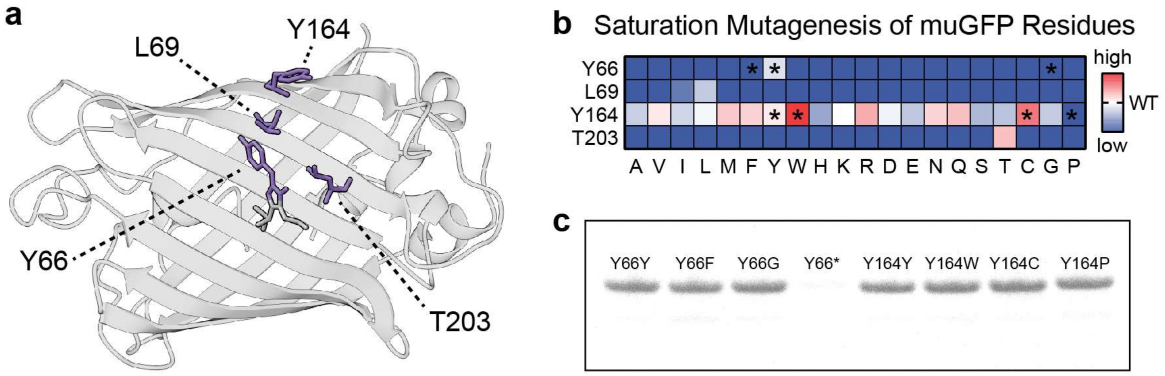


**Figure S2. Validation of the cell-free protein expression workflow with muGFP. a,** Crystal structure of monomeric ultra-stable green fluorescent protein (muGFP) with residues targeted for saturation mutagenesis highlighted (PDB: 5JZL). **b,** Green fluorescence of single mutants of muGFP generated by site saturation mutagenesis. Data are showing mean fluorescence for *n* = 3 replicates normalized to wild type (wt-muGFP). **c,** Autoradiogram of selected mutants (denoted with asterisks in **b**) showing full-length, soluble proteins are expressed, except in the case when the mutation is a premature stop codon (Y66*). Data shown in the gel are representative of three independent experiments.

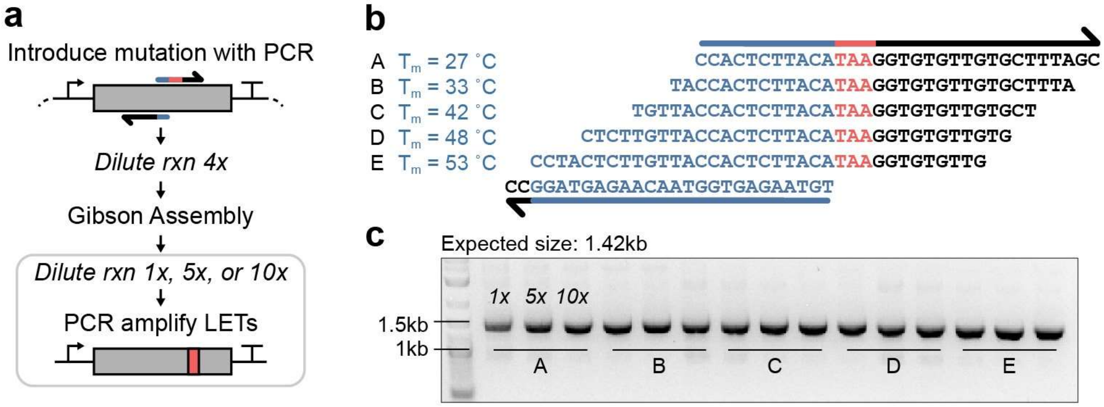


**Figure S3. Cell-free DNA assembly efficiency is robust to differences in homologous overlaps of mutagenic primers. a,** Schematic of cell-free DNA synthesis and mutation strategy. During protocol development, we considered and made modifications to increase efficiency of the Gibson assembly step: (i) the dilution of the Gibson assembly reaction (serving as the template for the second PCR) and (ii) the amount of homology between the forward and reverse primer in the first PCR. **b,** Five forward primers containing homologous overlaps with a single reverse primer were designed, each overlap differing in $T_m$ by approximately 5 °C. The primers for this experiment were used to introduce a stop codon at Y66 in muGFP. **c,** DNA gel showing the products of the second PCR using primers in **b** and dilutions in **a**. Linear expression templates (LETs) of the expected size are observed. Successful mutagenesis to a stop codon (TAA) was confirmed by Sanger sequencing of the produced LET. Data shown in the gel are representative of three independent experiments.

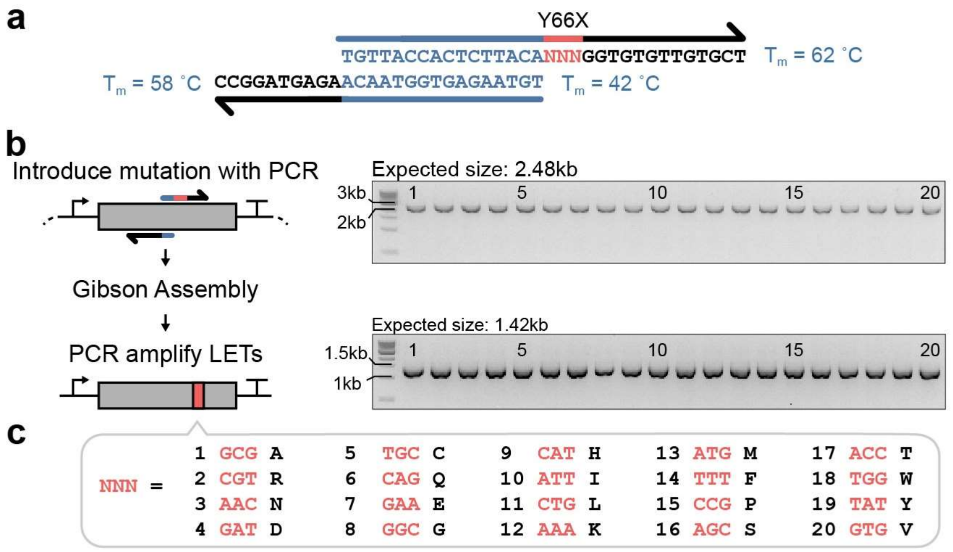


**Figure S4. Optimized reaction conditions for cell-free DNA assembly enable site-saturation mutagenesis. a,** Primer design schematic. Heuristics are based on primer melting temperatures and were guided by the optimization performed in **Fig. S3.** Shown here are the designed primers for the site saturation of muGFP Y66 with NNN indicating the targeted codon. Briefly, the forward primer containing the desired mutation has a Tm of ~62 °C, the reverse primer has a Tm of ~58 °C, and the homologous overlap between the forward and reverse has a Tm of ~42 °C. **b,** DNA gels demonstrating the expected product size for the two PCR steps of the site saturation of muGFP Y66. Top: linearized plasmid with integrated mutation. Bottom: amplified linear expression template (LET) with integrated mutation. Data shown are representative of three independent experiments. **c,** Amino acid table containing all codons used for site saturation mutagenesis in this study, numbered according to the gels in **b**. The codons correspond to the most prevalent codons found in *E. coli* and were held constant. We elected to not perform codon optimization as (i) tRNAs are supplemented in excess in CFPS reactions so we would not expect a single codon to significantly alter translation and (ii) holding codons constant reduces the complexity of primer design when scaling to numerous mutations.

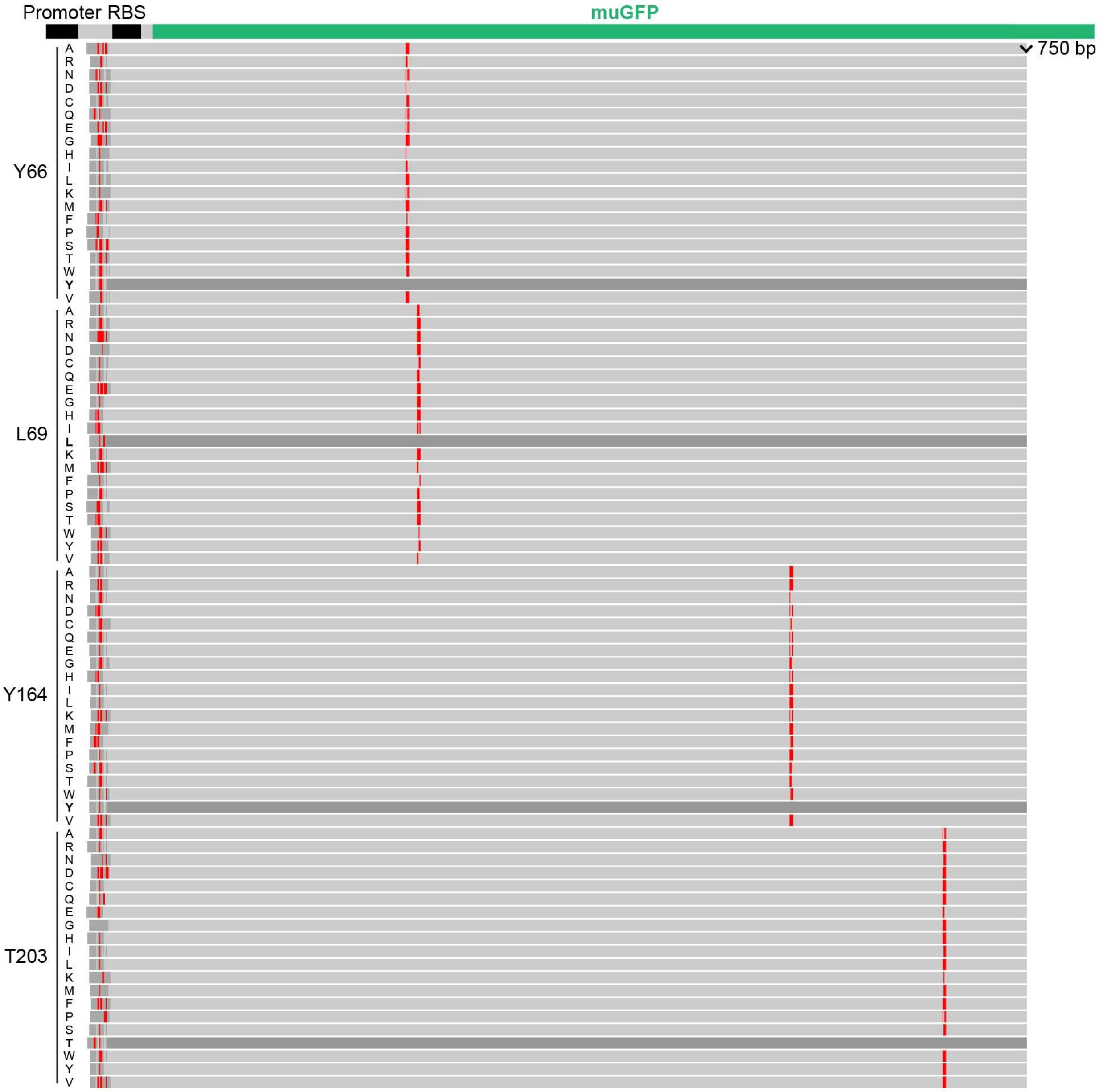


**Figure S5. Sequence alignment of muGFP four residue site saturation.** The Sanger Sequencing alignment for every muGFP mutant found in **Fig. S2b** generated using our cell-free DNA assembly method aligned to WT muGFP (top; promoter, RBS, and muGFP sequences are highlighted for convenience). Portions of the sequence alignment highlighted in red are mismatches compared to WT muGFP and all correspond to the expected mutation listed on the left-hand side of the alignment.

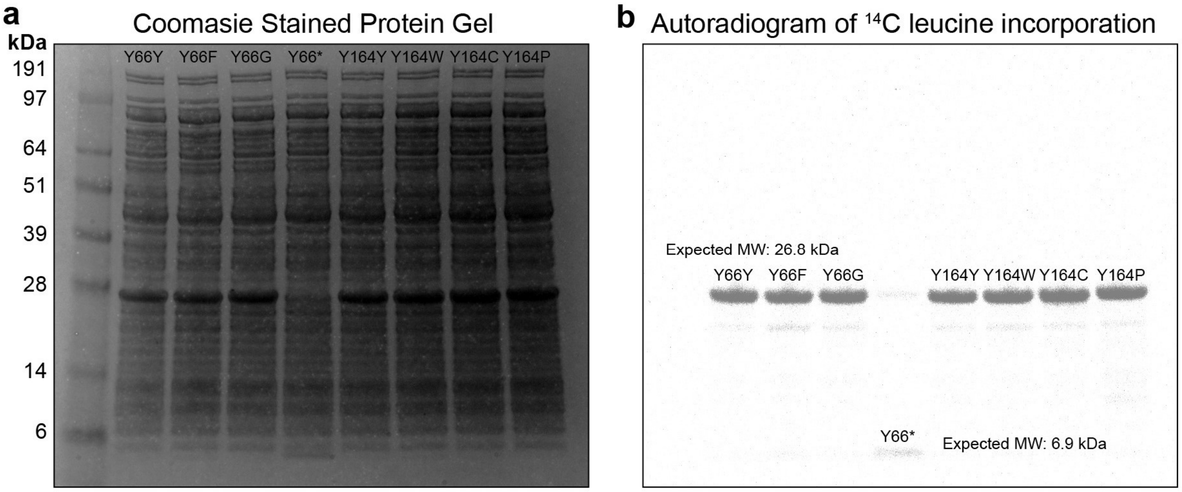


**Figure S6. Protein gels of selected muGFP single mutants with variable measured green fluorescence.** Complete protein gels of the truncated image shown in **Fig. S2c** depicting selected muGFP mutants from the site saturation mutagenesis screen. Mutants were expressed using CFPS with a radioactive leucine ($^{14}$C-leucine) to distinguish the expressed protein among the *E. coli* proteome present in CFPS. Two residues were selected (Y66 and Y164) and a stop codon (Y66*) was included as a control to produce a truncated protein. **a**, Coomassie stained protein gel of muGFP expressed in CFPS. **b,** The same gel imaged as an autoradiogram. Data shown are representative of three independent experiments.

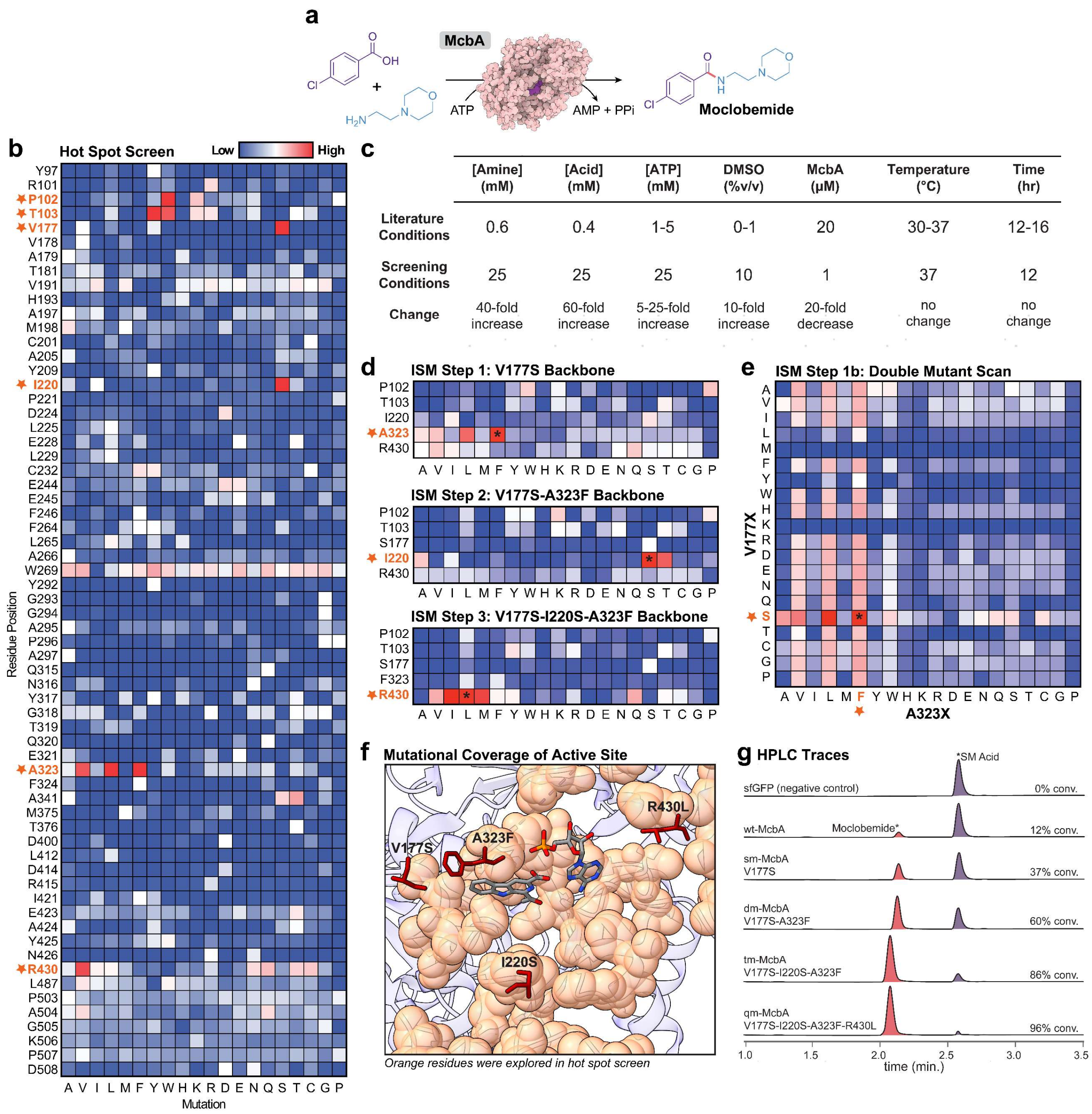


| | [Amine] (mM) | [Acid] (mM) | [ATP] (mM) | DMSO (%v/v) | McbA (μM) | Temperature (°C) | Time (hr) |
|---|---|---|---|---|---|---|---|
| Literature Conditions | 0.6 | 0.4 | 1-5 | 0-1 | 20 | 30-37 | 12-16 |
| Screening Conditions | 25 | 25 | 25 | 10 | 1 | 37 | 12 |
| Change | 40-fold increase | 60-fold increase | 5-25-fold increase | 10-fold increase | 20-fold decrease | no change | no change |

**Fig. S7. Engineering campaign for the small-molecule drug moclobemide. a,** Reaction scheme for engineering McbA to produce moclobemide. **b,** Hot spot screen of 64 identified residues in McbA showing percent conversion of moclobemide normalized to WT (*n* = 1). Highly mutable sites are starred and included in additional engineering rounds. **c,** Screening conditions for engineering McbA towards more industrially relevant conditions. **d,** Iterative site saturation mutagenesis of residues identified in the HSS, again showing normalized percent conversion of moclobemide (*n* = 1). **e,** Comprehensive double mutant scan of residues identified in the first two rounds of ISM, again showing normalized percent conversion of moclobemide (*n = 1). **f,** Crystal structure of McbA (PDB: 6SQ8) with bound native substrates. Orange residues highlight the near complete coverage of first shell residues in the active site explored in the HSS. g**,** Reversed-phase (RP)-HPLC traces of moclobemide product (red) and acid substrate (purple) of wt-McbA and engineered mutants. HPLC trace representative of three independent reactions (*n* = 3).

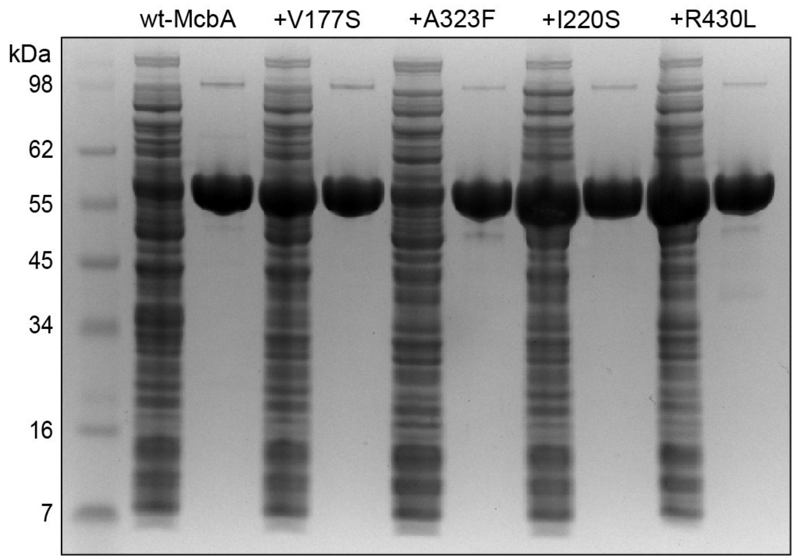


**Figure S8. Coomassie stained SDS-PAGE of purified McbA$_{moc}$ variants.** McbA variants were expressed in *E. coli* BL21 (DE3) and purified using an N-terminal strep tag. The first lane is the lysate soluble fraction of the overexpressed protein and the second lane is the purified protein. The expected molecular weight was 55.5 kDa, and the purified protein lane was loaded with 10 µg of protein.

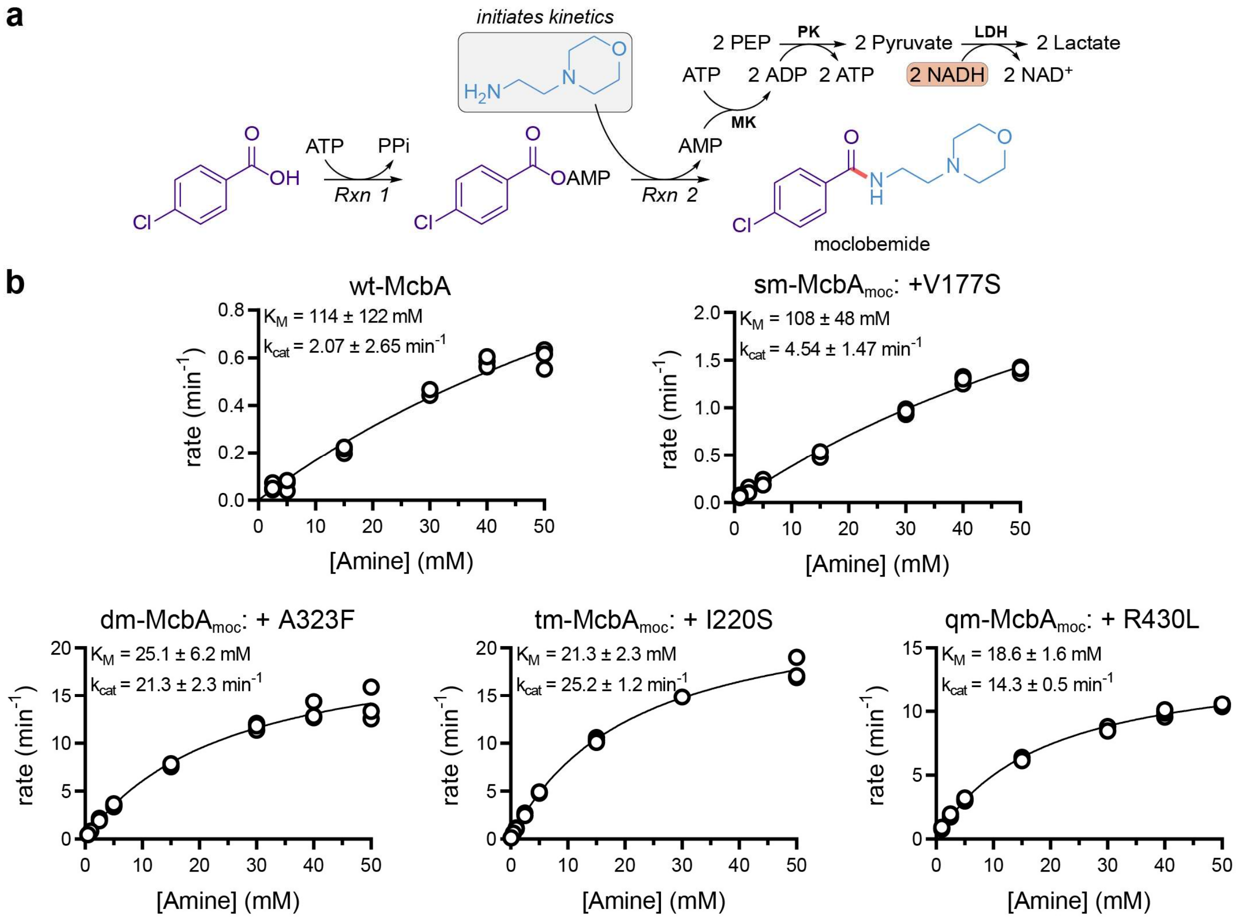


**Figure S9. Michaelis-Menten graphs of McbA$_{moc}$ variants. a,** The coupled enzymatic reaction scheme we used to measure the Michaelis-Menten kinetics of the amine substrate (4-(2-aminoethyl)morpholine). The reaction was first equilibrated at 30 °C in the presence of the acid (50 mM of 4-chlorobenzoic acid) and ATP (5 mM). The kinetic measurements were initiated by adding varying amounts of amine (0.5 – 50 mM) and the rate was determined by measuring NADH oxidation at 340 nM. **b,** Michaelis-Menten graphs of all McbA$_{moc}$ variants were plotted in GraphPad Prism and fit using the default Michaelis-Menten non-linear regression analysis tool. For both wt-McbA and sm-McbA, we were unable to reach a saturating concentration of amine and therefore have high uncertainty in the estimated kinetic parameters. All data replicates are shown as independent experimental points (*n* = 3).

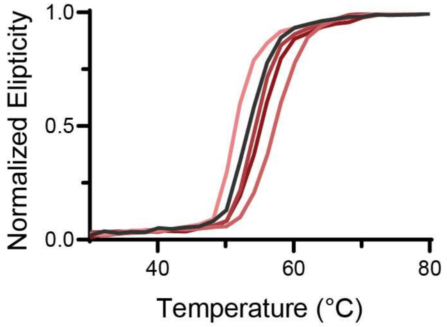


| Variant | $T_m$ (°C) |
|---|---|
| wt-McbA | 53.59 ± 0.05 |
| +V177S | 51.56 ± 0.08 |
| +A323F | 57.46 ± 0.04 |
| +I220S | 54.51 ± 0.07 |
| +R430L | 55.16 ± 0.05 |

**Figure S10. Melting temperatures of McbA$_{moc}$ variants.** Circular dichroism (CD) denaturation curves were min-max normalized for each sample for ease of comparison across mutants. Data shown are representative of three independent experiments.

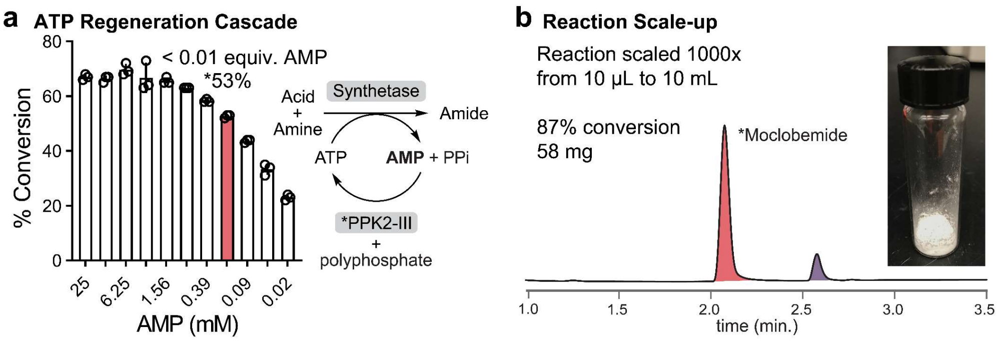


**Fig. S11. Preparative-scale synthesis of moclobemide with an engineered McbA. a,** An additional ATP regeneration system sustains high percent conversions using low stoichiometric addition of AMP ($n$ = 3)[1,2]. **b,** Preparative scale synthesis of moclobemide with purified product pictured.

**Figure S12. Compound NMR spectra of purified moclobemide synthesized by McbA.**

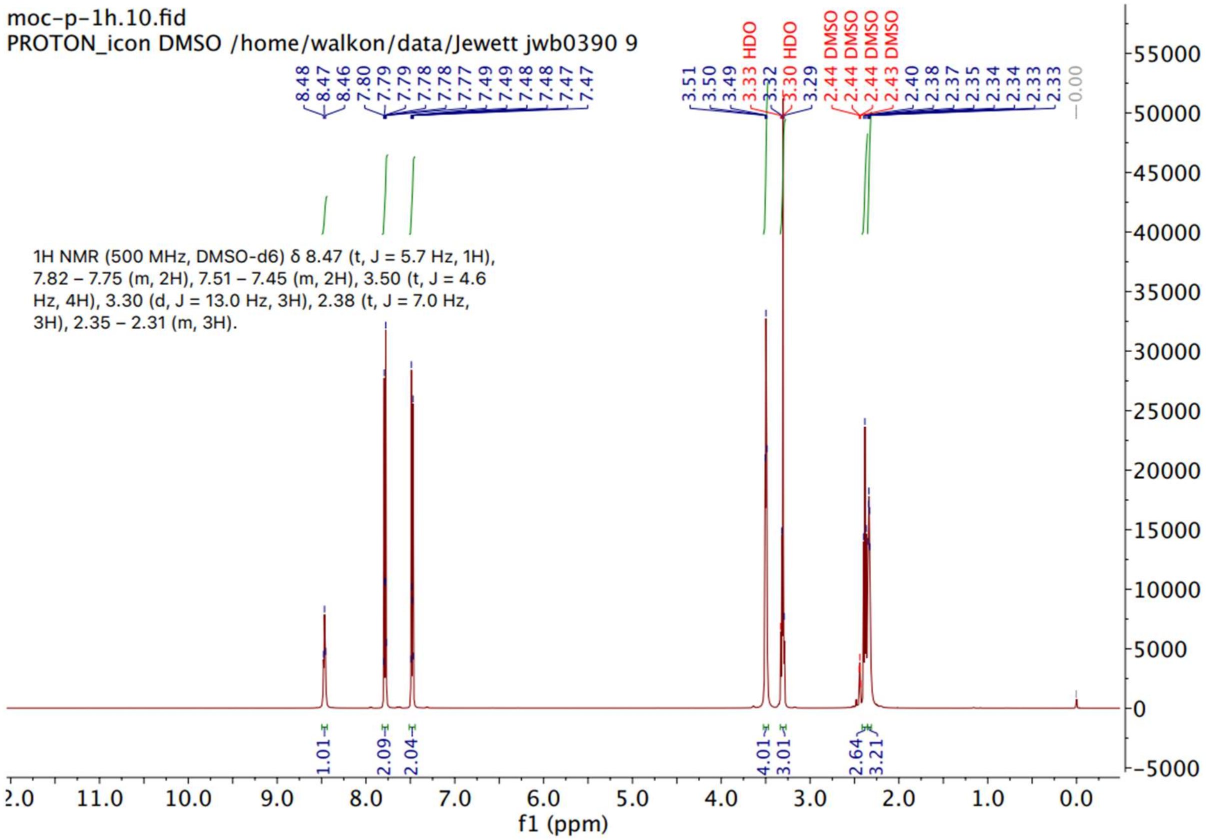

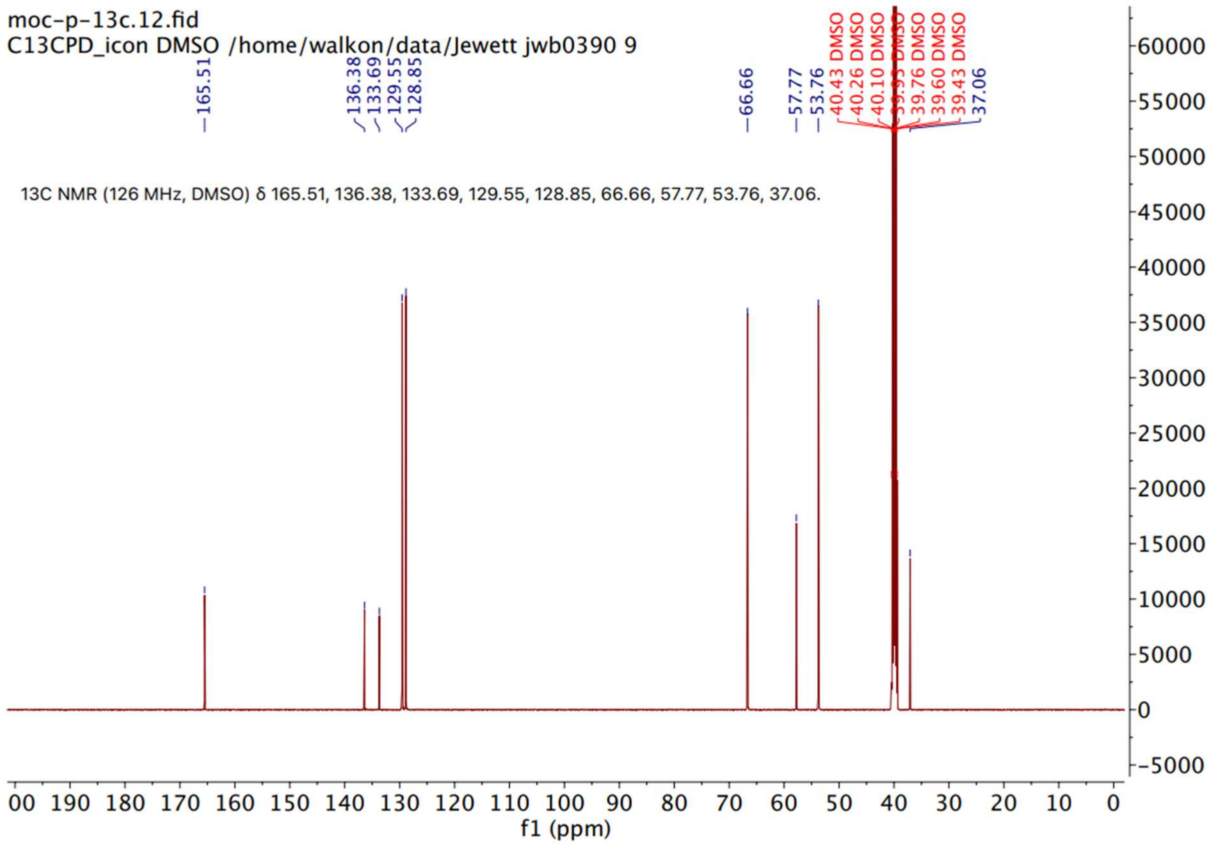
moc-p-13c.12.fid
C13CPD_icon DMSO /home/walkon/data/Jewett jwb0390 9
165.51
136.38
133.69
129.55
128.85
66.66
57.77
53.76
40.43 DMSO
40.26 DMSO
40.10 DMSO
39.93 DMSO
39.76 DMSO
39.60 DMSO
39.43 DMSO
37.06
13C NMR (126 MHz, DMSO) δ 165.51, 136.38, 133.69, 129.55, 128.85, 66.66, 57.77, 53.76, 37.06.
f1 (ppm)

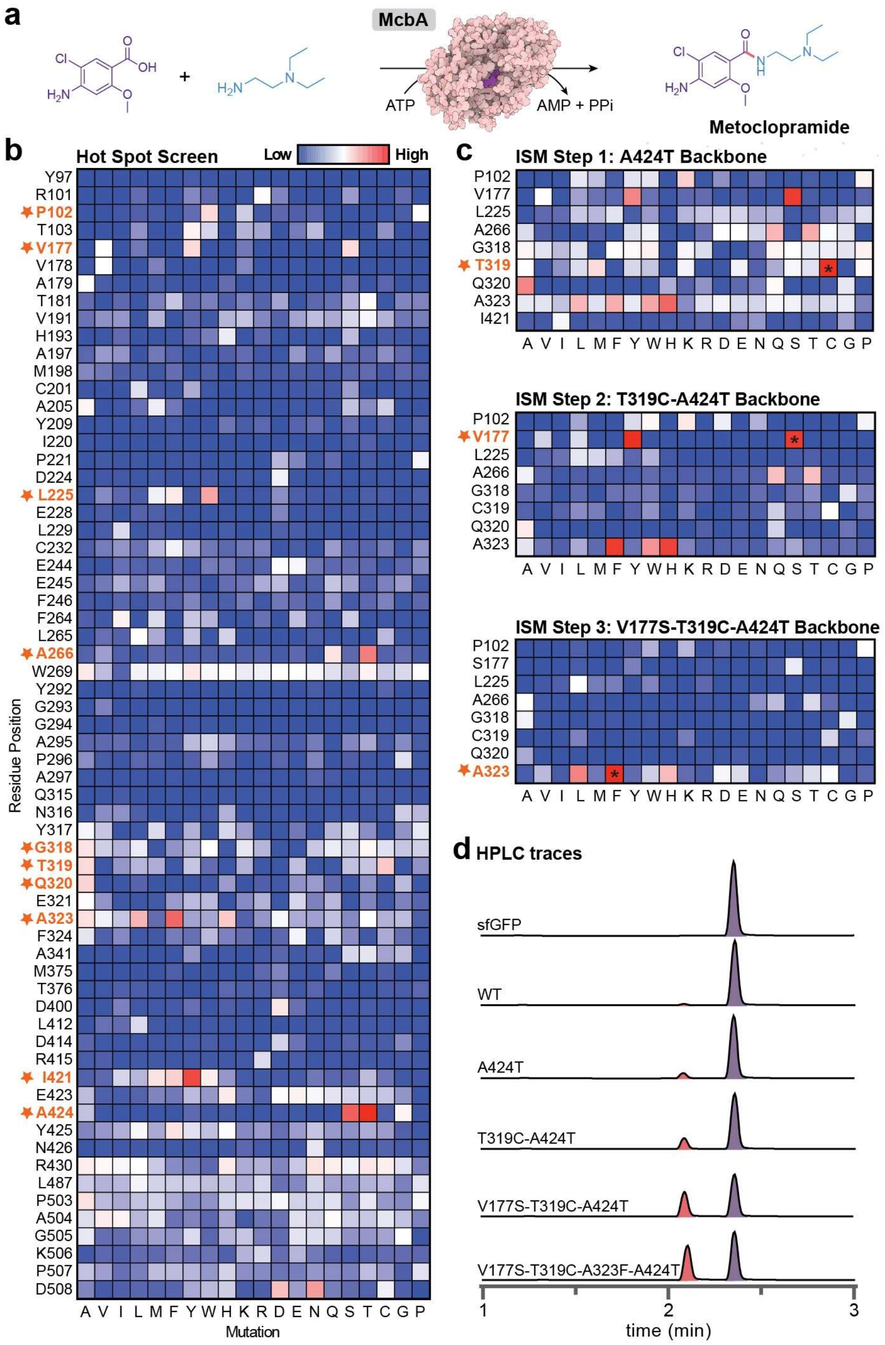


**Figure S13. Engineering campaign for the small-molecule drug metoclopramide. a,** Reaction scheme for the biosynthesis of metoclopramide using McbA. **b,** HSS of 64 identified residues in McbA showing percent conversion of metoclopramide normalized to WT ($n$ = 1) as determined by absorbance on HPLC. Highly mutable sites are starred and included in additional engineering rounds. **c,** Three additional ISM rounds of residues that were identified in the HSS. The mutation that had the greatest positive impact on activity is starred and was included in the backbone for the next round. **d,** RP-HPLC trace of metoclopramide product (red) and acid substrate (purple) of wt-McbA and engineered mutants. HPLC trace data are representative of three independent experiments.

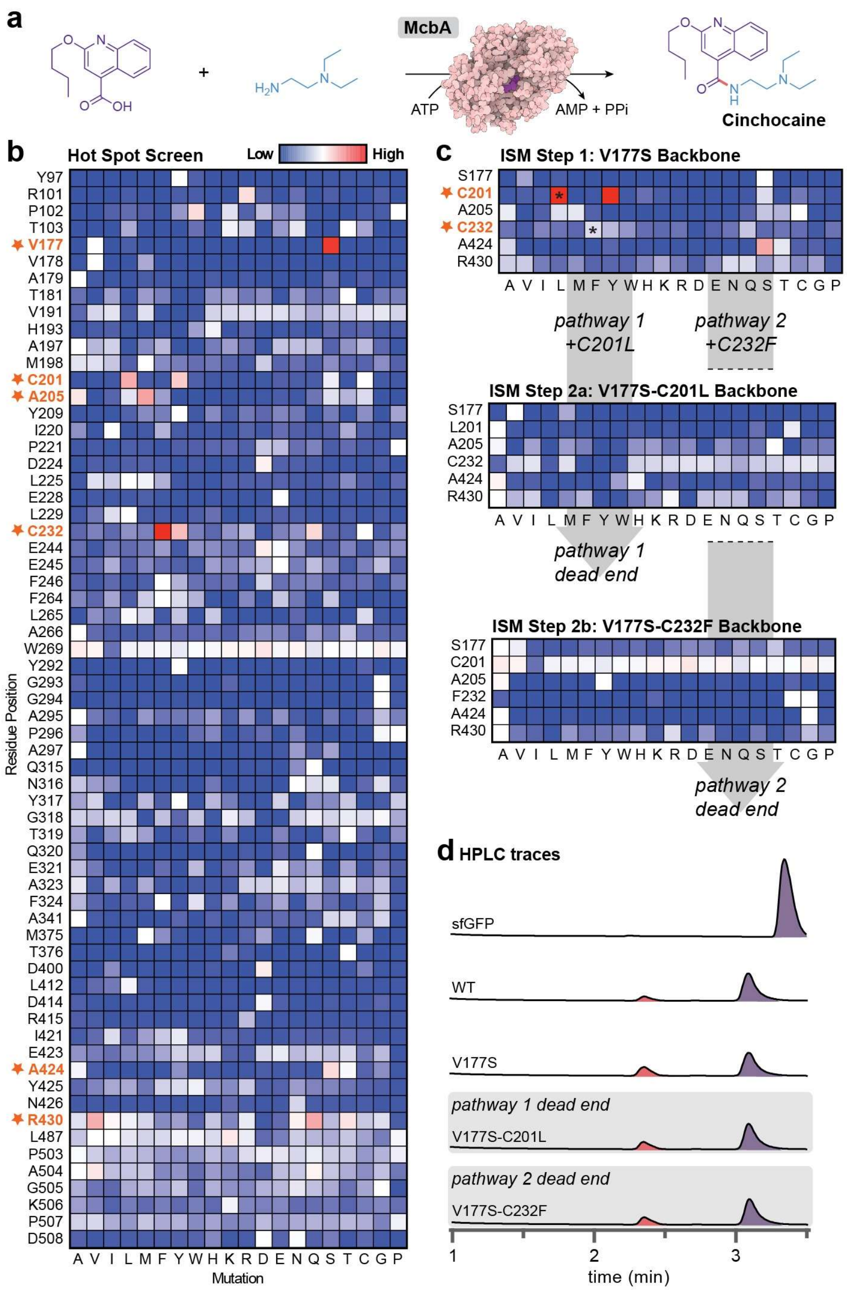


**Figure S14. Engineering campaign for the small-molecule drug cinchocaine. a,** Reaction scheme for the biosynthesis of cinchocaine using McbA. **b,** HSS of 64 identified residues in McbA showing percent conversion of cinchocaine normalized to WT (*n* = 1) as determined by absorbance on HPLC. Highly mutable sites are starred and included in additional engineering rounds. **c,** After one additional engineering round, we were unable to find an additional beneficial mutation beyond the double mutant (pathway 1). Notably, mutations that were previously observed to be beneficial were no longer in future rounds. To overcome this "dead end", we performed an additional engineering round using a double mutant consisting of the two best mutations found in the HSS (pathway 2). We also could not find additional mutations for this backbone and reached another "dead end". **d,** RP-HPLC trace of cinchocaine product (red) and acid substrate (purple) of wt-McbA and engineered mutants. HPLC trace data are representative of three independent experiments.

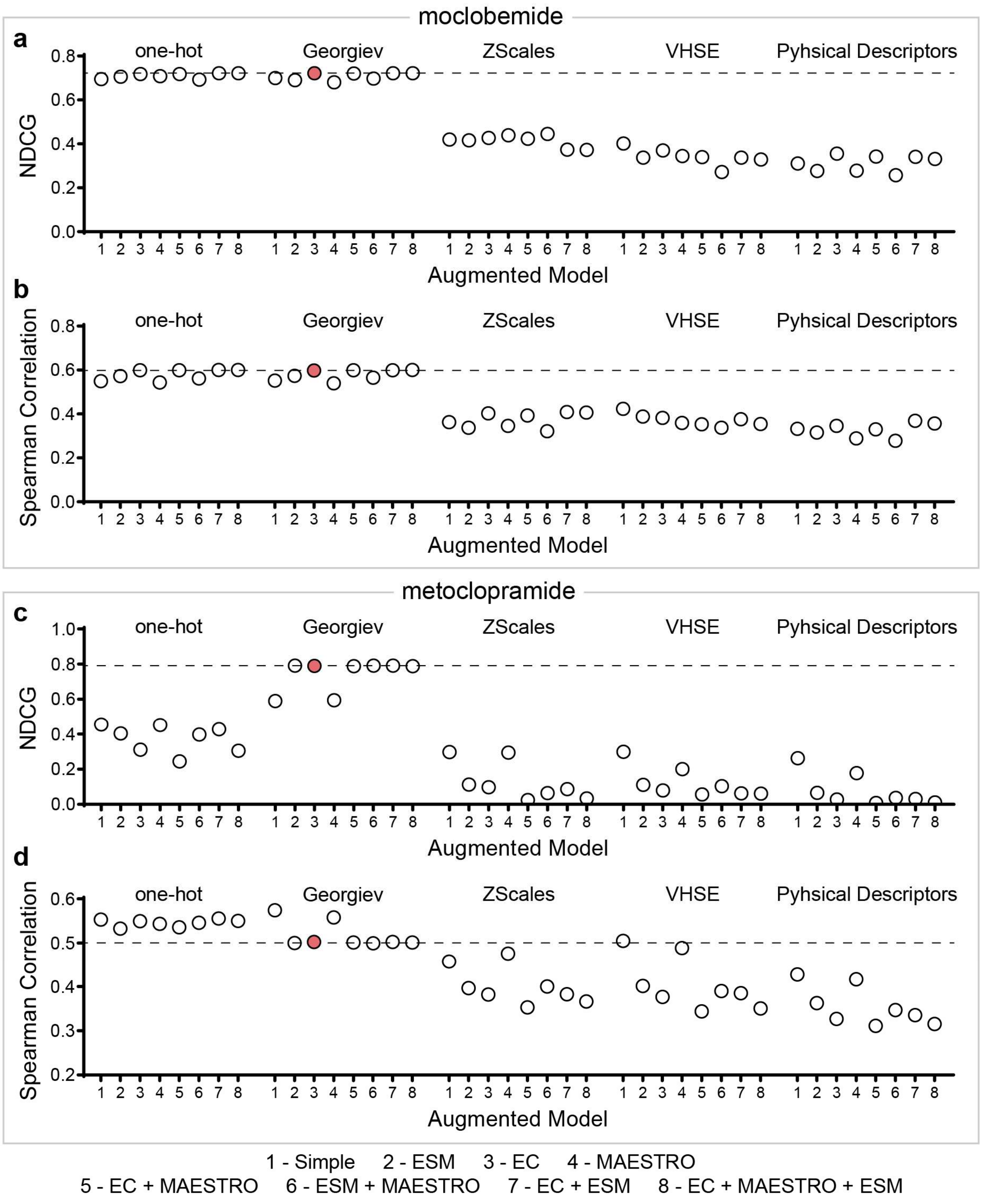


**Figure S15. Complete characterization of augmented ridge regression model performance using different combinations of fitness predictors and amino acid encodings.** While we explicitly used NDCG as our selection criteria for model performance, we also include the Spearman correlation coefficient to better explore the differences in all the models tested. In most instances, these two model performance indicators are closely aligned. The top graphs display NDCG (**a**) and Spearman correlation (**b**) for moclobemide and the bottom graphs display NDCG (**c**) and Spearman correlation (**d**) for metoclopramide.

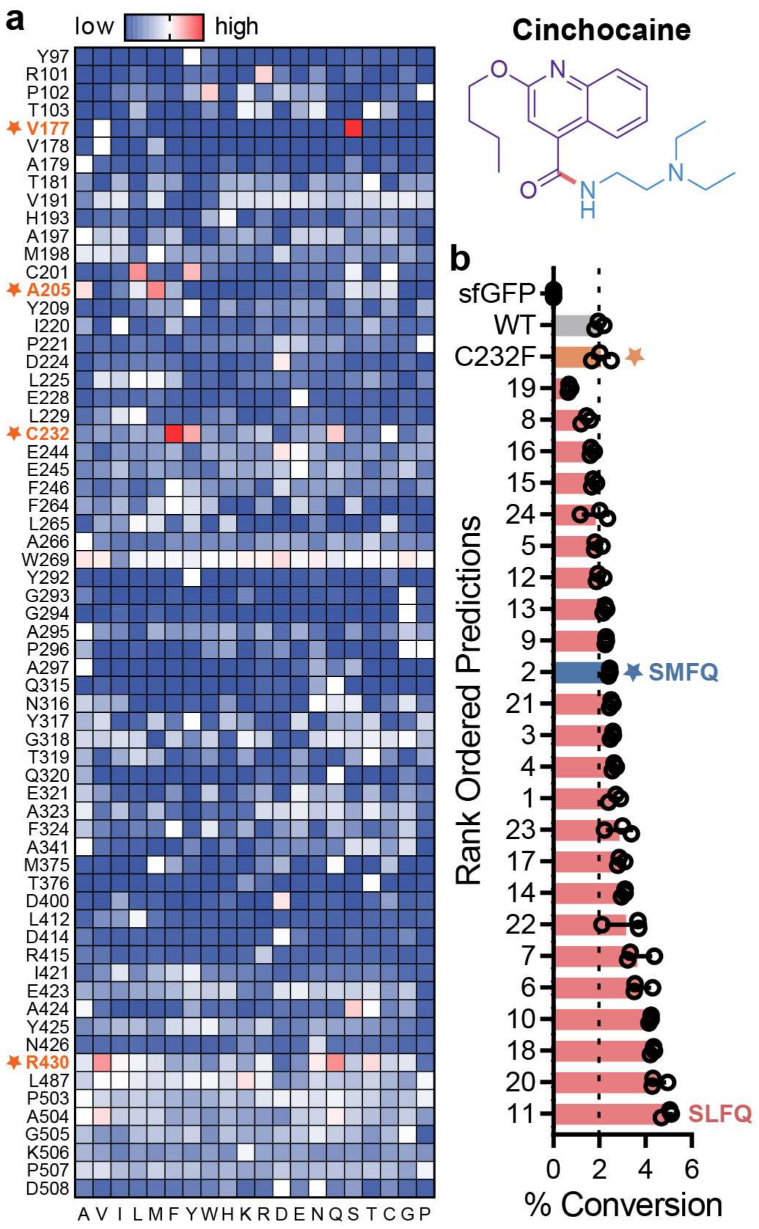


**Figure S16. Hot spot screen and machine-learning guided predictions for the biocatalytic synthesis of cinchocaine. a,** HSS of 64-site library to identify McbA residues that positively impact cinchocaine synthesis (*n* = 1). Yields are normalized to wt-McbA. **b,** Experimental validation of the top 24 ML-predictions, rank ordered by measured activity (*n* = 3). The variants are labeled by their predicted rank. The most active single mutant from the HSS (orange) and a "rational" design from combining the top four single mutations from the HSS (blue) were also included. The specific mutations corresponding to the ranks in this figure can be found in **Table S2**.

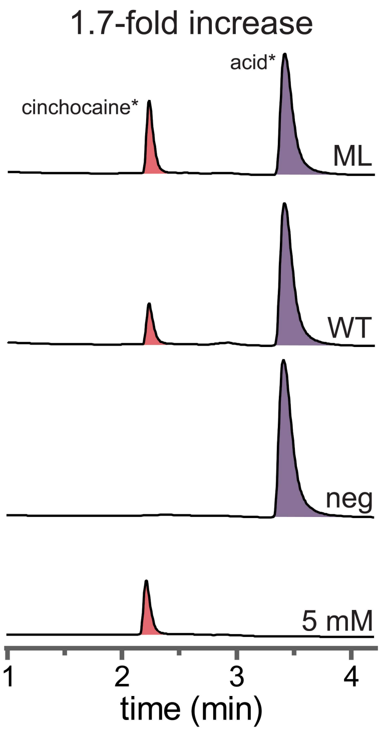


**Figure S17. Machine-learning guided predictions have increased activity over wt-McbA for the synthesis of cinchocaine.** Reversed-phase (RP)-HPLC traces of cinchocaine product (red) and acid substrate (purple) of wt-McbA ('WT') and the machine learning predicted mutant with the highest activity identified in **Fig. S16** ('ML') compared to an authentic standard. HPLC trace data are representative of three independent experiments.

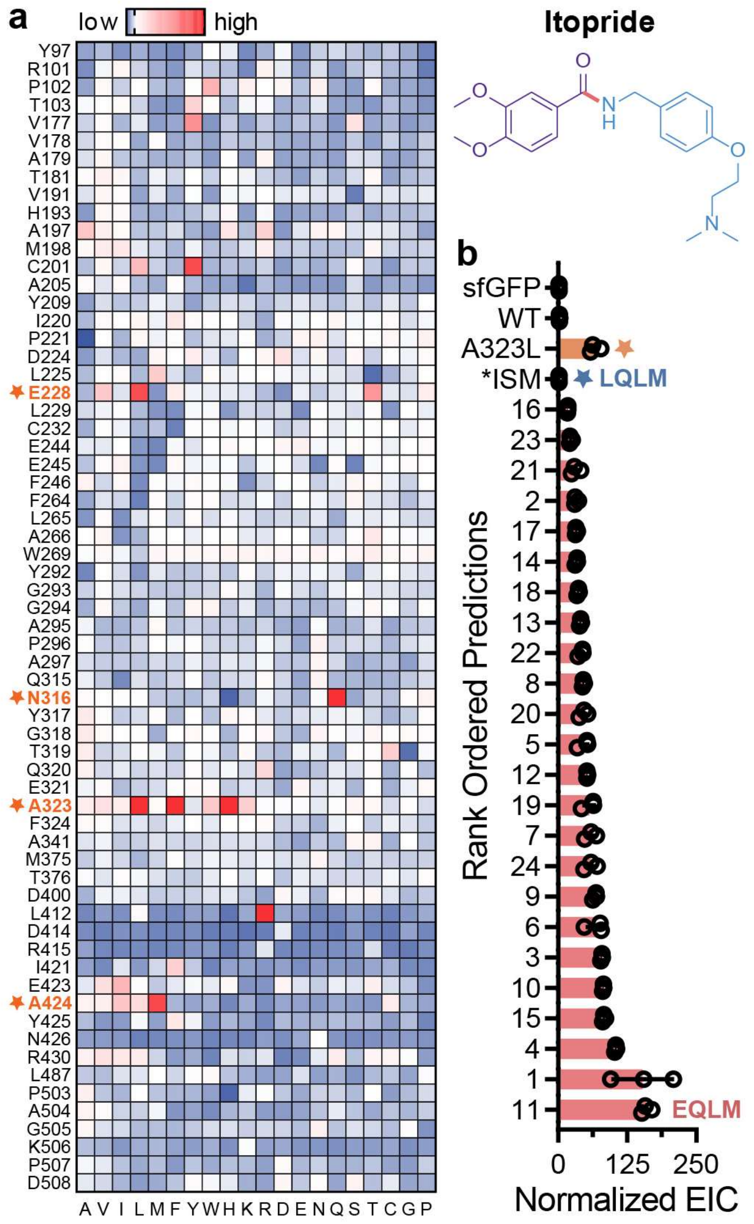


**Figure S18. Hot spot screen and machine-learning guided predictions for the biocatalytic synthesis of itopride. a,** HSS of 64-site library to identify McbA residues that positively impact itopride synthesis ($n$ = 1). Yields are normalized to wt-McbA. **b,** Experimental validation of the top 24 ML-predictions, rank ordered by measured activity ($n$ = 3). The variants are labeled by their predicted rank. As wt-McbA and several mutants produce trace amounts only detectable by MSD, we quantified product with EIC ($m/z$ of 359.2). The most active single mutant from the HSS (orange) and a "rational" design from combining the top four single mutations from the HSS (blue) were also included. The specific mutations corresponding to the ranks in this figure can be found in **Table S3**.

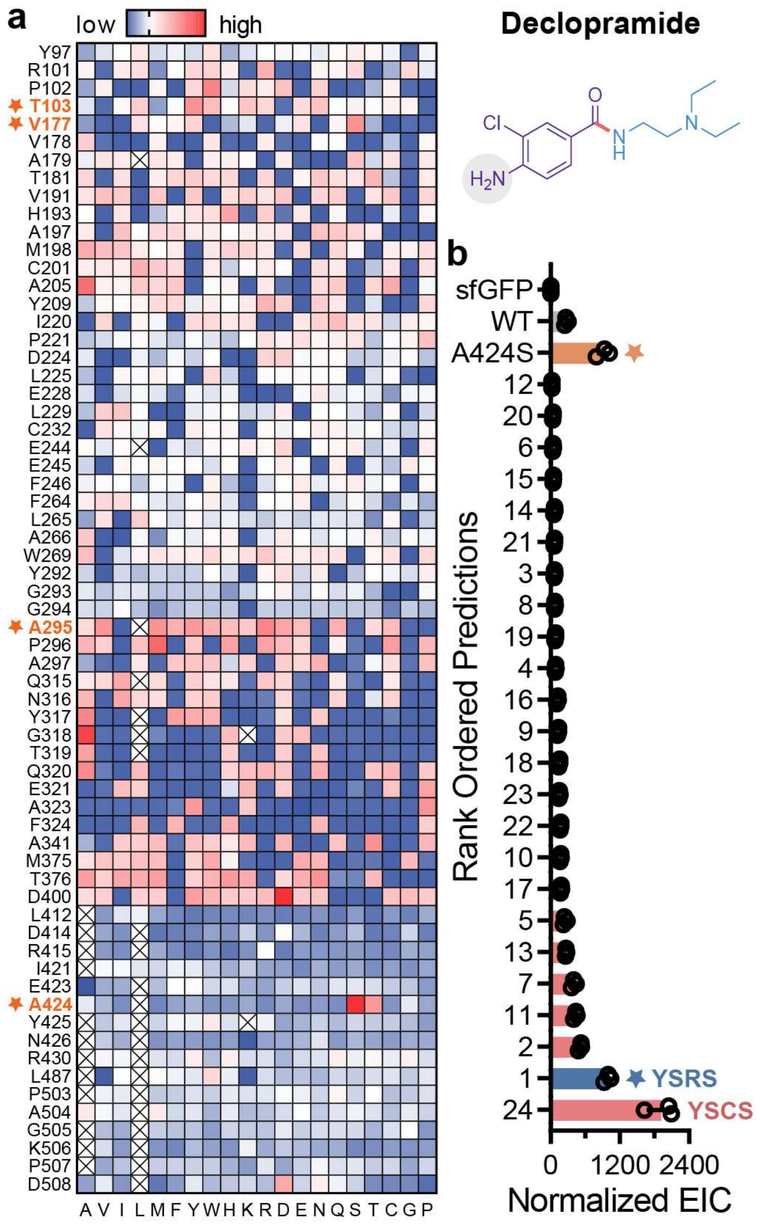


**Figure S19. Hot spot screen and machine-learning guided predictions for the biocatalytic synthesis of declopramide. a,** HSS of 64-site library to identify McbA residues that positively impact declopramide synthesis ($n$ = 1). Yields are normalized to wt-McbA. **b,** Experimental validation of the top 24 ML-predictions, rank ordered by measured activity ($n$ = 3). The variants are labeled by their predicted rank. As wt-McbA and several mutants produce trace amounts only detectable by MSD, we quantified product with EIC ($m/z$ of 270.1). The most active single mutant from the HSS (orange) and a “rational” design from combining the top four single mutations from the HSS (blue) were also included. The specific mutations corresponding to the ranks in this figure can be found in **Table S3.**

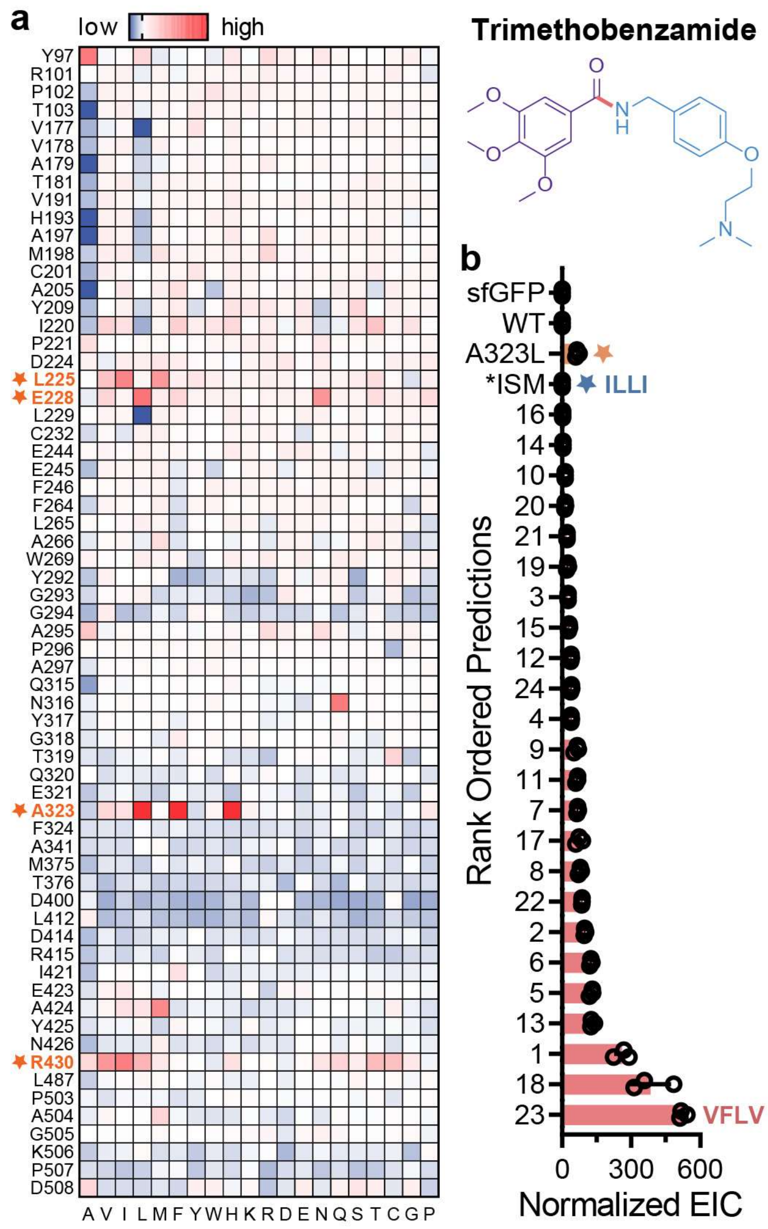


**Figure S20. Hot spot screen and machine-learning guided predictions for the biocatalytic synthesis of trimethobenzamide. a,** HSS of 64-site library to identify McbA residues that positively impact trimethobenzamide synthesis ($n$ = 1). Yields are normalized to wt-McbA. **b,** Experimental validation of the top 24 ML-predictions, rank ordered by measured activity ($n$ = 3). The variants are labeled by their predicted rank. As wt-McbA and several mutants produce trace amounts only detectable by MSD, we quantified product with EIC ($m/z$ of 389.2). The most active single mutant from the HSS (orange) and a "rational" design from combining the top four single mutations from the HSS (blue) were also included. The specific mutations corresponding to the ranks in this figure can be found in **Table S4.**

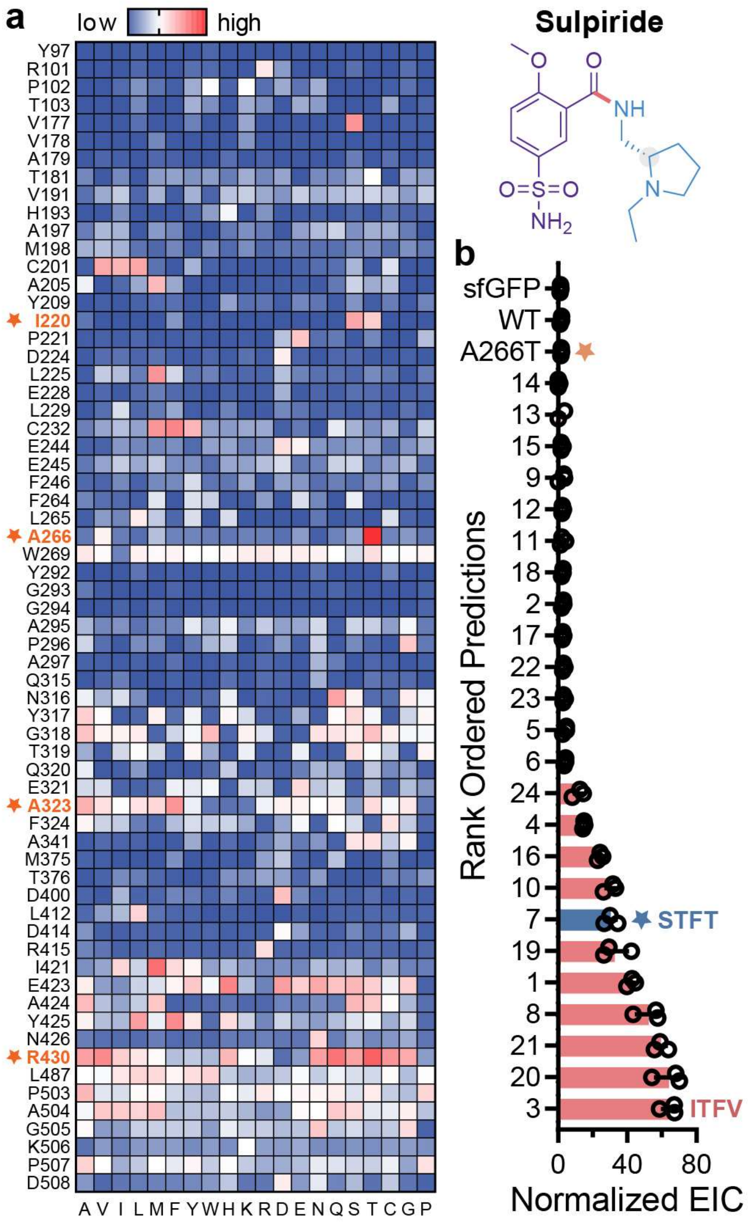


**Figure S21. Hot spot screen and machine-learning guided predictions for the biocatalytic synthesis of S-sulpiride. a,** HSS of 64-site library to identify McbA residues that positively impact S-sulpiride synthesis ($n$ = 1). Yields are normalized to wt-McbA. **b,** Experimental validation of the top 24 ML-predictions, rank ordered by measured activity ($n$ = 3). The variants are labeled by their predicted rank. As wt-McbA and several mutants produce trace amounts only detectable by MSD, we quantified product with EIC ($m/z$ of 342.1). The most active single mutant from the HSS (orange) and a “rational” design from combining the top four single mutations from the HSS (blue) were also included. The specific mutations corresponding to the ranks in this figure can be found in **Table S4**.

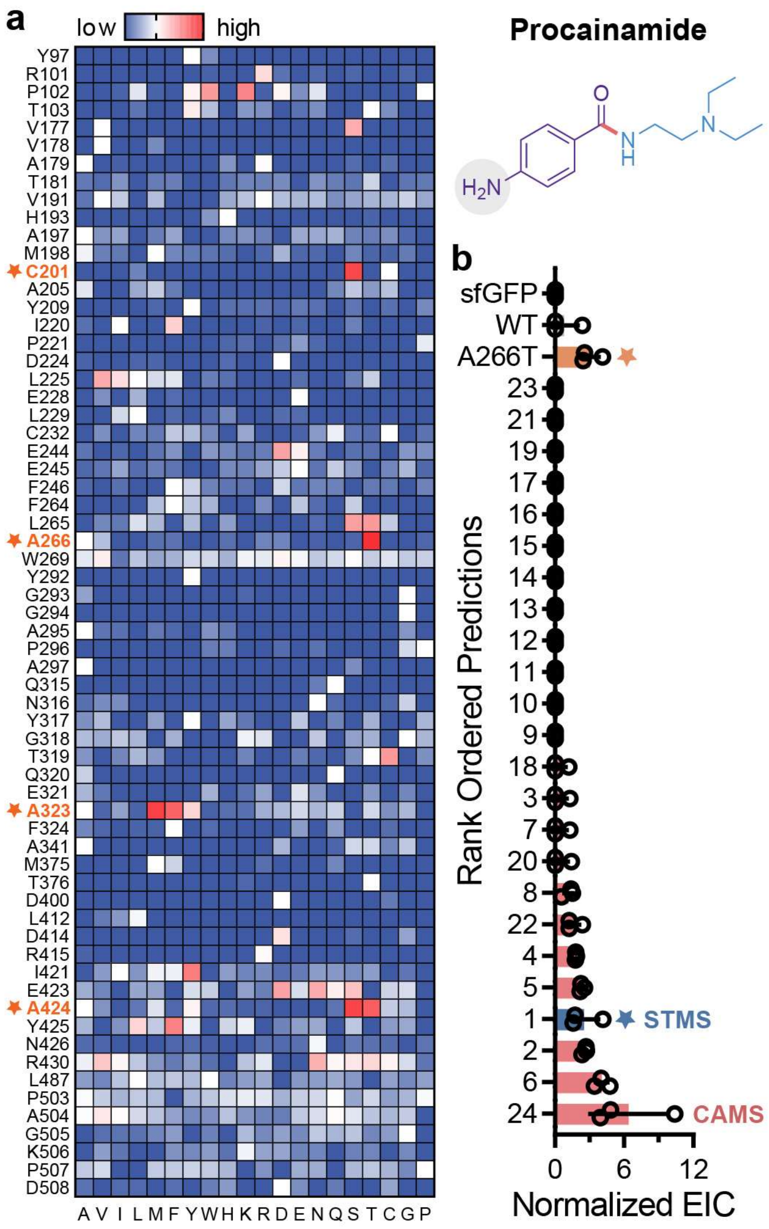


**Figure S22. Hot spot screen and machine-learning guided predictions for the biocatalytic synthesis of procainamide. a,** HSS of 64-site library to identify McbA residues that positively impact procainamide synthesis ($n$ = 1). Yields are normalized to wt-McbA. **b,** Experimental validation of the top 24 ML-predictions, rank ordered by measured activity ($n$ = 3). The variants are labeled by their predicted rank. As wt-McbA and several mutants produce trace amounts only detectable by MSD, we quantified product with EIC ($m/z$ of 236.1). The most active single mutant from the HSS (orange) and a “rational” design from combining the top four single mutations from the HSS (blue) were also included. The specific mutations corresponding to the ranks in this figure can be found in **Table S5.**

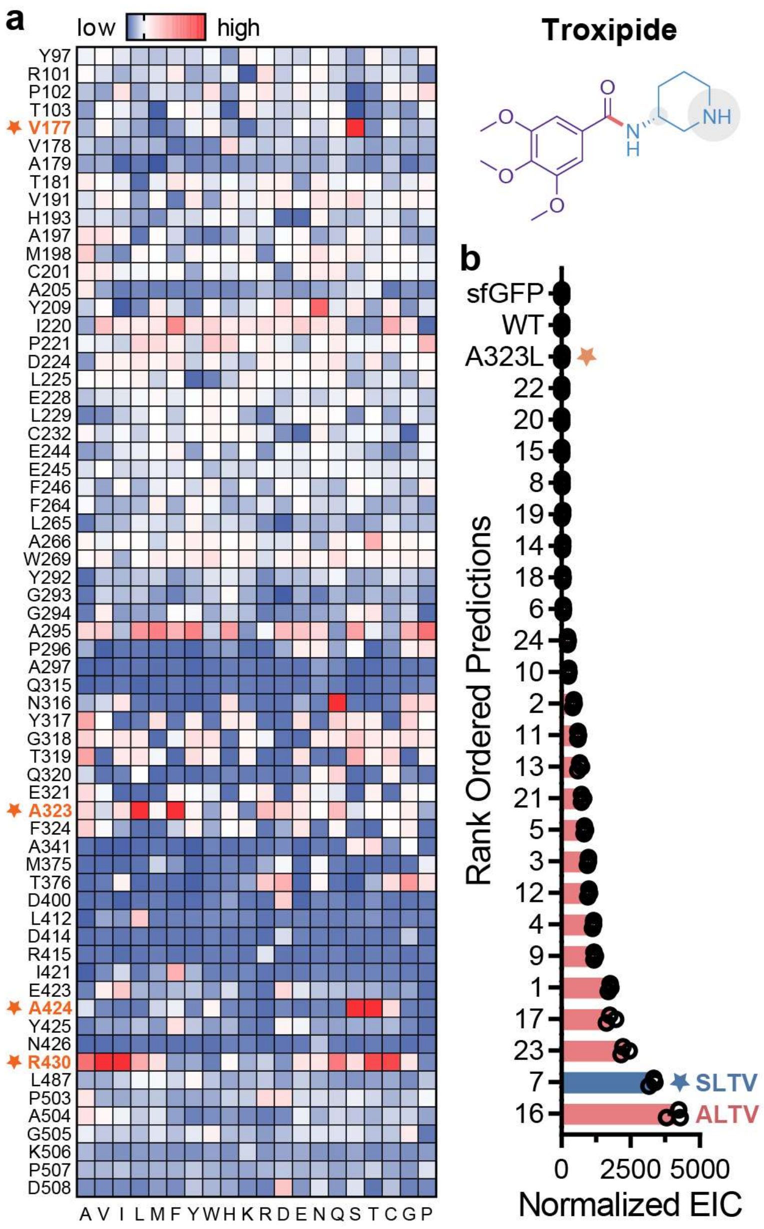


**Figure S23. Hot spot screen and machine-learning guided predictions for the biocatalytic synthesis of troxipide. a,** HSS of 64-site library to identify McbA residues that positively impact troxipide synthesis (*n* = 1). Yields are normalized to wt-McbA. **b,** Experimental validation of the top 24 ML-predictions, rank ordered by measured activity (*n* = 3). The variants are labeled by their predicted rank. As wt-McbA and several mutants produce trace amounts only detectable by MSD, we quantified product with EIC (*m*/z of 295.1). The most active single mutant from the HSS (orange) and a "rational" design from combining the top four single mutations from the HSS (blue) were also included. The specific mutations corresponding to the ranks in this figure can be found in **Table S5**.

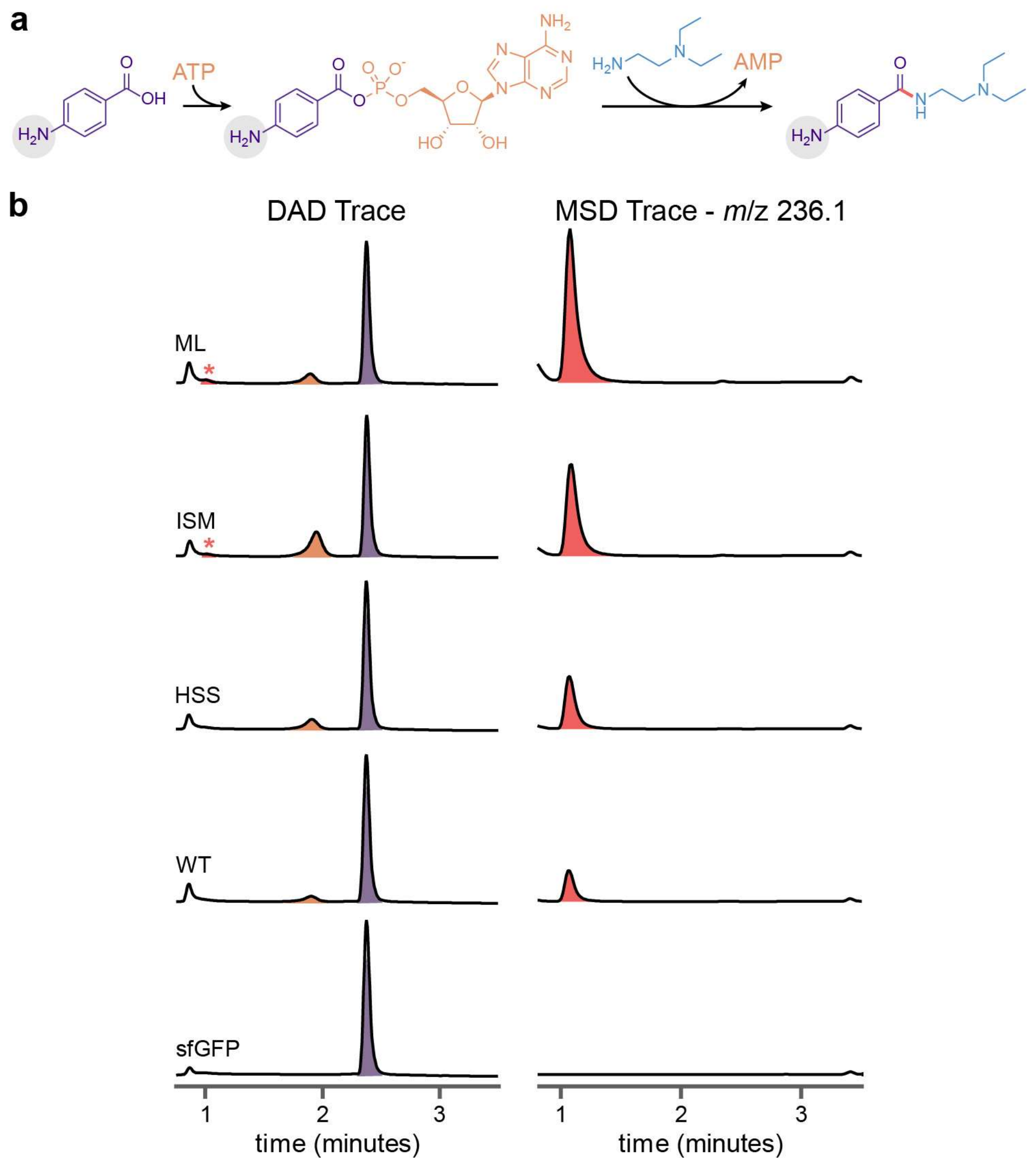


**Figure S24. The two-step reaction mechanism of McbA allows us to probe the impact of mutations on the distinct reaction steps. a,** The coupled reaction mechanism of ATP-dependent amide bond synthetases, such as McbA, is first the activation of an acid to an acyl-adenylate intermediate followed by substitution with an amine nucleophile. In this example, we focus on the synthesis of procainamide (red) using 4-aminobenzoic acid (purple) and *N,N*-diethylethylenediamine (blue). **b,** Since the acyl-adenylate intermediate (orange) is also measurable by DAD (245nm), we can compare how different mutations affect the acid adenylation step vs. the amide bond forming step. The MSD trace (EIC with *m/z* of 236.1) is also included due to the low signal of procainamide under UV absorbance. Key: 'ML' is highest activity ML-predicted mutant, 'ISM' is a 'rational' design from combining the 4 highest activity single mutations from the HSS, and 'HSS' is the highest activity single mutant from the HSS. Data are representative of three independent experiments.

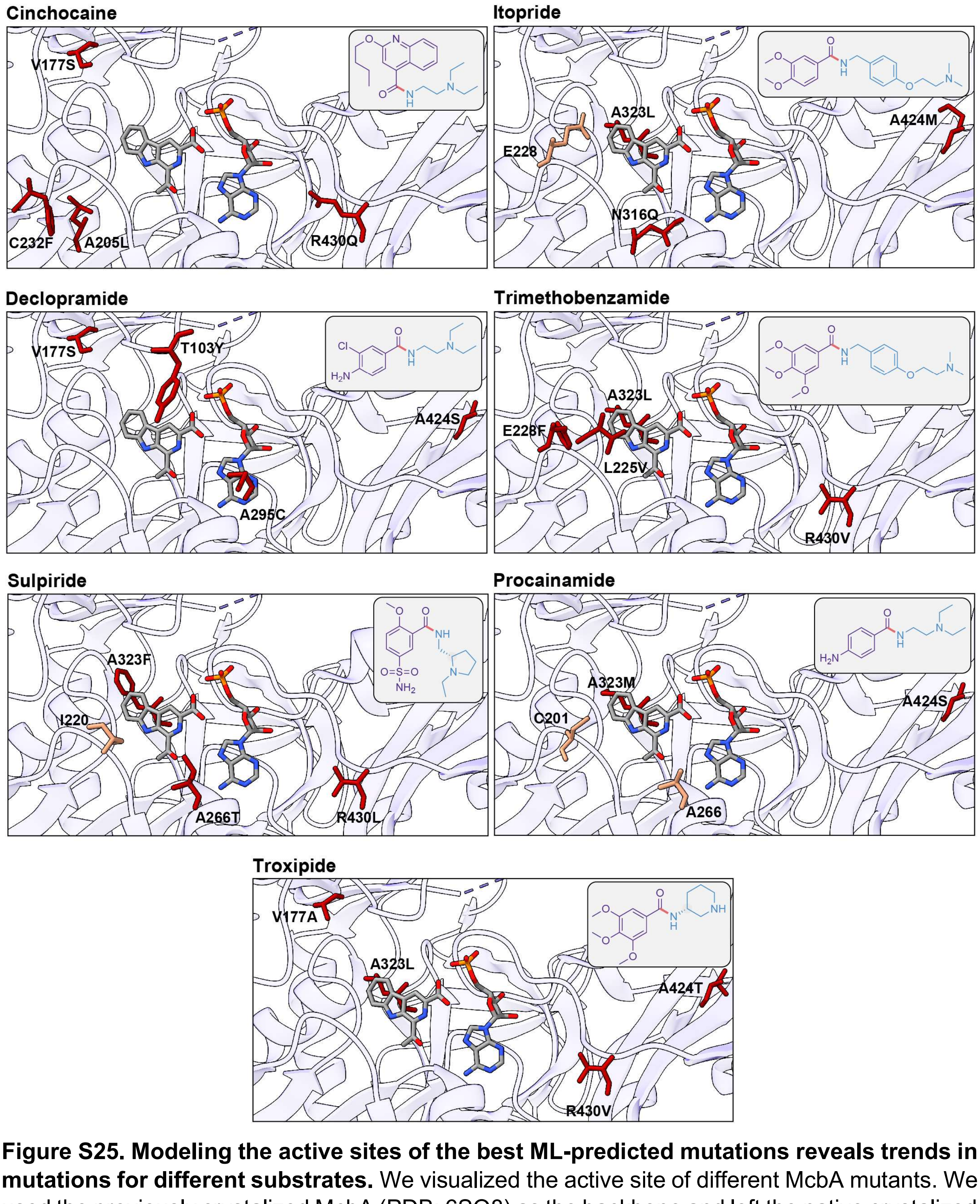


**Figure S25. Modeling the active sites of the best ML-predicted mutations reveals trends in mutations for different substrates.** We visualized the active site of different McbA mutants. We used the previously crystalized McbA (PDB: 6SQ8) as the backbone and left the native crystalized substrates in the active site. Amino acid changes were made at select residues using the rotamer tool with default parameters in ChimeraX.

### Supplementary Tables

**Table S1. Experimental validation of ML-predicted McbA mutants for moclobemide and metoclopramide.** Variant ID corresponds to the four residues mutated for moclobemide (V177, I220, A323, R430) and metoclopramide (V177, T319, A323, A424), in the given order. Variants are rank ordered according to their experimentally measured yield via liquid chromatography (LC) with their predicted rank given. Amino acids colored red correspond to a residue containing the wild type (unmutated) amino acid. An sfGFP negative control is also included. Reported yields are averages and standard deviations of *n* = 3 reactions.

| Moclobemide | | | Metoclopramide | | |
|---|---|---|---|---|---|
| **Variant ID** | **Predicted Rank** | **% Conversion** | **Variant ID** | **Predicted Rank** | **% Conversion** |
| SSFT | 7 | 70.1 ± 0.3 | SCFS | 10 | 60.5 ± 1.1 |
| SSFI | 12 | 67.6 ± 0.7 | SCFT | 2 | 47.8 ± 0.3 |
| SSFV | 1 | 67.6 ± 0.3 | SAFS | 16 | 45.2 ± 0.4 |
| SSFL | 14 | 67.4 ± 0.2 | SSFT | 22 | 44.6 ± 2.3 |
| SSFQ | 5 | 66.3 ± 0.4 | SCLT | 12 | 27.4 ± 0.4 |
| SSLC | 16 | 65.4 ± 0.2 | STFT | 8 | 25.2 ± 1.0 |
| SSFR | 8 | 64.7 ± 0.9 | SAFT | 4 | 22.8 ± 3.5 |
| SSLQ | 11 | 63.2 ± 0.7 | VCFT | 5 | 17.8 ± 0.9 |
| SSLT | 18 | 63.2 ± 0.4 | SLFT | 20 | 13.8 ± 0.7 |
| SSFC | 6 | 63.1 ± 0.2 | SEFT | 19 | 9.07 ± 1.3 |
| SSLV | 2 | 62.6 ± 0.4 | VAFT | 6 | 8.65 ± 1.9 |
| SSLI | 25 | 61.4 ± 0.2 | YSFT | 18 | 6.97 ± 0.0 |
| SSFN | 9 | 60.6 ± 1.0 | YCFS | 9 | 4.57 ± 0.6 |
| SSLN | 22 | 58.3 ± 0.3 | YCFT | 1 | 3.63 ± 0.1 |
| SSLR | 21 | 57.6 ± 0.3 | YCLT | 11 | 2.85 ± 0.5 |
| SAFV | 17 | 56.0 ± 0.7 | KCFT | 24 | 2.17 ± 0.3 |
| STFV | 19 | 53.8 ± 0.1 | YTFT | 7 | 2.04 ± 1.3 |
| SIFQ | 24 | 50.6 ± 0.0 | YAFS | 15 | 0.22 ± 0.1 |
| SIFV | 4 | 50.1 ± 0.3 | YLFT | 21 | 0.09 ± 0.0 |
| SSVV | 3 | 47.1 ± 1.0 | YEFT | 17 | 0.09 ± 0.0 |
| SILV | 10 | 46.8 ± 0.5 | SYFT | 14 | 0.08 ± 0.0 |
| SSVQ | 15 | 39.5 ± 0.6 | YMFT | 25 | 0.08 ± 0.0 |
| SIVV | 13 | 26.8 ± 0.8 | YYFT | 13 | 0.05 ± 0.0 |
| SSVT | 23 | 22.8 ± 1.8 | YKFT | 23 | 0.03 ± 0.0 |
| SSVC | 20 | 20.5 ± 0.8 | YAFT | 3 | 0 ± 0 |
| wt | - | 4.38 ± 1.1 | wt | - | 1.37 ± 0.5 |
| sfGFP | - | 0 ± 0 | sfGFP | - | 0 ± 0 |

**Table S2. Experimental validation of ML-predicted McbA mutants for cinchocaine.** Variant ID corresponds to the four residues mutated for cinchocaine (V177, A205, C232, R430) in the given order. Variants are rank ordered according to their experimentally measured yield via LC with their predicted rank given. Amino acids colored red correspond to a residue containing the wild type (unmutated) amino acid. An sfGFP negative control is also included. Reported yields are averages and standard deviations of $n$ = 3 reactions.

**Cinchocaine**

| Variant ID | Predicted Rank | % Conversion |
|---|---|---|
| SLFQ | 11 | 4.9 ± 0.1 |
| SCFV | 20 | 4.5 ± 0.3 |
| SSFQ | 18 | 4.3 ± 0.0 |
| SAFQ | 10 | 4.2 ± 0.0 |
| SLFV | 6 | 3.7 ± 0.3 |
| SAFV | 7 | 3.6 ± 0.5 |
| SAFT | 22 | 3.1 ± 0.7 |
| SSFV | 14 | 3.0 ± 0.0 |
| SMYQ | 17 | 2.9 ± 0.1 |
| STFV | 23 | 2.8 ± 0.4 |
| SMFV | 1 | 2.6 ± 0.2 |
| SMFN | 4 | 2.6 ± 0.0 |
| SMFT | 3 | 2.5 ± 0.0 |
| SLFT | 21 | 2.5 ± 0.0 |
| SMFQ | 2 | 2.4 ± 0.0 |
| SMFA | 9 | 2.2 ± 0.0 |
| SMYV | 13 | 2.2 ± 0.0 |
| SMFG | 12 | 1.9 ± 0.1 |
| SMFI | 5 | 1.9 ± 0.1 |
| SMFS | 24 | 1.8 ± 0.4 |
| SMFR | 15 | 1.7 ± 0.0 |
| SMFM | 16 | 1.6 ± 0.0 |
| SMFL | 8 | 1.4 ± 0.1 |
| SMQV | 19 | 0.6 ± 0.0 |
| WT | - | 1.9 ± 0.1 |
| sfGFP | - | 0 ± 0 |

**Table S3. Experimental validation of ML-predicted McbA mutants for itopride and declopramide.** Variant ID corresponds to the four residues mutated for itopride (E228, N316, A323, A424) and declopramide (T103, V177, A295, A424) in the given order. Variants are rank ordered according to their experimentally measured extracted ion chromatogram (EIC; *m*/z of 359.2 and 270.1 for itopride and declopramide, respectively) with their predicted rank given. Amino acids colored red correspond to a residue containing the wild type (unmutated) amino acid. An sfGFP negative control is also included. Reported EICs are averages plus or minus standard deviations of $n$ = 3 reactions. EIC values were normalized by dividing EIC by the measured DAD (245 nm) peak area of the acid substrate. Here, we use these values qualitatively to identify the highest activity variant.

| **Itopride** | | | **Declopramide** | | |
|---|---|---|---|---|---|
| **Variant ID** | **Predicted Rank** | **Normalized EIC** | **Variant ID** | **Predicted Rank** | **Normalized EIC** |
| EQLM | 11 | 159.3 ± 7.4 | YSCS | 24 | 1920 ± 209 |
| EQLA | 1 | 152.9 ± 46.3 | YSRS | 1 | 987.2 ± 45.7 |
| LQLG | 4 | 104.3 ± 1.2 | YSMS | 2 | 511.6 ± 23.2 |
| QQLG | 15 | 82.8 ± 2.0 | YSQS | 11 | 431.8 ± 23.7 |
| IQLG | 10 | 81.6 ± 0.7 | WSRS | 7 | 401.1 ± 33.4 |
| LQLA | 3 | 78.6 ± 1.0 | WSMS | 13 | 265.5 ± 6.4 |
| TQLG | 6 | 67.0 ± 14. | YVRS | 5 | 246.8 ± 27.6 |
| QQLA | 9 | 66.6 ± 2.9 | KSRS | 17 | 179.6 ± 9.0 |
| CQLG | 24 | 58.8 ± 9.8 | YVMS | 10 | 171.9 ± 7.7 |
| IQLA | 7 | 58.6 ± 8.6 | WSYS | 22 | 169.6 ± 6.4 |
| VQLA | 19 | 56.4 ± 10. | LSRS | 23 | 152.5 ± 7.3 |
| FQLG | 12 | 52.4 ± 0.6 | YSHS | 18 | 148.0 ± 9.0 |
| TQLA | 5 | 46.7 ± 8.2 | YSFS | 9 | 131.7 ± 8.6 |
| HQLA | 20 | 46.0 ± 6.1 | NSRS | 16 | 118.4 ± 9.4 |
| FQLA | 8 | 45.8 ± 1.4 | YSYS | 4 | 86.2 ± 6.8 |
| YQLG | 22 | 41.5 ± 3.8 | YSES | 19 | 79.6 ± 6.2 |
| AQLA | 13 | 40.3 ± 1.4 | YSWS | 8 | 73.8 ± 7.4 |
| CQLA | 18 | 36.2 ± 1.5 | YSVS | 3 | 66.5 ± 4.0 |
| YQLA | 14 | 33.2 ± 1.9 | WSVS | 21 | 64.7 ± 2.5 |
| SQLA | 17 | 32.8 ± 1.4 | YYRS | 14 | 63.8 ± 6.5 |
| EQLG | 2 | 32.6 ± 2.9 | YVYS | 15 | 46.7 ± 2.4 |
| AQLG | 21 | 31.4 ± 7.1 | YSDS | 6 | 40.9 ± 3.5 |
| PQLG | 23 | 22.4 ± 1.9 | YSKS | 20 | 40.2 ± 4.3 |
| PQLA | 16 | 17.4 ± 0.2 | YVVS | 12 | 23.6 ± 1.3 |
| WT | - | 2.6 ± 0.1 | WT | - | 271.7 ± 22.0 |
| sfGFP | - | 1.5 ± 0.0 | sfGFP | - | 1.9 ± 0.2 |

**Table S4. Experimental validation of ML-predicted McbA mutants for trimethobenzamide and S-sulpiride.** Variant ID corresponds to the four residues mutated for trimethobenzamide (L225, E228, A323, R430) and sulpiride (I220, A266, A323, R430) in the given order. Variants are rank ordered according to their experimentally measured EIC (*m*/z of 389.2 and 342.1 for trimethobenzamide and sulpiride, respectively) with their predicted rank given. Amino acids colored red correspond to a residue containing the wild type (unmutated) amino acid. An sfGFP negative control is also included. Reported EICs are averages plus or minus standard deviations of *n* = 3 reactions. EIC values were normalized by dividing EIC by the measured DAD (245 nm) peak area of the acid substrate. Here, we use these values qualitatively to identify the highest activity variant.

| Trimethobenzamide | | | Sulpiride | | |
|---|---|---|---|---|---|
| **Variant ID** | **Predicted Rank** | **Normalized EIC** | **Variant ID** | **Predicted Rank** | **Normalized EIC** |
| VFLV | 23 | 523.3 ± 12.8 | ITFV | 3 | 64.4 ± 4.0 |
| VILV | 18 | 384.5 ± 72.6 | TTFV | 20 | 64.1 ± 7.1 |
| LELV | 1 | 261.0 ± 27.9 | STFV | 21 | 59.4 ± 3.2 |
| LELI | 13 | 131.2 ± 7.3 | ITFQ | 8 | 52.4 ± 6.4 |
| LELA | 5 | 127.5 ± 6.3 | ITFT | 1 | 42.4 ± 2.0 |
| LELT | 6 | 124.9 ± 3.9 | ITFA | 19 | 32.7 ± 6.8 |
| VELV | 2 | 98.9 ± 2.2 | STFT | 7 | 30.4 ± 3.2 |
| VILR | 22 | 86.4 ± 1.0 | TTFT | 10 | 30.4 ± 2.9 |
| VELT | 8 | 78.2 ± 5.1 | ITMV | 16 | 24.3 ± 1.2 |
| LQLV | 17 | 73.7 ± 11. | ITMT | 4 | 14.8 ± 0.4 |
| VELA | 7 | 67.7 ± 2.2 | STMT | 24 | 11.7 ± 2.6 |
| VELI | 11 | 65.0 ± 3.4 | ITGT | 6 | 4.0 ± 0.3 |
| LILV | 9 | 62.9 ± 8.2 | ITLV | 5 | 4.0 ± 0.9 |
| LELR | 4 | 39.7 ± 1.0 | ITQT | 23 | 3.3 ± 0.5 |
| LALV | 24 | 39.7 ± 2.0 | STGT | 22 | 3.2 ± 0.5 |
| IELV | 12 | 38.7 ± 1.5 | ITAT | 17 | 2.7 ± 0.3 |
| LFLV | 15 | 30.5 ± 1.8 | ITLT | 2 | 2.7 ± 0.5 |
| VELR | 3 | 26.1 ± 0.3 | ITLA | 18 | 2.6 ± 0.4 |
| LILR | 19 | 24.0 ± 2.9 | ITLQ | 11 | 2.5 ± 1.2 |
| TELV | 21 | 21.3 ± 0.4 | TTLT | 12 | 2.5 ± 0.4 |
| LQLR | 20 | 13.7 ± 0.7 | STLT | 9 | 2.2 ± 1.6 |
| IELR | 10 | 13.6 ± 0.4 | ITTT | 15 | 2.1 ± 0.5 |
| SELV | 14 | 4.18 ± 0.0 | ITGV | 13 | 1.8 ± 1.8 |
| SELR | 16 | 1.91 ± 0.0 | ITVT | 14 | 0.3 ± 0.4 |
| WT | - | 1.04 ± 0.0 | WT | - | 1.7 ± 0.3 |
| sfGFP | - | 0.79 ± 0.0 | sfGFP | - | 1.3 ± 0.1 |

**Table S5. Experimental validation of predicted ML-McbA mutants for procainamide and troxipide.** Variant ID corresponds to the four residues mutated for procainamide (C201, A266, A323, A424) and troxipide (V177, A323, A424, R430) in the given order. Variants are rank ordered according to their experimentally measured EIC (*m*/z of 236.1 and 295.1 for procainamide and troxipide, respectively) with their predicted rank given. Amino acids colored red correspond to a residue containing the wild type (unmutated) amino acid. An sfGFP negative control is also included. Reported EICs are averages plus or minus standard deviations of $n$ = 3 reactions. EIC values were normalized by dividing EIC by the measured DAD (245 nm) peak area of the acid substrate. Here, we use these values qualitatively to identify the highest activity variant.

| Procainamide | | |
|---|---|---|
| **Variant ID** | **Predicted Rank** | **Normalized EIC** |
| CAMS | 24 | 6.3 ± 2.8 |
| CTMT | 6 | 4.0 ± 0.5 |
| CTMS | 2 | 2.5 ± 0.1 |
| STMS | 1 | 2.4 ± 1.1 |
| CTMG | 5 | 2.3 ± 0.1 |
| STMT | 4 | 1.8 ± 0.0 |
| STQS | 22 | 1.6 ± 0.5 |
| CTFS | 8 | 1.1 ± 0.4 |
| CTLS | 20 | 0.4 ± 0.6 |
| STFS | 7 | 0.4 ± 0.6 |
| STMG | 3 | 0.4 ± 0.6 |
| SAMS | 18 | 0.3 ± 0.5 |
| STMA | 9 | 0 ± 0 |
| STFG | 10 | 0 ± 0 |
| CTMA | 11 | 0 ± 0 |
| STFT | 12 | 0 ± 0 |
| CTFG | 13 | 0 ± 0 |
| STYS | 14 | 0 ± 0 |
| CTFT | 15 | 0 ± 0 |
| STGS | 16 | 0 ± 0 |
| STLS | 17 | 0 ± 0 |
| CTGS | 19 | 0 ± 0 |
| STAS | 21 | 0 ± 0 |
| CTYS | 23 | 0 ± 0 |
| WT | - | 0.7 ± 1.1 |
| sfGFP | - | 0 ± 0 |

| Troxipide | | |
|---|---|---|
| **Variant ID** | **Predicted Rank** | **Normalized EIC** |
| ALTV | 16 | 4109 ± 221 |
| SLTV | 7 | 3293 ± 91 |
| SLTA | 23 | 2275 ± 127 |
| MLTV | 17 | 1775 ± 137 |
| VLTV | 1 | 1742 ± 36 |
| VLTQ | 9 | 1188 ± 25 |
| VLTT | 4 | 1145 ± 17 |
| SLTR | 12 | 986 ± 27 |
| VLTA | 3 | 963 ± 19 |
| VLTI | 5 | 840 ± 23 |
| VLTL | 21 | 749 ± 34 |
| VLTN | 13 | 658 ± 53 |
| VLTS | 11 | 600 ± 16 |
| VLTR | 2 | 433 ± 17 |
| LLTV | 10 | 259 ± 9 |
| LLTA | 24 | 229 ± 13 |
| RLTV | 6 | 65.4 ± 1.6 |
| VLAV | 18 | 50.9 ± 5.4 |
| LLTR | 14 | 42.0 ± 2.6 |
| RLTT | 19 | 27.8 ± 17.9 |
| RLTR | 8 | 20.2 ± 0.6 |
| VLGV | 15 | 10.5 ± 0.8 |
| RLTA | 20 | 10.3 ± 0.3 |
| VLGR | 22 | 4.9 ± 0.3 |
| WT | - | 2.8 ± 0.0 |
| sfGFP | - | 2.2 ± 0.1 |

**Table S6. Codon-optimized DNA sequences.** Protein-encoding DNA sequences for McbA variants used in this study are shown below. We include wt-McbA, the two qm-McbA variants from our ISM engineering campaigns for moclobemide and metoclopramide, and the best experimentally validated ML-predicted variant for each compound (designated ml-McbA$^{compound}$). The Variant ID corresponding to **Table S1-5** is also included when available. For convenience, the N-terminal CSL-tag (containing a strep purification tag; see Methods) and TEV protease cleavage site on every variant is colored blue and the mutations made relative to wt-McbA are colored red. The amino acid labeling found in this work is based off these sequences (i.e., ATG = M1), which notably slightly deviates from previously published work and the crystal structure of McbA (PDB: 6SQ8) that does not include a CSL-tag.

| Enzyme | Sequence |
|---|---|
| wt-McbA | ATGGAGAAAAAAATCTGGAGCCATCCGCAGTTCGAAAAAGGCGGATCCGG<br>AGAAAACCTGTATTTCCAGGGCGGTTACGCTCGTCGTGTAATGGATGGTAT<br>CGGTGAAGTAGCGGTAACTGGCGCTGGTGGTTCTGTAACTGGTGCGCGTC<br>TGCGCCATCAGGTTCGTCTGCTGGCTCATGCTCTGACCGAAGCGGGTATT<br>CCGCCAGGCCGTGGTGTAGCATGTCTGCATGCTAACACCTGGCGTGCGAT<br>CGCACTGCGTCTGGCTGTTCAGGCGATTGGTTGCCACTATGTTGGTCTGC<br>GTCCTACCGCTGCTGTTACTGAACAGGCACGCGCAATTGCGGCTGCTGAT<br>TCTGCCGCACTGGTTTTCGAACCAAGCGTTGAAGCTCGTGCAGCTGACCT<br>GCTGGAACGTGTTTCTGTGCCGGTTGTGCTGTCTCTGGGTCCGACCTCTC<br>GTGGCCGTGATATCCTGGCAGCTAGCGTTCCGGAAGGTACGCCGCTGCGT<br>TACCGTGAACACCCAGAAGGTATCGCAGTTGTAGCCTTTACTAGCGGCAC<br>CACTGGCACCCCTAAAGGCGTTGCCCACTCCTCTACCGCTATGAGCGCTT<br>GTGTGGATGCTGCGGTTTCCATGTACGGTCGCGGTCCTTGGCGTTTCCTG<br>ATCCCGATCCCTCTGTCTGACCTGGGTGGCGAACTGGCACAGTGTACCCT<br>GGCTACCGGCGGTACCGTTGTGCTGCTGGAAGAGTTCCAACCGGACGCC<br>GTTCTGGAAGCTATCGAACGTGAACGTGCCACTCACGTGTTCCTGGCGCC<br>GAACTGGCTGTACCAGCTGGCTGAACATCCGGCTCTGCCGCGTTCTGATC<br>TGTCTTCTCTGCGTCGCGTTGTTTACGGCGGTGCACCGGCAGTACCATCT<br>CGTGTAGCAGCAGCACGTGAACGTATGGGTGCTGTGCTGATGCAGAACTA<br>CGGCACCCAGGAAGCAGCTTTCATCGCAGCACTGACTCCAGACGATCACG<br>CACGTCGTGAACTGCTGACCGCTGTAGGTCGTCCTCTGCCACACGTTGAG<br>GTGGAAATCCGTGATGACTCTGGTGGTACTCTGCCGCGTGGTGCGGTAGG<br>TGAAGTCTGGGTACGTTCCCCGATGACTATGTCTGGTTACTGGCGTGACC<br>CGGAACGTACGGCTCAGGTTCTGTCTGGTGGTTGGCTGCGTACTGGTGAT<br>GTTGGTACCTTCGATGAGGATGGTCACCTGCATCTGACCGATCGTCTGCA<br>GGACATCATCATCGTTGAAGCATATAACGTCTATTCCCGTCGTGTGGAACA<br>TGTTCTGACCGAACACCCAGATGTTCGCGCAGCTGCGGTTGTTGGCGTAC<br>CAGATCCGGACTCTGGTGAAGCTGTTTGCGCTGCGGTTGTAGTCGCGGAT<br>GGTGCGGATCCTGACCCTGAACACCTGCGTGCTCTGGTTCGTGATCACCT<br>GGGTGATCTGCACGTTCCTCGCCGTGTTGAGTTCGTTCGCTCCATCCCGG<br>TAACTCCTGCCGGCAAACCAGATAAAGTGAAAGTGCGTACCTGGTTCACC<br>GACTAA |
| qm-McbA$_{moclobemide}$ | ATGGAGAAAAAAATCTGGAGCCATCCGCAGTTCGAAAAAGGCGGATCCGG<br>AGAAAACCTGTATTTCCAGGGCGGTTACGCTCGTCGTGTAATGGATGGTAT<br>CGGTGAAGTAGCGGTAACTGGCGCTGGTGGTTCTGTAACTGGTGCGCGTC<br>TGCGCCATCAGGTTCGTCTGCTGGCTCATGCTCTGACCGAAGCGGGTATT<br>CCGCCAGGCCGTGGTGTAGCATGTCTGCATGCTAACACCTGGCGTGCGAT |

| | |
|---|---|
| | CGCACTGCGTCTGGCTGTTCAGGCGATTGGTTGCCACTATGTTGGTCTGC<br>GTCCTACCGCTGCTGTTACTGAACAGGCACGCGCAATTGCGGCTGCTGAT<br>TCTGCCGCACTGGTTTTCGAACCAAGCGTTGAAGCTCGTGCAGCTGACCT<br>GCTGGAACGTGTTTCTGTGCCGGTTGTGCTGTCTCTGGGTCCGACCTCTC<br>GTGGCCGTGATATCCTGGCAGCTAGCGTTCCGGAAGGTACGCCGCTGCGT<br>TACCGTGAACACCCAGAAGGTATCGCAAGCGTAGCCTTTACTAGCGGCAC<br>CACTGGCACCCCTAAAGGCGTTGCCCACTCCTCTACCGCTATGAGCGCTT<br>GTGTGGATGCTGCGGTTTCCATGTACGGTCGCGGTCCTTGGCGTTTCCTG<br>ATCCCGAGCCCTCTGTCTGACCTGGGTGGCGAACTGGCACAGTGTACCCT<br>GGCTACCGGCGGTACCGTTGTGCTGCTGGAAGAGTTCCAACCGGACGCC<br>GTTCTGGAAGCTATCGAACGTGAACGTGCCACTCACGTGTTCCTGGCGCC<br>GAACTGGCTGTACCAGCTGGCTGAACATCCGGCTCTGCCGCGTTCTGATC<br>TGTCTTCTCTGCGTCGCGTTGTTTACGGCGGTGCACCGGCAGTACCATCT<br>CGTGTAGCAGCAGCACGTGAACGTATGGGTGCTGTGCTGATGCAGAACTA<br>CGGCACCCAGGAAGCATTTTTCATCGCAGCACTGACTCCAGACGATCACG<br>CACGTCGTGAACTGCTGACCGCTGTAGGTCGTCCTCTGCCACACGTTGAG<br>GTGGAAATCCGTGATGACTCTGGTGGTACTCTGCCGCGTGGTGCGGTAGG<br>TGAAGTCTGGGTACGTTCCCCGATGACTATGTCTGGTTACTGGCGTGACC<br>CGGAACGTACGGCTCAGGTTCTGTCTGGTGGTTGGCTGCGTACTGGTGAT<br>GTTGGTACCTTCGATGAGGATGGTCACCTGCATCTGACCGATCGTCTGCA<br>GGACATCATCATCGTTGAAGCATATAACGTCTATTCCCTGCGTGTGGAACA<br>TGTTCTGACCGAACACCCAGATGTTCGCGCAGCTGCGGTTGTTGGCGTAC<br>CAGATCCGGACTCTGGTGAAGCTGTTTGCGCTGCGGTTGTAGTCGCGGAT<br>GGTGCGGATCCTGACCCTGAACACCTGCGTGCTCTGGTTCGTGATCACCT<br>GGGTGATCTGCACGTTCCTCGCCGTGTTGAGTTCGTTCGCTCCATCCCGG<br>TAACTCCTGCCGGCAAACCAGATAAAGTGAAAGTGCGTACCTGGTTCACC<br>GACTAA |
| qm-McbA$_{metoclopramide}$ | ATGGAGAAAAAAATCTGGAGCCATCCGCAGTTCGAAAAAGGCGGATCCGG<br>AGAAAACCTGTATTTCCAGGGCGGTTACGCTCGTCGTGTAATGGATGGTAT<br>CGGTGAAGTAGCGGTAACTGGCGCTGGTGGTTCTGTAACTGGTGCGCGTC<br>TGCGCCATCAGGTTCGTCTGCTGGCTCATGCTCTGACCGAAGCGGGTATT<br>CCGCCAGGCCGTGGTGTAGCATGTCTGCATGCTAACACCTGGCGTGCGAT<br>CGCACTGCGTCTGGCTGTTCAGGCGATTGGTTGCCACTATGTTGGTCTGC<br>GTCCTACCGCTGCTGTTACTGAACAGGCACGCGCAATTGCGGCTGCTGAT<br>TCTGCCGCACTGGTTTTCGAACCAAGCGTTGAAGCTCGTGCAGCTGACCT<br>GCTGGAACGTGTTTCTGTGCCGGTTGTGCTGTCTCTGGGTCCGACCTCTC<br>GTGGCCGTGATATCCTGGCAGCTAGCGTTCCGGAAGGTACGCCGCTGCGT<br>TACCGTGAACACCCAGAAGGTATCGCAAGCGTAGCCTTTACTAGCGGCAC<br>CACTGGCACCCCTAAAGGCGTTGCCCACTCCTCTACCGCTATGAGCGCTT<br>GTGTGGATGCTGCGGTTTCCATGTACGGTCGCGGTCCTTGGCGTTTCCTG<br>ATCCCGATCCCTCTGTCTGACCTGGGTGGCGAACTGGCACAGTGTACCCT<br>GGCTACCGGCGGTACCGTTGTGCTGCTGGAAGAGTTCCAACCGGACGCC<br>GTTCTGGAAGCTATCGAACGTGAACGTGCCACTCACGTGTTCCTGGCGCC<br>GAACTGGCTGTACCAGCTGGCTGAACATCCGGCTCTGCCGCGTTCTGATC<br>TGTCTTCTCTGCGTCGCGTTGTTTACGGCGGTGCACCGGCAGTACCATCT<br>CGTGTAGCAGCAGCACGTGAACGTATGGGTGCTGTGCTGATGCAGAACTA<br>CGGCTGCCAGGAAGCATTTTTCATCGCAGCACTGACTCCAGACGATCACG<br>CACGTCGTGAACTGCTGACCGCTGTAGGTCGTCCTCTGCCACACGTTGAG<br>GTGGAAATCCGTGATGACTCTGGTGGTACTCTGCCGCGTGGTGCGGTAGG<br>TGAAGTCTGGGTACGTTCCCCGATGACTATGTCTGGTTACTGGCGTGACC<br>CGGAACGTACGGCTCAGGTTCTGTCTGGTGGTTGGCTGCGTACTGGTGAT |

| | |
|---|---|
| | GTTGGTACCTTCGATGAGGATGGTCACCTGCATCTGACCGATCGTCTGCA<br>GGACATCATCATCGTTGAAACCTATAACGTCTATTCCCGTCGTGTGGAACA<br>TGTTCTGACCGAACACCCAGATGTTCGCGCAGCTGCGGTTGTTGGCGTAC<br>CAGATCCGGACTCTGGTGAAGCTGTTTGCGCTGCGGTTGTAGTCGCGGAT<br>GGTGCGGATCCTGACCCTGAACACCTGCGTGCTCTGGTTCGTGATCACCT<br>GGGTGATCTGCACGTTCCTCGCCGTGTTGAGTTCGTTCGCTCCATCCCGG<br>TAACTCCTGCCGGCAAACCAGATAAAGTGAAAGTGCGTACCTGGTTCACC<br>GACTAA |
| ml-McbA$_{moclobemide}$<br>(SSFT) | ATGGAGAAAAAAATCTGGAGCCATCCGCAGTTCGAAAAAGGCGGATCCGG<br>AGAAAACCTGTATTTCCAGGGCGGTTACGCTCGTCGTGTAATGGATGGTAT<br>CGGTGAAGTAGCGGTAACTGGCGCTGGTGGTTCTGTAACTGGTGCGCGTC<br>TGCGCCATCAGGTTCGTCTGCTGGCTCATGCTCTGACCGAAGCGGGTATT<br>CCGCCAGGCCGTGGTGTAGCATGTCTGCATGCTAACACCTGGCGTGCGAT<br>CGCACTGCGTCTGGCTGTTCAGGCGATTGGTTGCCACTATGTTGGTCTGC<br>GTCCTACCGCTGCTGTTACTGAACAGGCACGCGCAATTGCGGCTGCTGAT<br>TCTGCCGCACTGGTTTTCGAACCAAGCGTTGAAGCTCGTGCAGCTGACCT<br>GCTGGAACGTGTTTCTGTGCCGGTTGTGCTGTCTCTGGGTCCGACCTCTC<br>GTGGCCGTGATATCCTGGCAGCTAGCGTTCCGGAAGGTACGCCGCTGCGT<br>TACCGTGAACACCCAGAAGGTATCGCAAGCGTAGCCTTTACTAGCGGCAC<br>CACTGGCACCCCTAAAGGCGTTGCCCACTCCTCTACCGCTATGAGCGCTT<br>GTGTGGATGCTGCGGTTTCCATGTACGGTCGCGGTCCTTGGCGTTTCCTG<br>ATCCCGAGCCCTCTGTCTGACCTGGGTGGCGAACTGGCACAGTGTACCCT<br>GGCTACCGGCGGTACCGTTGTGCTGCTGGAAGAGTTCCAACCGGACGCC<br>GTTCTGGAAGCTATCGAACGTGAACGTGCCACTCACGTGTTCCTGGCGCC<br>GAACTGGCTGTACCAGCTGGCTGAACATCCGGCTCTGCCGCGTTCTGATC<br>TGTCTTCTCTGCGTCGCGTTGTTTACGGCGGTGCACCGGCAGTACCATCT<br>CGTGTAGCAGCAGCACGTGAACGTATGGGTGCTGTGCTGATGCAGAACTA<br>CGGCACCCAGGAAGCATTTTTCATCGCAGCACTGACTCCAGACGATCACG<br>CACGTCGTGAACTGCTGACCGCTGTAGGTCGTCCTCTGCCACACGTTGAG<br>GTGGAAATCCGTGATGACTCTGGTGGTACTCTGCCGCGTGGTGCGGTAGG<br>TGAAGTCTGGGTACGTTCCCCGATGACTATGTCTGGTTACTGGCGTGACC<br>CGGAACGTACGGCTCAGGTTCTGTCTGGTGGTTGGCTGCGTACTGGTGAT<br>GTTGGTACCTTCGATGAGGATGGTCACCTGCATCTGACCGATCGTCTGCA<br>GGACATCATCATCGTTGAAGCATATAACGTCTATTCCACCCGTGTGGAACA<br>TGTTCTGACCGAACACCCAGATGTTCGCGCAGCTGCGGTTGTTGGCGTAC<br>CAGATCCGGACTCTGGTGAAGCTGTTTGCGCTGCGGTTGTAGTCGCGGAT<br>GGTGCGGATCCTGACCCTGAACACCTGCGTGCTCTGGTTCGTGATCACCT<br>GGGTGATCTGCACGTTCCTCGCCGTGTTGAGTTCGTTCGCTCCATCCCGG<br>TAACTCCTGCCGGCAAACCAGATAAAGTGAAAGTGCGTACCTGGTTCACC<br>GACTAA |
| ml-McbA$_{metoclopramide}$<br>(SCFS) | ATGGAGAAAAAAATCTGGAGCCATCCGCAGTTCGAAAAAGGCGGATCCGG<br>AGAAAACCTGTATTTCCAGGGCGGTTACGCTCGTCGTGTAATGGATGGTAT<br>CGGTGAAGTAGCGGTAACTGGCGCTGGTGGTTCTGTAACTGGTGCGCGTC<br>TGCGCCATCAGGTTCGTCTGCTGGCTCATGCTCTGACCGAAGCGGGTATT<br>CCGCCAGGCCGTGGTGTAGCATGTCTGCATGCTAACACCTGGCGTGCGAT<br>CGCACTGCGTCTGGCTGTTCAGGCGATTGGTTGCCACTATGTTGGTCTGC<br>GTCCTACCGCTGCTGTTACTGAACAGGCACGCGCAATTGCGGCTGCTGAT<br>TCTGCCGCACTGGTTTTCGAACCAAGCGTTGAAGCTCGTGCAGCTGACCT<br>GCTGGAACGTGTTTCTGTGCCGGTTGTGCTGTCTCTGGGTCCGACCTCTC<br>GTGGCCGTGATATCCTGGCAGCTAGCGTTCCGGAAGGTACGCCGCTGCGT<br>TACCGTGAACACCCAGAAGGTATCGCAAGCGTAGCCTTTACTAGCGGCAC<br>CACTGGCACCCCTAAAGGCGTTGCCCACTCCTCTACCGCTATGAGCGCTT |

| | |
|---|---|
| | GTGTGGATGCTGCGGTTTCCATGTACGGTCGCGGTCCTTGGCGTTTCCTG<br>ATCCCGATCCCTCTGTCTGACCTGGGTGGCGAACTGGCACAGTGTACCCT<br>GGCTACCGGCGGTACCGTTGTGCTGCTGGAAGAGTTCCAACCGGACGCC<br>GTTCTGGAAGCTATCGAACGTGAACGTGCCACTCACGTGTTCCTGGCGCC<br>GAACTGGCTGTACCAGCTGGCTGAACATCCGGCTCTGCCGCGTTCTGATC<br>TGTCTTCTCTGCGTCGCGTTGTTTACGGCGGTGCACCGGCAGTACCATCT<br>CGTGTAGCAGCAGCACGTGAACGTATGGGTGCTGTGCTGATGCAGAACTA<br>CGGCTGCCAGGAAGCATTTTTCATCGCAGCACTGACTCCAGACGATCACG<br>CACGTCGTGAACTGCTGACCGCTGTAGGTCGTCCTCTGCCACACGTTGAG<br>GTGGAAATCCGTGATGACTCTGGTGGTACTCTGCCGCGTGGTGCGGTAGG<br>TGAAGTCTGGGTACGTTCCCCGATGACTATGTCTGGTTACTGGCGTGACC<br>CGGAACGTACGGCTCAGGTTCTGTCTGGTGGTTGGCTGCGTACTGGTGAT<br>GTTGGTACCTTCGATGAGGATGGTCACCTGCATCTGACCGATCGTCTGCA<br>GGACATCATCATCGTTGAAAGCTATAACGTCTATTCCCGTCGTGTGGAACA<br>TGTTCTGACCGAACACCCAGATGTTCGCGCAGCTGCGGTTGTTGGCGTAC<br>CAGATCCGGACTCTGGTGAAGCTGTTTGCGCTGCGGTTGTAGTCGCGGAT<br>GGTGCGGATCCTGACCCTGAACACCTGCGTGCTCTGGTTCGTGATCACCT<br>GGGTGATCTGCACGTTCCTCGCCGTGTTGAGTTCGTTCGCTCCATCCCGG<br>TAACTCCTGCCGGCAAACCAGATAAAGTGAAAGTGCGTACCTGGTTCACC<br>GACTAA |
| ml-McbA$_{cinchocaine}$<br>(SLFQ) | ATGGAGAAAAAAATCTGGAGCCATCCGCAGTTCGAAAAAGGCGGATCCGG<br>AGAAAACCTGTATTTCCAGGGCGGTTACGCTCGTCGTGTAATGGATGGTAT<br>CGGTGAAGTAGCGGTAACTGGCGCTGGTGGTTCTGTAACTGGTGCGCGTC<br>TGCGCCATCAGGTTCGTCTGCTGGCTCATGCTCTGACCGAAGCGGGTATT<br>CCGCCAGGCCGTGGTGTAGCATGTCTGCATGCTAACACCTGGCGTGCGAT<br>CGCACTGCGTCTGGCTGTTCAGGCGATTGGTTGCCACTATGTTGGTCTGC<br>GTCCTACCGCTGCTGTTACTGAACAGGCACGCGCAATTGCGGCTGCTGAT<br>TCTGCCGCACTGGTTTTCGAACCAAGCGTTGAAGCTCGTGCAGCTGACCT<br>GCTGGAACGTGTTTCTGTGCCGGTTGTGCTGTCTCTGGGTCCGACCTCTC<br>GTGGCCGTGATATCCTGGCAGCTAGCGTTCCGGAAGGTACGCCGCTGCGT<br>TACCGTGAACACCCAGAAGGTATCGCAAGCGTAGCCTTTACTAGCGGCAC<br>CACTGGCACCCCTAAAGGCGTTGCCCACTCCTCTACCGCTATGAGCGCTT<br>GTGTGGATGCTCTGGTTTCCATGTACGGTCGCGGTCCTTGGCGTTTCCTG<br>ATCCCGATCCCTCTGTCTGACCTGGGTGGCGAACTGGCACAGTTTACCCT<br>GGCTACCGGCGGTACCGTTGTGCTGCTGGAAGAGTTCCAACCGGACGCC<br>GTTCTGGAAGCTATCGAACGTGAACGTGCCACTCACGTGTTCCTGGCGCC<br>GAACTGGCTGTACCAGCTGGCTGAACATCCGGCTCTGCCGCGTTCTGATC<br>TGTCTTCTCTGCGTCGCGTTGTTTACGGCGGTGCACCGGCAGTACCATCT<br>CGTGTAGCAGCAGCACGTGAACGTATGGGTGCTGTGCTGATGCAGAACTA<br>CGGCACCCAGGAAGCAGCTTTCATCGCAGCACTGACTCCAGACGATCACG<br>CACGTCGTGAACTGCTGACCGCTGTAGGTCGTCCTCTGCCACACGTTGAG<br>GTGGAAATCCGTGATGACTCTGGTGGTACTCTGCCGCGTGGTGCGGTAGG<br>TGAAGTCTGGGTACGTTCCCCGATGACTATGTCTGGTTACTGGCGTGACC<br>CGGAACGTACGGCTCAGGTTCTGTCTGGTGGTTGGCTGCGTACTGGTGAT<br>GTTGGTACCTTCGATGAGGATGGTCACCTGCATCTGACCGATCGTCTGCA<br>GGACATCATCATCGTTGAAGCATATAACGTCTATTCCCAGCGTGTGGAACA<br>TGTTCTGACCGAACACCCAGATGTTCGCGCAGCTGCGGTTGTTGGCGTAC<br>CAGATCCGGACTCTGGTGAAGCTGTTTGCGCTGCGGTTGTAGTCGCGGAT<br>GGTGCGGATCCTGACCCTGAACACCTGCGTGCTCTGGTTCGTGATCACCT<br>GGGTGATCTGCACGTTCCTCGCCGTGTTGAGTTCGTTCGCTCCATCCCGG<br>TAACTCCTGCCGGCAAACCAGATAAAGTGAAAGTGCGTACCTGGTTCACC<br>GACTAA |

| | |
|---|---|
| ml-McbA$_{itopride}$<br>(EQLM) | ATGGAGAAAAAAATCTGGAGCCATCCGCAGTTCGAAAAAGGCGGATCCGG<br>AGAAAACCTGTATTTCCAGGGCGGTTACGCTCGTCGTGTAATGGATGGTAT<br>CGGTGAAGTAGCGGTAACTGGCGCTGGTGGTTCTGTAACTGGTGCGCGTC<br>TGCGCCATCAGGTTCGTCTGCTGGCTCATGCTCTGACCGAAGCGGGTATT<br>CCGCCAGGCCGTGGTGTAGCATGTCTGCATGCTAACACCTGGCGTGCGAT<br>CGCACTGCGTCTGGCTGTTCAGGCGATTGGTTGCCACTATGTTGGTCTGC<br>GTCCTACCGCTGCTGTTACTGAACAGGCACGCGCAATTGCGGCTGCTGAT<br>TCTGCCGCACTGGTTTTCGAACCAAGCGTTGAAGCTCGTGCAGCTGACCT<br>GCTGGAACGTGTTTCTGTGCCGGTTGTGCTGTCTCTGGGTCCGACCTCTC<br>GTGGCCGTGATATCCTGGCAGCTAGCGTTCCGGAAGGTACGCCGCTGCGT<br>TACCGTGAACACCCAGAAGGTATCGCAGTTGTAGCCTTTACTAGCGGCAC<br>CACTGGCACCCCTAAAGGCGTTGCCCACTCCTCTACCGCTATGAGCGCTT<br>GTGTGGATGCTGCGGTTTCCATGTACGGTCGCGGTCCTTGGCGTTTCCTG<br>ATCCCGATCCCTCTGTCTGACCTGGGTGGCGAACTGGCACAGTGTACCCT<br>GGCTACCGGCGGTACCGTTGTGCTGCTGGAAGAGTTCCAACCGGACGCC<br>GTTCTGGAAGCTATCGAACGTGAACGTGCCACTCACGTGTTCCTGGCGCC<br>GAACTGGCTGTACCAGCTGGCTGAACATCCGGCTCTGCCGCGTTCTGATC<br>TGTCTTCTCTGCGTCGCGTTGTTTACGGCGGTGCACCGGCAGTACCATCT<br>CGTGTAGCAGCAGCACGTGAACGTATGGGTGCTGTGCTGATGCAGCAGTA<br>CGGCACCCAGGAAGCACTGTTCATCGCAGCACTGACTCCAGACGATCACG<br>CACGTCGTGAACTGCTGACCGCTGTAGGTCGTCCTCTGCCACACGTTGAG<br>GTGGAAATCCGTGATGACTCTGGTGGTACTCTGCCGCGTGGTGCGGTAGG<br>TGAAGTCTGGGTACGTTCCCCGATGACTATGTCTGGTTACTGGCGTGACC<br>CGGAACGTACGGCTCAGGTTCTGTCTGGTGGTTGGCTGCGTACTGGTGAT<br>GTTGGTACCTTCGATGAGGATGGTCACCTGCATCTGACCGATCGTCTGCA<br>GGACATCATCATCGTTGAAATGTATAACGTCTATTCCCGTCGTGTGGAACA<br>TGTTCTGACCGAACACCCAGATGTTCGCGCAGCTGCGGTTGTTGGCGTAC<br>CAGATCCGGACTCTGGTGAAGCTGTTTGCGCTGCGGTTGTAGTCGCGGAT<br>GGTGCGGATCCTGACCCTGAACACCTGCGTGCTCTGGTTCGTGATCACCT<br>GGGTGATCTGCACGTTCCTCGCCGTGTTGAGTTCGTTCGCTCCATCCCGG<br>TAACTCCTGCCGGCAAACCAGATAAAGTGAAAGTGCGTACCTGGTTCACC<br>GACTAA |
| ml-McbA$_{declopramide}$<br>(YSCS) | ATGGAGAAAAAAATCTGGAGCCATCCGCAGTTCGAAAAAGGCGGATCCGG<br>AGAAAACCTGTATTTCCAGGGCGGTTACGCTCGTCGTGTAATGGATGGTAT<br>CGGTGAAGTAGCGGTAACTGGCGCTGGTGGTTCTGTAACTGGTGCGCGTC<br>TGCGCCATCAGGTTCGTCTGCTGGCTCATGCTCTGACCGAAGCGGGTATT<br>CCGCCAGGCCGTGGTGTAGCATGTCTGCATGCTAACACCTGGCGTGCGAT<br>CGCACTGCGTCTGGCTGTTCAGGCGATTGGTTGCCACTATGTTGGTCTGC<br>GTCCTTATGCTGCTGTTACTGAACAGGCACGCGCAATTGCGGCTGCTGATT<br>CTGCCGCACTGGTTTTCGAACCAAGCGTTGAAGCTCGTGCAGCTGACCTG<br>CTGGAACGTGTTTCTGTGCCGGTTGTGCTGTCTCTGGGTCCGACCTCTCG<br>TGGCCGTGATATCCTGGCAGCTAGCGTTCCGGAAGGTACGCCGCTGCGTT<br>ACCGTGAACACCCAGAAGGTATCGCAAGCGTAGCCTTTACTAGCGGCACC<br>ACTGGCACCCCTAAAGGCGTTGCCCACTCCTCTACCGCTATGAGCGCTTG<br>TGTGGATGCTGCGGTTTCCATGTACGGTCGCGGTCCTTGGCGTTTCCTGA<br>TCCCGATCCCTCTGTCTGACCTGGGTGGCGAACTGGCACAGTGTACCCTG<br>GCTACCGGCGGTACCGTTGTGCTGCTGGAAGAGTTCCAACCGGACGCCGT<br>TCTGGAAGCTATCGAACGTGAACGTGCCACTCACGTGTTCCTGGCGCCGA<br>ACTGGCTGTACCAGCTGGCTGAACATCCGGCTCTGCCGCGTTCTGATCTG<br>TCTTCTCTGCGTCGCGTTGTTTACGGCGGTTGCCCGGCAGTACCATCTCGT<br>GTAGCAGCAGCACGTGAACGTATGGGTGCTGTGCTGATGCAGAACTACGG<br>CACCCAGGAAGCAGCTTTCATCGCAGCACTGACTCCAGACGATCACGCAC |

| | |
|---|---|
| | GTCGTGAACTGCTGACCGCTGTAGGTCGTCCTCTGCCACACGTTGAGGTG GAAATCCGTGATGACTCTGGTGGTACTCTGCCGCGTGGTGCGGTAGGTGA AGTCTGGGTACGTTCCCCGATGACTATGTCTGGTTACTGGCGTGACCCGG AACGTACGGCTCAGGTTCTGTCTGGTGGTTGGCTGCGTACTGGTGATGTT GGTACCTTCGATGAGGATGGTCACCTGCATCTGACCGATCGTCTGCAGGA CATCATCATCGTTGAAAGCTATAACGTCTATTCCCGTCGTGTGGAACATGTT CTGACCGAACACCCAGATGTTCGCGCAGCTGCGGTTGTTGGCGTACCAGA TCCGGACTCTGGTGAAGCTGTTTGCGCTGCGGTTGTAGTCGCGGATGGTG CGGATCCTGACCCTGAACACCTGCGTGCTCTGGTTCGTGATCACCTGGGT GATCTGCACGTTCCTCGCCGTGTTGAGTTCGTTCGCTCCATCCCGGTAACT CCTGCCGGCAAACCAGATAAAGTGAAAGTGCGTACCTGGTTCACCGACTA A |
| ml-McbA$_{trimethobenzamide}$ (VFLV) | ATGGAGAAAAAAATCTGGAGCCATCCGCAGTTCGAAAAAGGCGGATCCGG AGAAAACCTGTATTTCCAGGGCGGTTACGCTCGTCGTGTAATGGATGGTAT CGGTGAAGTAGCGGTAACTGGCGCTGGTGGTTCTGTAACTGGTGCGCGTC TGCGCCATCAGGTTCGTCTGCTGGCTCATGCTCTGACCGAAGCGGGTATT CCGCCAGGCCGTGGTGTAGCATGTCTGCATGCTAACACCTGGCGTGCGAT CGCACTGCGTCTGGCTGTTCAGGCGATTGGTTGCCACTATGTTGGTCTGC GTCCTACCGCTGCTGTTACTGAACAGGCACGCGCAATTGCGGCTGCTGAT TCTGCCGCACTGGTTTTCGAACCAAGCGTTGAAGCTCGTGCAGCTGACCT GCTGGAACGTGTTTCTGTGCCGGTTGTGCTGTCTCTGGGTCCGACCTCTC GTGGCCGTGATATCCTGGCAGCTAGCGTTCCGGAAGGTACGCCGCTGCGT TACCGTGAACACCCAGAAGGTATCGCAGTTGTAGCCTTTACTAGCGGCAC CACTGGCACCCCTAAAGGCGTTGCCCACTCCTCTACCGCTATGAGCGCTT GTGTGGATGCTGCGGTTTCCATGTACGGTCGCGGTCCTTGGCGTTTCCTG ATCCCGATCCCTCTGTCTGACGTGGGTGGCTTTCTGGCACAGTGTACCCT GGCTACCGGCGGTACCGTTGTGCTGCTGGAAGAGTTCCAACCGGACGCC GTTCTGGAAGCTATCGAACGTGAACGTGCCACTCACGTGTTCCTGGCGCC GAACTGGCTGTACCAGCTGGCTGAACATCCGGCTCTGCCGCGTTCTGATC TGTCTTCTCTGCGTCGCGTTGTTTACGGCGGTGCACCGGCAGTACCATCT CGTGTAGCAGCAGCACGTGAACGTATGGGTGCTGTGCTGATGCAGAACTA CGGCACCCAGGAAGCACTGTTCATCGCAGCACTGACTCCAGACGATCACG CACGTCGTGAACTGCTGACCGCTGTAGGTCGTCCTCTGCCACACGTTGAG GTGGAAATCCGTGATGACTCTGGTGGTACTCTGCCGCGTGGTGCGGTAGG TGAAGTCTGGGTACGTTCCCCGATGACTATGTCTGGTTACTGGCGTGACC CGGAACGTACGGCTCAGGTTCTGTCTGGTGGTTGGCTGCGTACTGGTGAT GTTGGTACCTTCGATGAGGATGGTCACCTGCATCTGACCGATCGTCTGCA GGACATCATCATCGTTGAAGCATATAACGTCTATTCCGTGCGTGTGGAACA TGTTCTGACCGAACACCCAGATGTTCGCGCAGCTGCGGTTGTTGGCGTAC CAGATCCGGACTCTGGTGAAGCTGTTTGCGCTGCGGTTGTAGTCGCGGAT GGTGCGGATCCTGACCCTGAACACCTGCGTGCTCTGGTTCGTGATCACCT GGGTGATCTGCACGTTCCTCGCCGTGTTGAGTTCGTTCGCTCCATCCCGG TAACTCCTGCCGGCAAACCAGATAAAGTGAAAGTGCGTACCTGGTTCACC GACTAA |
| ml-McbA$_{sulpiride}$ (ITFV) | ATGGAGAAAAAAATCTGGAGCCATCCGCAGTTCGAAAAAGGCGGATCCGG AGAAAACCTGTATTTCCAGGGCGGTTACGCTCGTCGTGTAATGGATGGTAT CGGTGAAGTAGCGGTAACTGGCGCTGGTGGTTCTGTAACTGGTGCGCGTC TGCGCCATCAGGTTCGTCTGCTGGCTCATGCTCTGACCGAAGCGGGTATT CCGCCAGGCCGTGGTGTAGCATGTCTGCATGCTAACACCTGGCGTGCGAT CGCACTGCGTCTGGCTGTTCAGGCGATTGGTTGCCACTATGTTGGTCTGC GTCCTACCGCTGCTGTTACTGAACAGGCACGCGCAATTGCGGCTGCTGAT TCTGCCGCACTGGTTTTCGAACCAAGCGTTGAAGCTCGTGCAGCTGACCT |

| | |
|---|---|
| | GCTGGAACGTGTTTCTGTGCCGGTTGTGCTGTCTCTGGGTCCGACCTCTC<br>GTGGCCGTGATATCCTGGCAGCTAGCGTTCCGGAAGGTACGCCGCTGCGT<br>TACCGTGAACACCCAGAAGGTATCGCAGTTGTAGCCTTTACTAGCGGCAC<br>CACTGGCACCCCTAAAGGCGTTGCCCACTCCTCTACCGCTATGAGCGCTT<br>GTGTGGATGCTGCGGTTTCCATGTACGGTCGCGGTCCTTGGCGTTTCCTG<br>ATCCCGATCCCTCTGTCTGACCTGGGTGGCGAACTGGCACAGTGTACCCT<br>GGCTACCGGCGGTACCGTTGTGCTGCTGGAAGAGTTCCAACCGGACGCC<br>GTTCTGGAAGCTATCGAACGTGAACGTGCCACTCACGTGTTCCTGACCCC<br>GAACTGGCTGTACCAGCTGGCTGAACATCCGGCTCTGCCGCGTTCTGATC<br>TGTCTTCTCTGCGTCGCGTTGTTTACGGCGGTGCACCGGCAGTACCATCT<br>CGTGTAGCAGCAGCACGTGAACGTATGGGTGCTGTGCTGATGCAGAACTA<br>CGGCACCCAGGAAGCATTTTTCATCGCAGCACTGACTCCAGACGATCACG<br>CACGTCGTGAACTGCTGACCGCTGTAGGTCGTCCTCTGCCACACGTTGAG<br>GTGGAAATCCGTGATGACTCTGGTGGTACTCTGCCGCGTGGTGCGGTAGG<br>TGAAGTCTGGGTACGTTCCCCGATGACTATGTCTGGTTACTGGCGTGACC<br>CGGAACGTACGGCTCAGGTTCTGTCTGGTGGTTGGCTGCGTACTGGTGAT<br>GTTGGTACCTTCGATGAGGATGGTCACCTGCATCTGACCGATCGTCTGCA<br>GGACATCATCATCGTTGAAGCATATAACGTCTATTCCGTGCGTGTGGAACA<br>TGTTCTGACCGAACACCCAGATGTTCGCGCAGCTGCGGTTGTTGGCGTAC<br>CAGATCCGGACTCTGGTGAAGCTGTTTGCGCTGCGGTTGTAGTCGCGGAT<br>GGTGCGGATCCTGACCCTGAACACCTGCGTGCTCTGGTTCGTGATCACCT<br>GGGTGATCTGCACGTTCCTCGCCGTGTTGAGTTCGTTCGCTCCATCCCGG<br>TAACTCCTGCCGGCAAACCAGATAAAGTGAAAGTGCGTACCTGGTTCACC<br>GACTAA |
| ml-McbA$_{procainamide}$<br>(CAMS) | ATGGAGAAAAAAATCTGGAGCCATCCGCAGTTCGAAAAAGGCGGATCCGG<br>AGAAAACCTGTATTTCCAGGGCGGTTACGCTCGTCGTGTAATGGATGGTAT<br>CGGTGAAGTAGCGGTAACTGGCGCTGGTGGTTCTGTAACTGGTGCGCGTC<br>TGCGCCATCAGGTTCGTCTGCTGGCTCATGCTCTGACCGAAGCGGGTATT<br>CCGCCAGGCCGTGGTGTAGCATGTCTGCATGCTAACACCTGGCGTGCGAT<br>CGCACTGCGTCTGGCTGTTCAGGCGATTGGTTGCCACTATGTTGGTCTGC<br>GTCCTACCGCTGCTGTTACTGAACAGGCACGCGCAATTGCGGCTGCTGAT<br>TCTGCCGCACTGGTTTTCGAACCAAGCGTTGAAGCTCGTGCAGCTGACCT<br>GCTGGAACGTGTTTCTGTGCCGGTTGTGCTGTCTCTGGGTCCGACCTCTC<br>GTGGCCGTGATATCCTGGCAGCTAGCGTTCCGGAAGGTACGCCGCTGCGT<br>TACCGTGAACACCCAGAAGGTATCGCAGTTGTAGCCTTTACTAGCGGCAC<br>CACTGGCACCCCTAAAGGCGTTGCCCACTCCTCTACCGCTATGAGCGCTT<br>GTGTGGATGCTGCGGTTTCCATGTACGGTCGCGGTCCTTGGCGTTTCCTG<br>ATCCCGATCCCTCTGTCTGACCTGGGTGGCGAACTGGCACAGTGTACCCT<br>GGCTACCGGCGGTACCGTTGTGCTGCTGGAAGAGTTCCAACCGGACGCC<br>GTTCTGGAAGCTATCGAACGTGAACGTGCCACTCACGTGTTCCTGGCGCC<br>GAACTGGCTGTACCAGCTGGCTGAACATCCGGCTCTGCCGCGTTCTGATC<br>TGTCTTCTCTGCGTCGCGTTGTTTACGGCGGTGCACCGGCAGTACCATCT<br>CGTGTAGCAGCAGCACGTGAACGTATGGGTGCTGTGCTGATGCAGAACTA<br>CGGCACCCAGGAAGCAATGTTCATCGCAGCACTGACTCCAGACGATCACG<br>CACGTCGTGAACTGCTGACCGCTGTAGGTCGTCCTCTGCCACACGTTGAG<br>GTGGAAATCCGTGATGACTCTGGTGGTACTCTGCCGCGTGGTGCGGTAGG<br>TGAAGTCTGGGTACGTTCCCCGATGACTATGTCTGGTTACTGGCGTGACC<br>CGGAACGTACGGCTCAGGTTCTGTCTGGTGGTTGGCTGCGTACTGGTGAT<br>GTTGGTACCTTCGATGAGGATGGTCACCTGCATCTGACCGATCGTCTGCA<br>GGACATCATCATCGTTGAAAGCTATAACGTCTATTCCCGTCGTGTGGAACA<br>TGTTCTGACCGAACACCCAGATGTTCGCGCAGCTGCGGTTGTTGGCGTAC<br>CAGATCCGGACTCTGGTGAAGCTGTTTGCGCTGCGGTTGTAGTCGCGGAT |

| | |
|---|---|
| | GGTGCGGATCCTGACCCTGAACACCTGCGTGCTCTGGTTCGTGATCACCT<br>GGGTGATCTGCACGTTCCTCGCCGTGTTGAGTTCGTTCGCTCCATCCCGG<br>TAACTCCTGCCGGCAAACCAGATAAAGTGAAAGTGCGTACCTGGTTCACC<br>GACTAA |
| ml-McbA$_{troxipide}$<br>(ALTV) | ATGGAGAAAAAAATCTGGAGCCATCCGCAGTTCGAAAAAGGCGGATCCGG<br>AGAAAACCTGTATTTCCAGGGCGGTTACGCTCGTCGTGTAATGGATGGTAT<br>CGGTGAAGTAGCGGTAACTGGCGCTGGTGGTTCTGTAACTGGTGCGCGTC<br>TGCGCCATCAGGTTCGTCTGCTGGCTCATGCTCTGACCGAAGCGGGTATT<br>CCGCCAGGCCGTGGTGTAGCATGTCTGCATGCTAACACCTGGCGTGCGAT<br>CGCACTGCGTCTGGCTGTTCAGGCGATTGGTTGCCACTATGTTGGTCTGC<br>GTCCTACCGCTGCTGTTACTGAACAGGCACGCGCAATTGCGGCTGCTGAT<br>TCTGCCGCACTGGTTTTCGAACCAAGCGTTGAAGCTCGTGCAGCTGACCT<br>GCTGGAACGTGTTTCTGTGCCGGTTGTGCTGTCTCTGGGTCCGACCTCTC<br>GTGGCCGTGATATCCTGGCAGCTAGCGTTCCGGAAGGTACGCCGCTGCGT<br>TACCGTGAACACCCAGAAGGTATCGCAGCGGTAGCCTTTACTAGCGGCAC<br>CACTGGCACCCCTAAAGGCGTTGCCCACTCCTCTACCGCTATGAGCGCTT<br>GTGTGGATGCTGCGGTTTCCATGTACGGTCGCGGTCCTTGGCGTTTCCTG<br>ATCCCGATCCCTCTGTCTGACCTGGGTGGCGAACTGGCACAGTGTACCCT<br>GGCTACCGGCGGTACCGTTGTGCTGCTGGAAGAGTTCCAACCGGACGCC<br>GTTCTGGAAGCTATCGAACGTGAACGTGCCACTCACGTGTTCCTGGCGCC<br>GAACTGGCTGTACCAGCTGGCTGAACATCCGGCTCTGCCGCGTTCTGATC<br>TGTCTTCTCTGCGTCGCGTTGTTTACGGCGGTGCACCGGCAGTACCATCT<br>CGTGTAGCAGCAGCACGTGAACGTATGGGTGCTGTGCTGATGCAGAACTA<br>CGGCACCCAGGAAGCACTGTTCATCGCAGCACTGACTCCAGACGATCACG<br>CACGTCGTGAACTGCTGACCGCTGTAGGTCGTCCTCTGCCACACGTTGAG<br>GTGGAAATCCGTGATGACTCTGGTGGTACTCTGCCGCGTGGTGCGGTAGG<br>TGAAGTCTGGGTACGTTCCCCGATGACTATGTCTGGTTACTGGCGTGACC<br>CGGAACGTACGGCTCAGGTTCTGTCTGGTGGTTGGCTGCGTACTGGTGAT<br>GTTGGTACCTTCGATGAGGATGGTCACCTGCATCTGACCGATCGTCTGCA<br>GGACATCATCATCGTTGAAACCTATAACGTCTATTCCGTGCGTGTGGAACA<br>TGTTCTGACCGAACACCCAGATGTTCGCGCAGCTGCGGTTGTTGGCGTAC<br>CAGATCCGGACTCTGGTGAAGCTGTTTGCGCTGCGGTTGTAGTCGCGGAT<br>GGTGCGGATCCTGACCCTGAACACCTGCGTGCTCTGGTTCGTGATCACCT<br>GGGTGATCTGCACGTTCCTCGCCGTGTTGAGTTCGTTCGCTCCATCCCGG<br>TAACTCCTGCCGGCAAACCAGATAAAGTGAAAGTGCGTACCTGGTTCACC<br>GACTAA |

**Table S7. Codon table for designing site saturation mutagenesis primers.** The following codons were used in the forward primers in our cell-free DNA assembly workflow to mutate a desired residue into the corresponding amino acid. While the addition of excess tRNA in CFPS reactions mitigates the negative effects of unoptimized codons, we used the most prevalent codon found in *E. coli* for the compatibility of *in vivo* expression and to prevent the need for re-optimizing the entire sequence.

| Amino Acid | Codon |
|---|---|
| A | GCG |
| R | CGT |
| N | AAC |
| D | GAT |
| C | TGC |
| Q | CAG |
| E | GAA |
| G | GGC |
| H | CAT |
| I | ATT |
| L | CTG |
| K | AAA |
| M | ATG |
| F | TTT |
| P | CCG |
| S | AGC |
| T | ACC |
| W | TGG |
| Y | TAT |
| V | GTG |

**Table S8. Thermocycler parameters for cell-free DNA assembly.** The following thermocycler parameters were consistent throughout this study, with extension time being the only variable changing to compensate for different amplicon lengths. The first step uses touchdown PCR, in which the initial annealing temperature decreases by 1 °C each cycle until a final set temperature is reached.

**PCR 1 parameters:**

| Step | Temp (°C) | Time (min:sec) |
|---|---|---|
| Initial Denaturation | 98 | 3:00 |
| | 98 | 0:30 |
| 6x | 70 (-1 °C/cycle) | 0:30 |
| | 72 | 20s/kbp |
| | 98 | 0:30 |
| 20x | 64 | 0:30 |
| | 72 | 20s/kbp |
| Final Extension | 72 | 10:00 |
| Hold | 12 | ∞ |

**PCR 2 parameters:**

| Step | Temp (°C) | Time (min:sec) |
|---|---|---|
| Initial Denaturation | 98 | 3:00 |
| | 98 | 0:30 |
| 30x | 68 | 0:30 |
| | 72 | 20s/kbp |
| Final Extension | 72 | 10:00 |
| Hold | 12 | ∞ |

**Table S9. Primers for site saturation mutagenesis of wt-McbA.** The base forward (fwd) and reverse (rvs) primers for site saturation mutagenesis of wt-McbA using our cell-free DNA assembly workflow. While there is only a single reverse primer for saturating a single residue, the forward primer carries the desired mutation (so there are subsequently 20 forward primers per residue). We label this flexible position as 'NNN', indicating that all 20 codons found in **Table S7** are inserted here.

| Residue | Direction | Sequence |
|---|---|---|
| Y97 | fwd | CGATTGGTTGCCACNNNGTTGGTCTGCG |
| | rvs | GTGGCAACCAATCGCCTGAACA |
| R101 | fwd | CCACTATGTTGGTCTGNNNCCTACCGC |
| | rvs | CAGACCAACATAGTGGCAACCAATCG |
| P102 | fwd | TGTTGGTCTGCGTNNNACCGCTG |
| | rvs | ACGCAGACCAACATAGTGGCAAC |
| T103 | fwd | GGTCTGCGTCCTNNNGCTGCTG |
| | rvs | AGGACGCAGACCAACATAGTGGC |
| V177 | fwd | CAGAAGGTATCGCANNNGTAGCCTTTACTAGCG |
| | rvs | TGCGATACCTTCTGGGTGTTCACG |
| V178 | fwd | GAAGGTATCGCAGTTNNNGCCTTTACTAGCGG |
| | rvs | AACTGCGATACCTTCTGGGTGTTCA |
| A179 | fwd | GGTATCGCAGTTGTANNNTTTACTAGCGGC |
| | rvs | TACAACTGCGATACCTTCTGGGTGTTC |
| T181 | fwd | GCAGTTGTAGCCTTTNNNAGCGGCACCA |
| | rvs | AAAGGCTACAACTGCGATACCTTCTGG |
| V191 | fwd | CACCCCTAAAGNNNCGGCCCACTC |
| | rvs | GCCTTTAGGGGTGCCAGTGGT |
| H193 | fwd | TAAAGGCGTTGCCNNNTCCTCTACCG |
| | rvs | GGCAACGCCTTTAGGGGTGC |
| A197 | fwd | CCCACTCCTCTACCNNNATGAGCGC |
| | rvs | GGTAGAGGAGTGGGCAACGC |
| M198 | fwd | CTCCTCTACCGCTNNNAGCGCTTGTGTG |
| | rvs | AGCGGTAGAGGAGTGGGCAAC |
| C201 | fwd | GCTATGAGCGCTNNNGTGGATGCTGC |
| | rvs | AGCGCTCATAGCGGTAGAGGAG |
| A205 | fwd | CTTGTGTGGATGCTNNNGTTTCCATGT |
| | rvs | AGCATCCACACAAGCGCTCATAG |
| Y209 | fwd | TGCGGTTTCCATGNNNGGTCGCG |
| | rvs | CATGGAAACCGCAGCATCCACA |
| I220 | fwd | GTTTCCTGATCCCGNNNCCTCTGTCTGAC |
| | rvs | CGGGATCAGGAAACGCCAAGG |
| P221 | fwd | TCCTGATCCCGATCNNNCTGTCTGACC |
| | rvs | GATCGGGATCAGGAAACGCCAAG |
| D224 | fwd | CGATCCCTCTGTCTNNNCTGGGTGG |
| | rvs | AGACAGAGGGATCGGGATCAGGA |
| L225 | fwd | TCCCTCTGTCTGACNNNGGTGGC |
| | rvs | GTCAGACAGAGGGATCGGGATCAG |
| E228 | fwd | GACCTGGGTGNNNCGCTGGCAC |
| | rvs | GCCACCCAGGTCAGACAGAGG |
| L229 | fwd | CTGGGTGGCGAANNNGCACAGTG |

| | | |
|---|---|---|
| | rvs | TTCGCCACCCAGGTCAGACAG |
| C232 | fwd | CGAACTGGCACAGNNNACCCTGGC |
| | rvs | CTGTGCCAGTTCGCCACCC |
| E244 | fwd | CGTTGTGCTGCTGNNNGAGTTCCAACC |
| | rvs | CAGCAGCACAACGGTACCGC |
| E245 | fwd | TGTGCTGCTGGAANNNTTCCAACCG |
| | rvs | TTCCAGCAGCACAACGGTACC |
| F246 | fwd | GCTGCTGGAAGAGNNNCAACCGGAC |
| | rvs | CTCTTCCAGCAGCACAACGGTAC |
| F264 | fwd | GCCACTCACGTGNNNCTGGCG |
| | rvs | CACGTGAGTGGCACGTTCACG |
| L265 | fwd | CCACTCACGTGTTCNNNGCGC |
| | rvs | GAACACGTGAGTGGCACGTTCA |
| A266 | fwd | CTCACGTGTTCCTGNNNCCGAA |
| | rvs | CAGGAACACGTGAGTGGCACG |
| W269 | fwd | TGGCGCCGAACNNNCTGTACC |
| | rvs | GTTCGGCGCCAGGAACACG |
| V292 | fwd | CGTCGCGTTGTTNNNGGCGGTG |
| | rvs | AACAACGCGACGCAGAGAAGAC |
| G293 | fwd | GTCGCGTTGTTTACNNNGGTGC |
| | rvs | GTAAACAACGCGACGCAGAGAAGA |
| G294 | fwd | CGTTGTTTACGNNNCGGCACCG |
| | rvs | GCCGTAAACAACGCGACGC |
| A295 | fwd | TGTTTACGNNNGTGCACCGG |
| | rvs | ACCGCCGTAAACAACGCGAC |
| P296 | fwd | CGGCGGTGCANNNGCAG |
| | rvs | TGCACCGCCGTAAACAACGC |
| A297 | fwd | CGGTGCACCGNNNGTACCATCTC |
| | rvs | CGGTGCACCGCCGTAAACA |
| Q315 | fwd | TGCTGTGCTGATGNNNAACTACGGC |
| | rvs | CATCAGCACAGCACCCATACGTTC |
| N316 | fwd | TGTGCTGATGCAGNNNTACGGCACC |
| | rvs | CTGCATCAGCACAGCACCCATAC |
| Y317 | fwd | TGCTGATGCAGAACNNNGGCACCC |
| | rvs | GTTCTGCATCAGCACAGCACCC |
| G318 | fwd | CTGATGCAGAACTACNNNACCCAGGAA |
| | rvs | GTAGTTCTGCATCAGCACAGCACC |
| T319 | fwd | GCAGAACTACGNNNCGCAGGAAGC |
| | rvs | GCCGTAGTTCTGCATCAGCACAG |
| Q320 | fwd | GAACTACGGCACCNNNGAAGCAGC |
| | rvs | GGTGCCGTAGTTCTGCATCAGC |
| E321 | fwd | TACGGCACCCAGNNNGCAGCTTTCA |
| | rvs | CTGGGTGCCGTAGTTCTGCATC |
| A323 | fwd | CACCCAGGAAGCANNNTTCATCGCAG |
| | rvs | TGCTTCCTGGGTGCCGTAGT |
| F324 | fwd | CCAGGAAGCAGCTNNNATCGCAGCA |
| | rvs | AGCTGCTTCCTGGGTGCC |
| A341 | fwd | GTGAACTGCTGACCNNNGTAGGTCGT |
| | rvs | GGTCAGCAGTTCACGACGTGC |

| | | |
|---|---|---|
| M375 | fwd | GTACGTTCCCCGNNNACTATGTCTGGTTACTGG |
| | rvs | CGGGGAACGTACCCAGACTTCA |
| T376 | fwd | ACGTTCCCCGATGNNNATGTCTGGTTACTGG |
| | rvs | CATCGGGGAACGTACCCAGACT |
| D400 | fwd | GCTGCGTACTGGTNNNGTTGGTACCTTC |
| | rvs | ACCAGTACGCAGCCAACCAC |
| L412 | fwd | TGGTCACCTGCATNNNACCGATCGTC |
| | rvs | ATGCAGGTGACCATCCTCATCGAAG |
| D414 | fwd | CCTGCATCTGACCNNNCGTCTGCAG |
| | rvs | GGTCAGATGCAGGTGACCATCCT |
| R415 | fwd | TGCATCTGACCGATNNNCTGCAGGAC |
| | rvs | ATCGGTCAGATGCAGGTGACCATC |
| I421 | fwd | TGCAGGACATCATCNNNGTTGAAGCATATAACGTC |
| | rvs | GATGATGTCCTGCAGACGATCGGT |
| E423 | fwd | GGACATCATCATCGTTNNNGCATATAACGTCTATTCCCG |
| | rvs | AACGATGATGATGTCCTGCAGACGA |
| A424 | fwd | CATCATCATCGTTGAANNNTATAACGTCTATTCCCGTCG |
| | rvs | TTCAACGATGATGATGTCCTGCAGAC |
| Y425 | fwd | ATCATCGTTGAAGCANNNAACGTCTATTCCCGTCGTG |
| | rvs | TGCTTCAACGATGATGATGTCCTGC |
| N426 | fwd | CATCGTTGAAGCATATNNNGTCTATTCCCGTCGTG |
| | rvs | ATATGCTTCAACGATGATGATGTCCTGC |
| R430 | fwd | GCATATAACGTCTATTCCNNNCGTGTGGAACATG |
| | rvs | GGAATAGACGTTATATGCTTCAACGATGATGATG |
| L487 | fwd | ATCACCTGGGTGATNNNCACGTTCCTC |
| | rvs | ATCACCCAGGTGATCACGAACCAG |
| P503 | fwd | CCATCCCGGTAACTNNNGCCGG |
| | rvs | AGTTACCGGGATGGAGCGAACG |
| A504 | fwd | CCCGGTAACTCCTNNNGGCAAAC |
| | rvs | AGGAGTTACCGGGATGGAGCG |
| G505 | fwd | GGTAACTCCTGCCNNNAAACCAGATAAAGT |
| | rvs | GGCAGGAGTTACCGGGATGGAG |
| K506 | fwd | CTCCTGCCGNNNCGCCAGATAAAGTGAAAGT |
| | rvs | GCCGGCAGGAGTTACCGG |
| P507 | fwd | CCTGCCGGCAAANNNGATAAAGTGAAAGTG |
| | rvs | TTTGCCGGCAGGAGTTACCG |
| D508 | fwd | GCCGGCAAACCANNNAAAGTGAAAGTGCG |
| | rvs | TGGTTTGCCGGCAGGAGTTAC |

**Table S10. Primers for site saturation mutagenesis of muGFP.** The forward (fwd) and reverse (rvs) primers for site saturation mutagenesis of muGFP using our cell-free DNA assembly workflow. We also include the primers used in the optimization of the homologous overlap between the two primers found in **Fig. S3**. The temperature next to the primer for Y66 corresponds to the melting temperature of the overlap.

| Residue | Direction | Sequence |
|---|---|---|
| Y66 | fwd – 27 °C | CCACTCTTACATATGGTGTGTTGTGCTTTAGC |
| | fwd – 33 °C | TACCACTCTTACATATGGTGTGTTGTGCTTTA |
| | fwd – 42 °C | TGTTACCACTCTTACATATGGTGTGTTGTGCT |
| | fwd – 48 °C | CTCTTGTTACCACTCTTACATATGGTGTGTTGTG |
| | fwd – 53 °C | CCTACTCTTGTTACCACTCTTACATATGGTGTGTTG |
| | rvs | TGTAAGAGTGGTAACAAGAGTAGGCC |
| L69 | fwd | CTCTTACATATGGTGTGTTGTGCTTTAGCCG |
| | rvs | CACACCATATGTAAGAGTGGTAACAAGAG |
| Y164 | fwd | AATGGGATCAAAGCATACTTCAAAATCCGC |
| | rvs | TGCTTTGATCCCATTTTTTTGCTTATCAG |
| T203 | fwd | CAATCACTACCTTAGCACACAGTCGGTATT |
| | rvs | GCTAAGGTAGTGATTGTCTGGCAAC |

**Table S11. Primers for amplifying LETs using pJL1 plasmid as a backbone.** These primers are universally used to amplify LETs off pJL1 containing any gene of interest. They add approximately 300 base pairs both upstream and downstream of the coding region to help protect against exonucleases present in the *E. coli* lysate.

| Direction | Sequence |
|---|---|
| LET_fwd | CTGAGATACCTACAGCGTGAGC |
| LET_rvs | CGTCACTCATGGTGATTTCTCACTTG |

**Table S12. BLOSUM50 reduced amino libraries.** We randomly selected four sets of 10 amino acids from groupings derived from the BLOSUM50 matrix. These groupings were used in the retrospective analysis found in **Fig. 4**. To trim the datasets, only single mutations containing one of the following amino acids were included in that training set. Depending on the amino acid at each residue for wt-McbA, this sometimes led to slight changes (+/- 4 data points max) in the total number of samples in the set.

| Group 1 | Group 2 | Group 3 | Group 4 |
|---|---|---|---|
| L | I | M | V |
| C | C | C | C |
| A | A | A | A |
| G | G | G | G |
| S | S | T | T |
| P | P | P | P |
| F | Y | F | W |
| E | D | N | Q |
| K | R | R | K |
| H | H | H | H |

**Table S13. CAS numbers of select compounds used in this study.**

| Product | Acid | Amine | Amide |
|---|---|---|---|
| Moclobemide | 74-11-3 | 2038-03-1 | 71320-77-9 |
| Metoclopramide | 7206-70-4 | 100-36-7 | 364-62-5 |
| Cinchocaine | 10222-61-4 | 100-36-7 | 85-79-0 |
| Itopride | 93-07-2 | 20059-73-8 | 122892-31-3 |
| Declopramide | 2486-71-7 | 100-36-7 | 891-60-0 |
| Trimethobenzamide | 118-41-2 | 20059-73-8 | 554-92-7 |
| (S)-Sulpiride | 22117-85-7 | 22795-99-9 | 15676-16-1 |
| Procainamide | 150-13-0 | 100-36-7 | 51-06-9 |
| Troxipide | 118-41-2 | 334618-23-4 | 30751-05-4 |
